# Comparative analysis of morphological and acoustic correlates of bush-cricket tympanic membranes

**DOI:** 10.1101/2025.11.11.687687

**Authors:** Md Niamul Islam, Lewis B. Holmes, Dominic Rooke, Fabio A. Sarria-S, Fernando Montealegre-Z

## Abstract

Bush-crickets exhibit remarkable auditory diversity, having the ears in the forelegs, where the tympana could be exposed or covered by cuticular flaps known as auditory pinnae. These tympanic membranes receive sound either directly or internally via an acoustic tracheal tube. Despite extensive research on auditory tuning, the relationships between tympanal structure, body size, and acoustic traits remain unclear. In this study, µCT reconstructions with AI-assisted segmentation and phylogenetically informed regressions across 18 bush-cricket species were used to investigate how tympanal surface area and mean cross-sectional thickness relate to body size, carrier frequency, and the presence of auditory pinnae. Tympanal surface area scaled positively with body size, whereas thickness was independent of size. Carrier frequency decreased with increasing body size but showed no direct association with tympanal properties. Among 14 species with auditory pinnae, tympanal dimensions showed no correlation with peak cavity resonance frequency, indicating semi-independent evolution of pinna cavity and tympanal traits. Across species, pinnae did not alter tympanal surface area, although unilateral pinnae were linked to thicker tympana. Within these unilateral species, the pinna-covered tympanum remained consistently larger in area but thinner than the exposed side. Overall, these findings indicate that tympanal evolution reflects a balance of scaling constraints and localised effects of auditory pinnae.

**Summary Statement:** The tympanal morphology in bush-crickets reflects a balance of scaling and pinna effects. Surface area scales with body size and unilateral pinnae generate consistent asymmetries.

## 1. Introduction

Bush-crickets (Orthoptera: Tettigoniidae) possess hearing organs on their forelegs, just below the knee joint on the proximal tibia. Each ear includes a pair of tympanic membranes positioned on opposite sides of the leg, forming an auditory system that shows parallels to certain processing features found in mammalian hearing [1]. However, unlike mammals, sound reaches these membranes through two routes: directly from the surrounding air and indirectly via the acoustic trachea, an internal air-filled tube that connects the thoracic spiracle to the ear through the leg [2]. For both pathways, the tympanic membranes (tympana) convert sound pressure stimuli into mechanical vibrations [3,4], which are transmitted to the *crista-acustica* for frequency analysis and further directional interpretation at the CNS [2,5].

Currently there are more than 8400 recognised species of bush-cricket worldwide, exhibiting a broad range of morphological diversity [6]. Their acoustic communication in the ultrasonic bands for long-distance conspecifics makes them interesting subject for bioacoustics research [7,8]. However, this communication comes at the cost of increased exposure to predation, particularly from bats [9,10]. To counteract this, many bush-cricket species have evolved auditory pinnae (Fig. 1a) as natural ‘bat’. These pinna cavities induce large sound pressure gains at frequencies corresponding to early bat echolocation calls [11,12]. The prevailing view is that the ancestral function of auditory pinnae was protective, shielding the tympanal membranes [13–16]. Interestingly, within the Phaneropterinae, some taxa possess unilateral pinna (Fig. 1b), producing an asymmetry with one pinna-covered and one exposed tympanum [17,18]. In contrast, several groups such as the Phaneropterinae and Phlugidini entirely lack pinnae (Fig. 1c), leaving both tympana exposed [19–23].

**Fig. 1:**
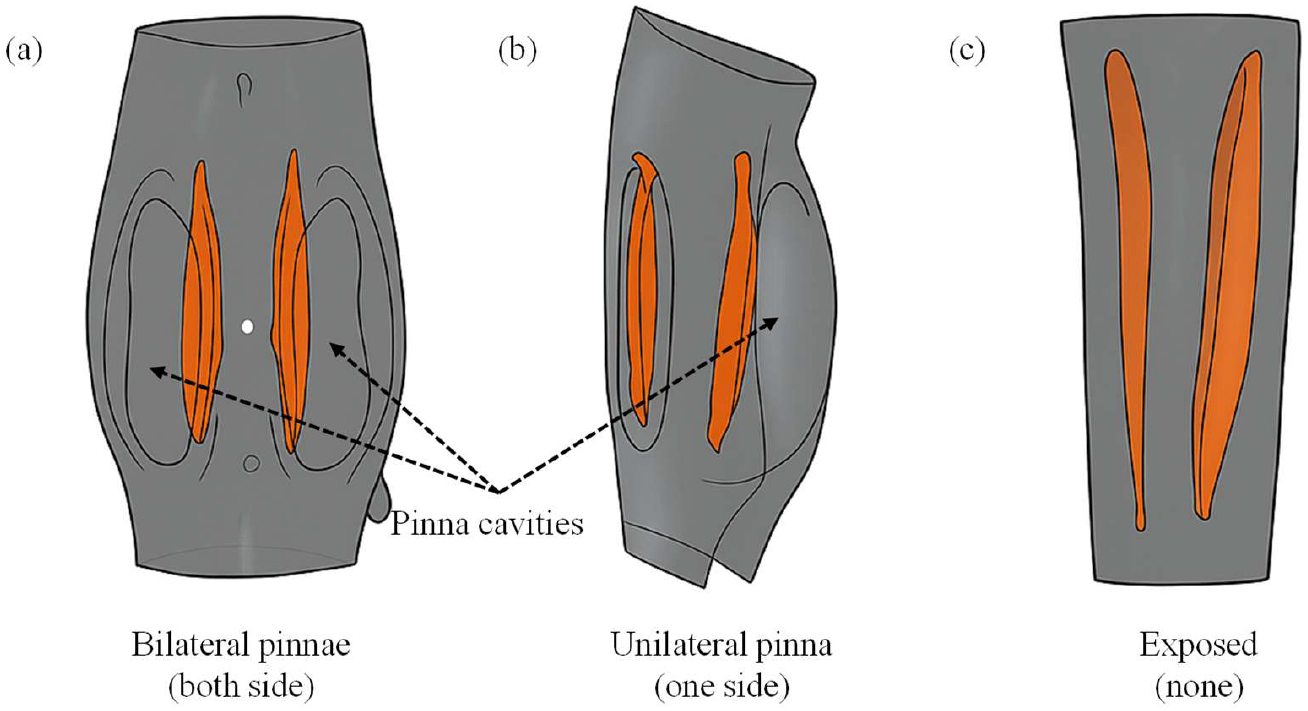
Variations in tympanal pinna configuration among bush-crickets. (a) Bilateral pinnae, with both tympana enclosed within paired pinna cavities. (b) Unilateral pinna, where only one tympanum is covered while the opposite remains exposed. (c) Exposed tympana, where both tympanic membranes are uncovered and lack any pinna structures. Orange regions indicate the tympanic membranes, and dashed arrows highlight the positions of the pinna cavities.

A The key morphological traits of tympana are surface area and thickness. Larger surface areas capture more sound energy, whereas membrane thickness strongly influences frequency response and sensitivity. Thinner regions are often more sensitive to higher frequencies while thicker areas can dampen vibrations, enabling bush-crickets to detect a broad range of acoustic signals [24,25]. Across the Tettigoniids, there is great variation in body size, ear morphology, acoustic function and likely the tympanic membranes [26,27]. This raises the question of how exposed tympana remain undamaged, perhaps they are thicker and less fragile than those shielded by pinnae. Despite extensive research into bush-cricket auditory tuning [2,28–31], a research gap remains in the relationship between tympana structures and other characteristics. This limitation likely stems from the extremely thin and delicate nature of the tympanic membrane, which makes precise morphological characterisation challenging. However, recent advances in high-resolution micro-computed tomography (µCT) and AI-assisted segmentation now allow accurate quantification of tympanal geometry and thickness across species [32].

This study investigates the influence of body size, acoustic properties, ear morphology and geography on the tympana structural properties through comparative analysis of multiple species. Specifically, the effects of pronotum length (as a proxy for body size), carrier frequency, pinna resonance frequency, and pinna presence (exposed, unilateral and bilateral) on tympanal surface area and thickness were assessed. Additionally, asymmetries between pinna-covered and exposed tympana in unilateral species were examined. Considering the critical role of these membranes as the primary receptor of external stimuli, the findings of this study will act as an inspiration and aid towards the development of miniature ultrasound sensitive, frequency-tuned bio-inspired sensors.

## 2. Methods

### 2.1. Specimen preparation

A total of 18 specimens were investigated in this study (listed in Table S1, Supplementary Document 1). Individuals used in this study were either obtained from laboratory colonies or collected from the field. Euthanasia was performed by placing the insects in a freezer at -20 °C for approximately 15 minutes, or they died naturally following experimental or behavioural observations. After euthanasia, the forelegs were carefully removed for subsequent preparation and imaging, including µCT scanning. Each leg was stored in increasing ethanol concentrations, beginning at 20% and increasing in 20% increments every hour up to 100%. Specimens were then submerged for 24 hours in a 1% Lugol’s iodine solution to enhance tissue contrast for µCT imaging. After staining, samples were washed with ethanol 2–3 times to remove excess iodine and subsequently submerged in hexamethyldisilazane (HMDS) overnight. Finally, the legs were desiccated for 24 hours under a fume hood to remove residual liquid and maintain structural integrity for µCT scanning.

### 2.2. µCT imaging and volumetric reconstruction

Specimens were scanned using a SkyScan 1172 µCT scanner (Bruker Corporation, Billerica, MA, USA). Groups of 3–4 legs were placed inside a tube and mounted in the scanner, and the entire tube was imaged as a batch. Scans extended from the tip of the tarsal claws to the top of the knee of the foretibiae. The scan resolution was set to 4032 × 2688 pixels, with a voxel size of 2–3 µm and rotation step of 0.1°. Current and voltage were held constant across all scans at 150 mA and 40 kV, respectively. Raw projection data were reconstructed into orthogonal slices using nRecon (v.1.6.9.18, Bruker Corporation, Billerica, MA, USA), with identical reconstruction parameters applied to each scan. Volumetric reconstructions were generated in Dragonfly 3D software (Object Research Systems, Montreal, Canada).

### 2.3. 3D printing and resonance acoustic testing

Stereolithography (SLA) was used to fabricate physical models of the bush-cricket outer ear with pinna cavities, scaled 10–15× from the original anatomy. Printing was performed on an Elegoo Saturn printer (Elegoo, Shenzhen, China) using ChituBox slicer software. Models were produced in Acrylonitrile Butadiene Styrene (ABS; ABS-Like 2.0 grey resin) with a layer height of 0.06 mm. The bottom layers were exposed for 30 s (first five layers), followed by 7.5 s for subsequent layers. Printed parts were rinsed in 100% isopropyl alcohol and cured in a UV curing station (GeeeTech, Shenzhen, China) for 1 hour.

The pinna cavities of the printed specimen were drilled with 2 mm radius drill bit to make the cavity accessible for the probe microphone. A 25 mm tipped B&K Type 4182 probe microphone (Brüel & Kjær, Nærum, Denmark) with a 1 × 25 mm (0.99″) probe tube length and 1.24 mm (0.05″) interior diameter, calibrated using a B&K Type 4237 sound pressure calibrator was entered through the cavity hole. The ear moved on the microphone using an electronic micromanipulator (TR10/MP-245, Sutter Instrument, Novato, California, USA), to a position 1 cm from the back of the cavity. Received signals were amplified using a B&K 1708 conditioning amplifier (Brüel & Kjær, Nærum, Denmark), acquired using a PSV-500 internal data acquisition board at a sampling frequency of 512 kHz.

Avisoft ultrasonic speaker Vifa with SPEAKON connector (Avisoft-UltraSoundGate; Glienicke/Nordbahn, Germany), positioned 30 cm from the samples and secured on a magnetic stand, was used to deliver acoustic stimuli. The loudspeaker was connected to a portable ultrasonic power amplifier (Avisoft-UltraSoundGate; Glienicke/Nordbahn, Germany) to ensure controlled sound delivery. The scanning vibrometer, loudspeaker, and microphone were connected to a PSV-500 internal data acquisition board (Polytec PSV-500 vibrometer front-head; Waldbronn, Germany), which interfaced with a computer running Polytec 10.1.1 software for system control and data analysis. To match the scale of the printed models, a broadband stimulus consisting of periodic chirps with a simulated frequency range of 2–20 kHz was delivered by the loudspeaker which represented the natural range of 20–200 kHz for 10 times scaled models. To ensure an inital flat stimulus intensity at all frequencies, the amplitude of the broadband signal was mathematically corrected within the software, providing a uniform sound pressure level of 40 dB across all frequencies. The data were post-processed and analysed using Polytec 10.1.1 software and the peak pinna resonance frequency for each species is listed in Table S2 of Supplementary Document 1.

### 2.4. Morphometric measurement and analyses

Tympanic membranes were segmented in Dragonfly 3D software using deep segmentation (AI-assisted) based on a U-net architecture, followed by manual refinement to ensure accuracy [32]. Tympanal surface area was calculated directly from the segmented 3D models, and membrane thickness was quantified as the mean cross-sectional thickness distribution of each tympanum. Summarised mean values are presented in Table S1 (Supplementary Document 1), while full thickness distribution data, along with segmentation and calculation outputs for each of the 18 species, are provided in Supplementary Document 2. For unilateral pinna species, both the exposed and pinna-covered tympana properties were measured separately to allow within-species comparisons, and the results are summarised in Table S3 of Supplementary Document 1.

All morphometric variables were ln-transformed prior to analysis. Pronotum length was used as a proxy for body size. Relationships between tympanal properties, body size, carrier frequency, and pinna resonance frequency were tested using ordinary least squares (OLS) and phylogenetic generalised least squares (PGLS) regressions. PGLS models incorporated Pagel’s λ to estimate phylogenetic signal. Analyses were performed in R (v. 2025.05.1+513), using the packages *ape, caper*, and *nlme*. To control for phylogeny, the phylogenetic tree for the 18 species studied was reconstructed based on latest genetic data and subfamily [33] which is shown in Fig S1 of Supplementary Document 1. For all regression models, λ indicates the phylogenetic signal, R^2^ the coefficient of determination, AICc the corrected Akaike information criterion, B ± SE the regression slope ± standard error, t the test statistic, and p the probability value.

## 3. Results

### 3.1. Body size, carrier frequency, and tympanal morphology

The relationships between body size, carrier frequency, and tympanal membrane morphology are presented in Fig. 1, while the corresponding statistical results are summarised in Table 1. Together, these analyses illustrate how variation in pronotum length, used here as a proxy for body size, relates to acoustic traits and tympanal properties across the 18 bush-cricket species examined.

**Table 1:** Phylogenetically controlled and non-phylogenetically controlled regression analyses of bush-cricket acoustic and morphological traits. Results of phylogenetic generalised least squares (PGLS; blue values) and ordinary least squares (OLS; red values) models examining relationships between carrier frequency, pronotum length (proxy for body size), and tympanal membrane properties (surface area and thickness) across 18 bush-cricket species.

| Parameter | Phylogenetically controlled |  |  |  |  |  | Not phylogenetically controlled |  |  |
| --- | --- | --- | --- | --- | --- | --- | --- | --- | --- |
| | $\lambda$ | $R^2$ | $AICc$ | $B + SE$ | $t$ | $p$ | $B + SE$ | $t$ | $p$ |
| Carrier frequency ~<br>Pronotum length | 0.203 | 0.212 | 41.49 | $-0.995 \pm 0.454$ | -2.191 | 0.044 | $-0.994 \pm 0.479$ | -2.074 | 0.055 |
| Tympana properties ~ pronotum length \| carrier frequency |  |  |  |  |  |  |  |  |  |
| Tympana surface<br>area | 0.242 | 0.495 | 25.72 | $1.179 \pm 0.308$ | 3.831 | 0.002 | $1.145 \pm 0.325$ | 3.519 | 0.003 |
| Tympana thickness | 0.126 | 0.199 | 20.43 | $0.031 \pm 0.270$ | 0.114 | 0.911 | $-0.017 \pm 0.276$ | -0.061 | 0.952 |
| Tympana properties ~ carrier frequency \| pronotum length |  |  |  |  |  |  |  |  |  |
| Tympana surface<br>area | 0.242 | 0.495 | 25.72 | $0.086 \pm 0.149$ | 0.576 | 0.573 | $0.026 \pm 0.151$ | 0.173 | 0.865 |
| Tympana thickness | 0.126 | 0.199 | 20.43 | $-0.184 \pm 0.129$ | -1.431 | 0.173 | $-0.224 \pm 0.128$ | -1.753 | 0.100 |

Carrier frequency was negatively related to body size (pronotum length), with this relationship significant under phylogenetic correction (PGLS: *B* = –0.995 ± 0.454, *t* = –2.191, *p* = 0.044; Fig. 2a; Table 1). When controlling for carrier frequency, tympanal membrane surface area was positively related to body size (PGLS: *B* = 1.179 ± 0.308, *t* = 3.831, *p* = 0.002; Fig. 2; Table 1), whereas tympanal membrane thickness showed no significant relationship with body size (PGLS: *p* = 0.911; Fig. 2d; Table 1). When controlling for body size, tympanal membrane surface area was not significantly related to carrier frequency (PGLS: *p* = 0.573; Fig. 2c; Table 1). Tympanal membrane thickness was negatively related to carrier frequency (PGLS: *B* = – 0.184 ± 0.129, *t* = –1.431, *p* = 0.173; Fig. 2e; Table 1), but this trend was not statistically significant. Phylogenetic signal was low for carrier frequency (λ = 0.20), moderate for tympanal membrane surface area (λ = 0.24), and low for tympanal membrane thickness (λ = 0.13).

**Fig. 2:**
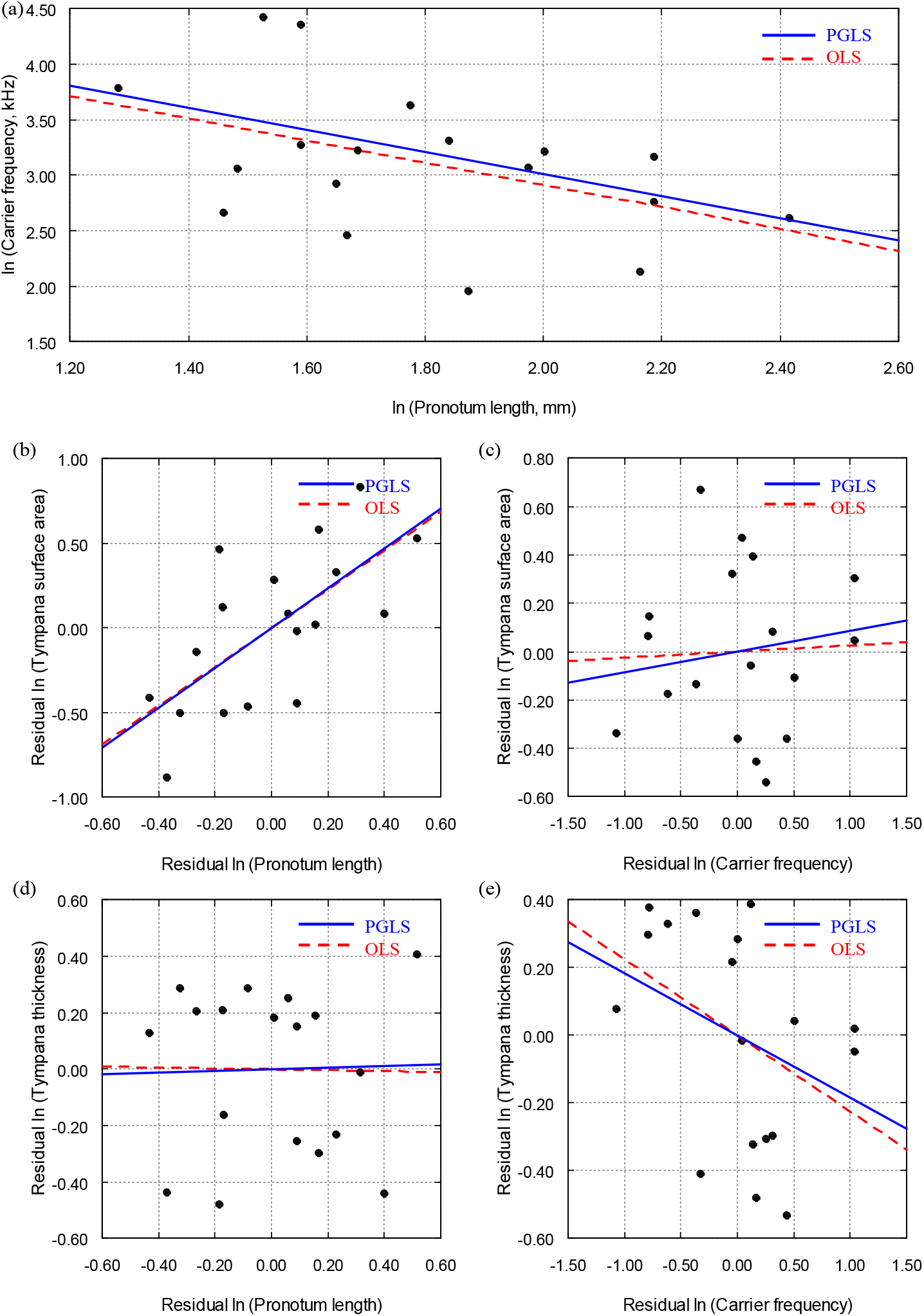
Comparative relationships between body size, carrier frequency, and tympanal morphology in 18 species of bush bush-crickets. (a) Phylogenetic generalised least squares (PGLS, solid blue line) and ordinary least squares (OLS, dashed red line) regressions of ln-transformed carrier frequency (kHz) against ln-transformed pronotum length (mm; used here as a proxy for body size). (b–c) Standardised residuals of ln-transformed pronotum length and carrier frequency plotted against residual ln-transformed tympanal membrane surface area. (d–e) Standardised residuals of pronotum length and carrier frequency plotted against residual ln-transformed tympanal membrane thickness.

### 3.2. Tympanal morphology and pinna resonance frequency

Tympanal membrane morphology was examined in relation to peak pinna resonance frequency while controlling for body size and carrier frequency in the 14 species with pinna cavities (Fig. 3; Table 2). Neither tympanal membrane surface area nor thickness showed significant associations with resonance frequency (surface area: PGLS: p = 0.219; OLS: p = 0.219; Fig. 3a; Table 2; thickness: PGLS: p = 0.990; OLS: p = 0.990; Fig. 3b; Table 2). Phylogenetic signal was absent in both models (λ = 0.00).

**Fig. 3:**
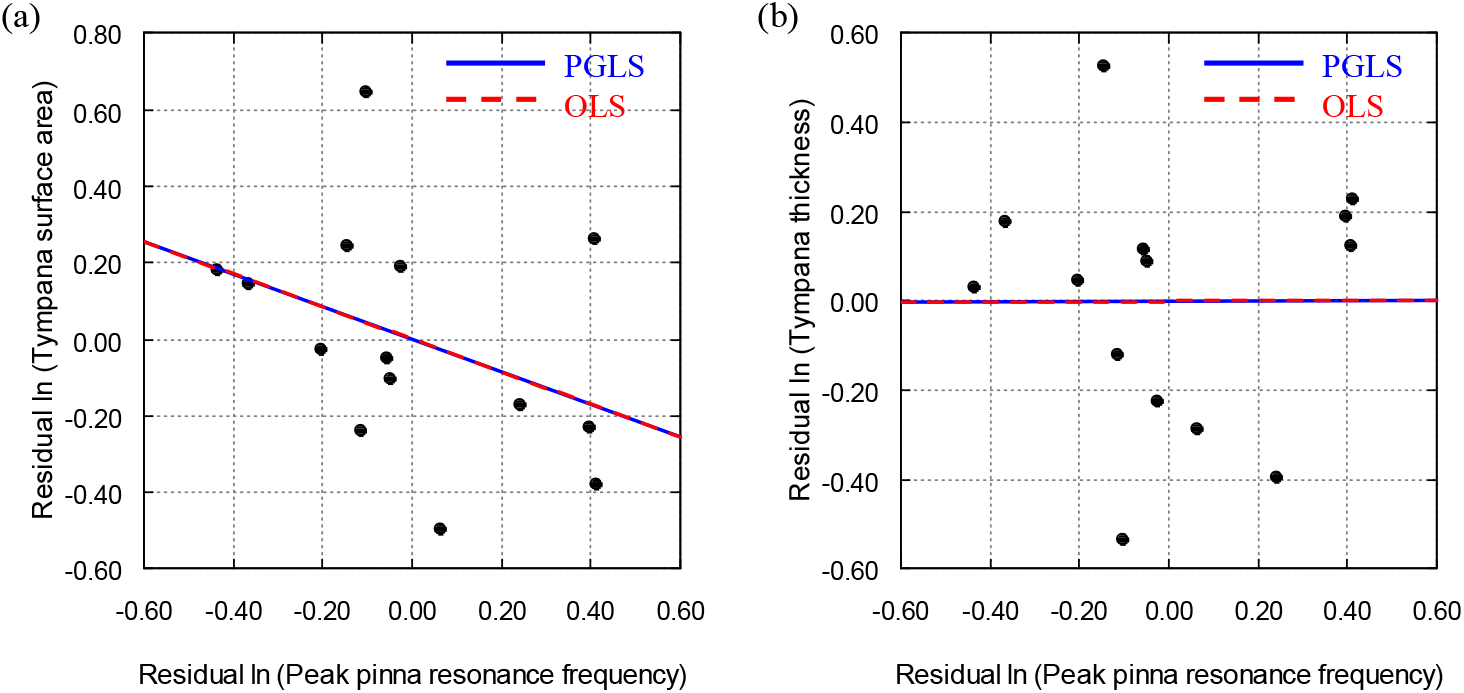
Relationships between tympanal morphology and peak pinna resonance frequency in bush-crickets with pinna cavities, controlling for body size and carrier frequency. Standardised residuals of ln-transformed peak pinna resonance frequency plotted against residual ln-transformed tympanal membrane surface area (a) and thickness (b) across 14 species with pinna cavities (unilateral and bilateral). Phylogenetic generalised least squares (PGLS, solid blue line) and ordinary least squares (OLS, dashed red line) regressions are shown.

**Table 2:**
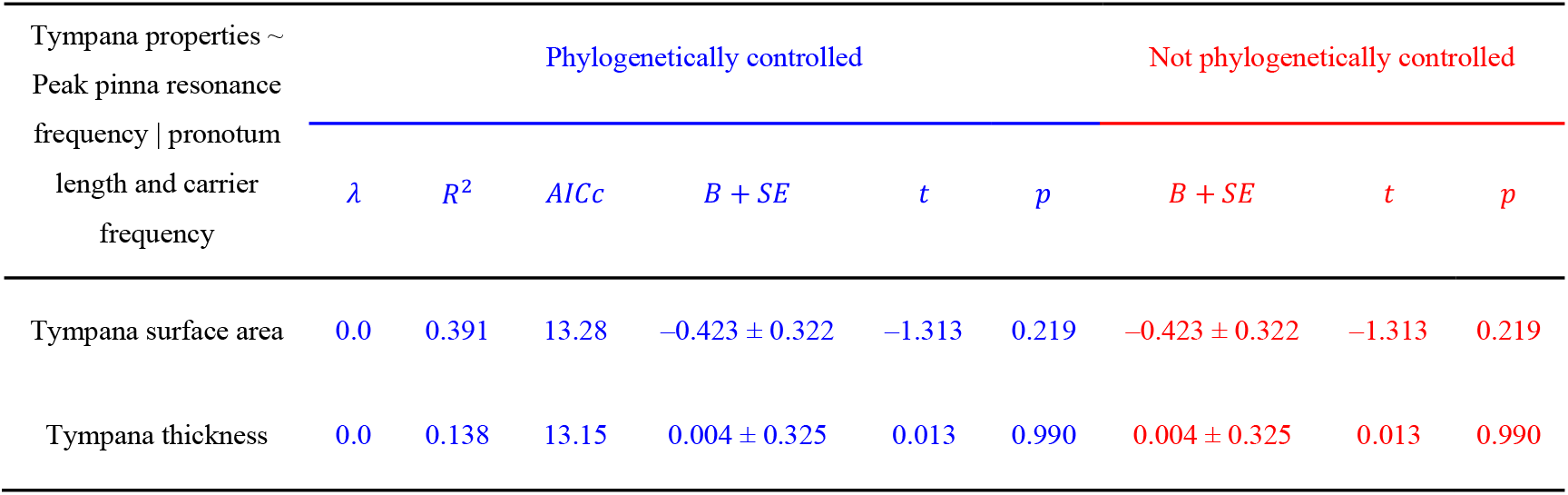
Regression analyses of tympanal properties with peak pinna resonance frequency in bush-crickets with pinna cavities, controlling for body size and carrier frequency. Results of phylogenetic generalised least squares (PGLS; blue values) and ordinary least squares (OLS; red values) models.

### 3.3. Effects of pinna presence on tympanal morphology

Tympanal membrane morphology was compared across the 18 species differing in pinna condition (exposed, unilateral and bilateral) while controlling for body size and carrier frequency (Fig. 4; Table 3). Tympanal membrane surface area did not differ significantly with pinna presence (PGLS: p = 0.617 for unilateral, p = 0.685 for bilateral; OLS: p ≥ 0.703; Fig. 4a–b; Table 3).

**Fig. 4:**
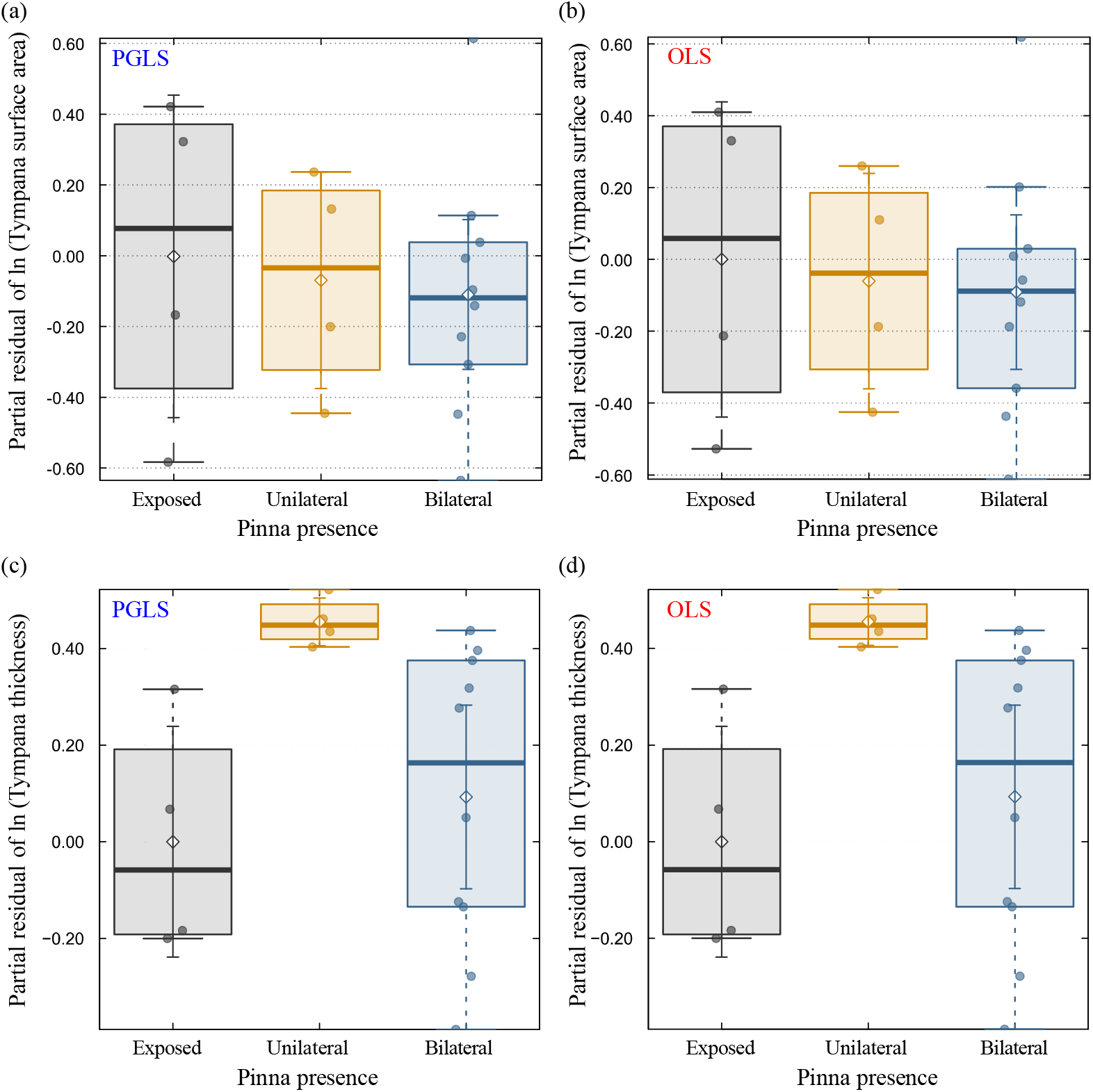
Effects of pinna presence on tympanal morphology in bush-crickets, controlling for body size and carrier frequency. Box-and-whisker plots showing variation in tympanal membrane surface area (a–b) and thickness (c–d) across species with exposed (grey), unilateral (orange), and bilateral (blue) pinnae-tympanal configurations. Phylogenetic generalised least squares (PGLS; a, c) and Ordinary least squares (OLS; b, d) estimates are shown.

**Table 3:** Regression analyses of tympanal membrane surface area and thickness in relation to pinna presence, controlling for body size and carrier frequency. Results of phylogenetic generalised least squares (PGLS; blue values) and ordinary least squares (OLS; red values) models.

| Parameter | Phylogenetically controlled |  |  |  |  |  | Not phylogenetically controlled |  |  |
| --- | --- | --- | --- | --- | --- | --- | --- | --- | --- |
| | $\lambda$ | $R^2$ | $AICc$ | $B + SE$ | $t$ | $p$ | $B + SE$ | $t$ | $p$ |
| Tympana surface area ~ pronotum length + carrier frequency + pinna presence |  |  |  |  |  |  |  |  |  |
| Intercept | 0.300 | 0.495 | 29.98 | 11.721 ± 1.025 | 11.440 | <0.001 | 11.859 ± 1.099 | 10.790 | <0.001 |
| Pronotum length |  |  |  | 1.135 ± 0.350 | 3.240 | 0.006 | 1.146 ± 0.386 | 2.970 | 0.011 |
| Carrier frequency |  |  |  | 0.092 ± 0.161 | 0.570 | 0.579 | 0.037 ± 0.171 | 0.220 | 0.832 |
| Pinnae presence:<br>unilateral |  |  |  | -0.139 ± 0.272 | -0.510 | 0.617 | -0.061 ± 0.296 | -0.200 | 0.841 |
| Pinnae presence:<br>bilateral |  |  |  | -0.094 ± 0.228 | -0.410 | 0.685 | -0.091 ± 0.234 | -0.390 | 0.703 |
| Tympana thickness ~ pronotum length + carrier frequency + pinna presence |  |  |  |  |  |  |  |  |  |
| Intercept | 0.000 | 0.427 | 19.23 | 3.042 ± 0.795 | 3.830 | 0.002 | 3.042 ± 0.795 | 3.830 | 0.002 |
| Pronotum length |  |  |  | 0.250 ± 0.279 | 0.900 | 0.386 | 0.250 ± 0.279 | 0.900 | 0.386 |
| Carrier frequency |  |  |  | -0.154 ± 0.123 | -1.250 | 0.233 | -0.154 ± 0.123 | -1.250 | 0.233 |
| Pinnae presence:<br>unilateral |  |  |  | 0.456 ± 0.2142 | 2.130 | 0.053 | 0.456 ± 0.2142 | 2.130 | 0.053 |
| Pinnae presence:<br>bilateral |  |  |  | 0.093 ± 0.169 | 0.550 | 0.592 | 0.093 ± 0.169 | 0.550 | 0.592 |

However, tympanal membrane thickness varied with pinna condition, with unilateral pinna associated with significantly thicker tympana compared with species lacking pinnae (PGLS: *B* = 0.456 ± 0.214, *t* = 2.130, *p* = 0.053; OLS identical; Fig. 4c–d; Table 3). No such effect was detected for bilateral pinnae (*p* = 0.592; OLS identical; Fig. 4c–d; Table 3). Phylogenetic signal was moderate for surface area (λ = 0.300) and absent for thickness (λ = 0.000).

### 3.4. Asymmetries in tympanal morphology of unilateral pinna species

Analyses of the four Phaneropterinae taxa with unilateral pinnae revealed consistent morphological differences between the pinna-covered and exposed tympana (Fig. 5). Tympanal membrane surface area was generally larger on the pinna-covered side, with increases ranging from +10.03% (*Phaulula galeata*) to +23.58% (*Arnobia pilipes*). In contrast, tympanal membrane thickness was generally reduced on the pinna-covered side, with decreases ranging from –20.24% (*Phygela marginata*) to –23.92% (*Stictophaula sp*.).

**Fig. 5:**
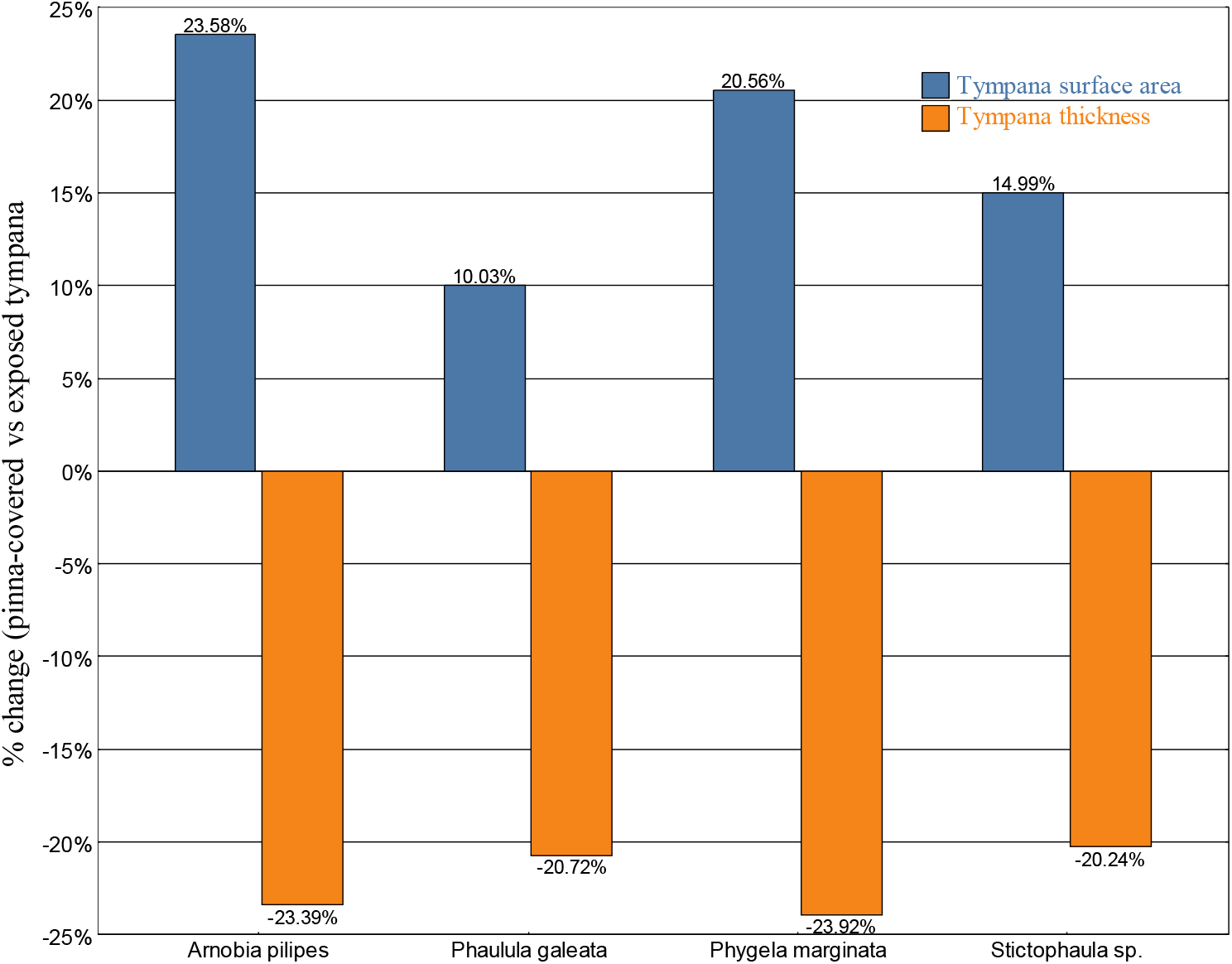
Within-species comparisons of tympanal properties between pinna-covered and exposed ears in bush-crickets with unilateral pinna. Percentage change in tympanal membrane surface area (blue bars) and thickness (orange bars) between pinna-covered and exposed tympana for five species with unilateral pinna (*Arnobia pilipes, Phaulula galeata, Phygela marginata*, and *Stictophaula sp*.). Positive values indicate larger or thicker tympana under the pinna relative to the exposed side, and negative values indicate smaller or thinner tympana under the pinna.

## 4. Discussion

This comparative study provides new insights into the relationships between body size, acoustic properties, and tympanal morphology in bush-crickets, while also testing the potential influence of geography and auditory pinnae on tympanal structure. Across the 18 species examined, we found that body size, measured via pronotum length, was a significant predictor of carrier frequency, consistent with general scaling relationships observed in bush-cricket bioacoustics [34–37]. Larger species produced lower-frequency calls, a relationship that persisted after controlling for phylogeny. This finding supports the view that body size imposes fundamental biomechanical constraints on sound production and that the frequency ranges used in communication are inherently linked to overall morphology [37].

Body size also predicted tympanal membrane surface area but not thickness. Larger species possessed proportionally larger tympanal membranes, suggesting that ear morphology co-scales with body size to maintain sensitivity to conspecific signals. In contrast, tympanal thickness was unrelated to size, implying that this trait may be under different selective pressures, perhaps linked more directly to tuning or to mechanical protection of the membrane. When controlling for size, tympanal surface area showed no significant association with carrier frequency, whereas tympanal thickness exhibited a weak, non-significant negative relationship with carrier frequency. These results suggest that tympanal morphology alone does not determine acoustic output. Although tympanic membranes typically exhibit a broad frequency response suited to detecting a range of biologically relevant sounds, they also show enhanced sensitivity around the species’ calling frequency. Therefore, other components of the auditory system, such as tracheal resonances or pinna cavities, are likely to contribute to fine-tuning and directional sensitivity.

In species with pinna cavities, neither tympanal surface area nor thickness correlated with peak pinna resonance frequency. This indicates that tympanal morphology and cavity resonance may evolve semi-independently, with cavity geometry rather than tympanal structure dictating the resonance properties. Importantly, phylogenetic signal was absent in these models, supporting the view that these traits may be labile and shaped by ecological rather than phylogenetic constraints.

The presence of pinnae did not significantly affect tympanal surface area across the broader dataset, but tympanal thickness did vary with pinna configuration. Species with unilateral pinnae had, on average, thicker tympana than species lacking pinnae, whereas bilateral pinnae showed no detectable effect on tympanal dimensions. All unilateral taxa in this study belonged to the Phaneropterinae subfamily, so this pattern could in principle reflect clade-specific morphology. However, the effect of unilateral pinnae on thickness remained when phylogeny was incorporated via PGLS (λ ≈ 0), suggesting that it is not solely a by-product of shared ancestry within this group. One interpretation is that tympanal dimensions are generally constrained within a functional range by the mechanical requirements of sound detection, membranes must remain thin and compliant enough for sensitivity, yet structurally robust, while unilateral pinnae impose additional, asymmetric mechanical demands that are met by a modest global thickening of the tympanum. Consequently, this finding challenges the traditional view that auditory pinnae primarily evolved to provide mechanical protection to tympanal membranes [14–16], instead supports recent work emphasising their role in acoustic gain and predator detection [11,12].

By contrast, within-species comparisons in Phaneropterinae taxa with unilateral pinnae revealed consistent morphological asymmetries between covered and exposed tympana. Despite unilateral species having thicker tympana overall at the interspecific level, pinna-covered tympana exhibited larger surface areas but thinner membranes than their exposed counterparts. These asymmetries suggest that the presence of a pinna may influence tympanal development or mechanical properties in a localised manner. Larger surface area may enhance sound capture under the pinna, whereas reduced thickness could increase sensitivity to ultrasonic frequencies, potentially complementing the cavity’s acoustic gain [38,39].

Taken together, these findings demonstrate that tympanal morphology in bush-crickets arises from multiple interacting influences. Tympanal surface area scales predictably with body size, reflecting biomechanical constraints on acoustic sensitivity, whereas thickness remains decoupled from size and more evolutionarily labile. The absence of broad effects of pinna presence, coupled with consistent asymmetries in unilateral species, suggests that pinnae influence tympanal morphology in a localised rather than generalised manner. Thus, tympanal structures reflect a complex interplay of scaling, ecological divergence, and structural innovation rather than simple co-variation with acoustic output.

## 5. Conclusion

This comparative analysis of bush-cricket tympanal membranes reveals that ear morphology is governed by multiple interacting factors. Scaling effects were evident, with tympanal surface area increasing predictably with body size, thereby maintaining sensitivity across species of different sizes, while thickness remained decoupled from size and more variable. The role of auditory pinnae was more nuanced. While pinnae did not affect tympanal surface area across species, unilateral pinnae were associated with thicker tympana, whereas bilateral pinnae showed no change. These contrasting patterns point to condition-specific effects and challenge the idea that pinnae originally evolved as protective structures. However, within-species comparisons in taxa with unilateral pinnae consistently revealed asymmetries: pinna-covered tympana were larger in surface area but thinner than exposed tympana. These localised effects imply that pinnae can shape tympanal development and mechanics in ways that may complement their role in generating acoustic gain.

The comparative patterns identified here provide a foundation for future work investigating how tympanal morphology interacts with other elements of the bush-cricket auditory system, such as tracheal pathways, cuticular mechanics, and neural processing. Incorporating wider taxonomic sampling, behavioural assays, and *in vivo* vibrational measurements will help clarify how morphological variation translates into functional differences in sensitivity and tuning. Beyond biological relevance, our findings have implications for engineering applications. The diversity of tympanal structures across species offers natural templates for bio-inspired acoustic sensors, particularly in the ultrasonic range. The consistent scaling of tympanal surface area with body size and the localised asymmetries associated with pinnae provide design principles for creating lightweight, frequency-tuned devices. Such approaches could inform the development of miniature microphones for robotics, surveillance, or medical diagnostics where sensitivity to specific frequency bands is critical.

## Acknowledgements

This research is part of the project “Biophysical and ecological function of microscale ears using scaled 3D prints” funded by the Leverhulme Trust, Grant RPG-2023-204 to Fernando Montealegre-Z; and was also funded by the Natural Environment Research Council (NERC), grant DEB-1937815 to Fernando Montealegre-Z.

## Author contributions

**Md Niamul Islam:** Conceptualization, Data curation, Formal analysis, Investigation, Methodology, Software, Validation, Visualization, Writing – original draft, Writing – review & editing.

**Lewis Holmes:** Conceptualization, Data curation, Investigation, Methodology, Writing – original draft, Writing – review & editing.

**Dominic Rooke:** Investigation, Methodology, Software, Writing – review & editing.

**Fabio A. Sarria-S:** Investigation, Methodology, Software, Writing – review & editing.

**Fernando Montealegre-Z:** Conceptualization, Funding acquisition, Project administration, Resources, Supervision, Validation, Writing – review & editing.

All authors gave final approval for publication.

## Data availability

Data supporting the findings of this study are provided in the Supplementary Documents and available upon request from the corresponding authors.

