## Supplementary Document 1 for "Comparative analysis of morphological and acoustic correlates of bush-cricket tympanic membranes"

**Table S1:** Morphological and acoustic traits of bush-cricket species examined in this study. For each species, the table lists geographic location, subfamily, presence or absence of cuticular pinnae, carrier frequency of the call, pronotum length, mean tympanal membrane surface area, and mean tympanal membrane thickness.

| Species | Location | Subfamily | Pinnae type | Carrier frequency (kHz) | Pronotum length (mm) | Mean tympana surface area (µm²) | Mean tympana thickness (µm) |
|---|---|---|---|---|---|---|---|
| *Arachnoscelis sp.* | South America | Meconematinae | Bilateral | 83.3 | 4.6 | 1,169,955.93 | 21.32 |
| *Arnobia pilipes* | Asia | Phaneropterinae | Unilateral | 26.4 [2] | 4.9 | 1,278,630.35 | 28.16 |
| *Balboana tibialis* | South America | Pseudophyllinae | Bilateral | 13.7 [5] | 11.2 | 2,204,844.61 | 39.66 |
| *Chibchella nigrospecula* | South America | Pseudophyllinae | Bilateral | 25 [3] | 7.4 | 1,592,153.93 | 18.39 |
| *Copiphora gorgonensis* | South America | Conocephalinae | Bilateral | 23.7 [4] | 8.9 | 1,257,312.79 | 15.07 |
| *Elimaea signata* | Asia | Phaneropterinae | Bilateral | 14.3 [2] | 4.3 | 854,592.34 | 29.75 |
| *Haenschiella sp.* | South America | Pseudophyllinae | Bilateral | 78.2 | 4.9 | 968,515.83 | 23.16 |
| *Leptoderes ornatipennis* | Asia | Phaneropterinae | Exposed | 8.4 | 8.7 | 1,474,803.50 | 35.56 |
| *Mecopoda elongata* | Asia | Mecopodinae | Exposed | 15.9 | 8.9 | 2,889,944.44 | 25.28 |
| *Moncheca elegans* | South America | Conocephalinae | Bilateral | 38 | 5.9 | 1,024,815.33 | 24.58 |
| *Phaulula galeata* | Asia | Phaneropterinae | Unilateral | 21.3 [2] | 4.4 | 716,804.25 | 31.91 |
| *Phlugis poecilla* | South America | Meconematinae | Exposed | 44.2 | 3.6 | 416,243.96 | 13.17 |
| *Phygela marginata* | Asia | Phaneropterinae | Unilateral | 11.7 [2] | 5.3 | 1,168,524.38 | 33.63 |
| *Phyllomimus detersus* | Asia | Pseudophyllinae | Bilateral | 7.1 [2] | 6.5 | 906,920.30 | 26.01 |
| *Ragoniella pulchella* | South America | Conocephalinae | Bilateral | 27.6 | 6.3 | 714,463.38 | 17.58 |
| *Satizabalus jorgevargasi* | South America | Pseudophyllinae | Bilateral | 18.7 [1] | 5.2 | 1,934,015.49 | 15.25 |
| *Stictophaula sp.* | Asia | Phaneropterinae | Unilateral | 25.1 [2] | 5.4 | 718,740.64 | 30.82 |
| *Stilpnochlora sp.* | South America | Phaneropterinae | Exposed | 21.5 | 7.2 | 2,115,788.91 | 17.8 |

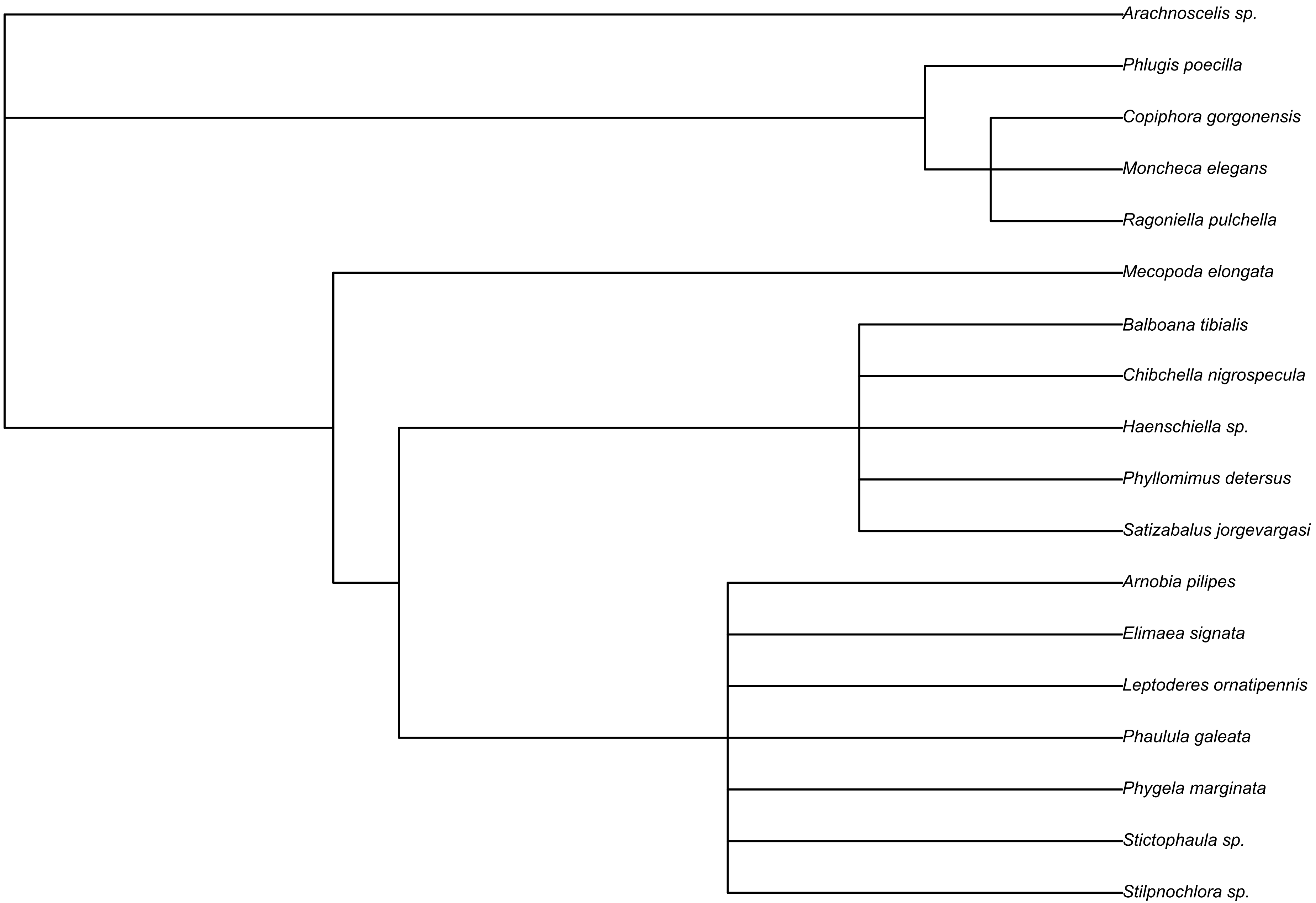


**Fig. S1:** Phylogenetic relationships of the bush-cricket species examined in this study. The tree illustrates the evolutionary placement of species for which morphological and acoustic traits were measured. The phylogeny is presented to provide context for phylogeny-controlled comparative analyses of tympanal morphology with pronotum length and carrier frequency.

**Table S2:** Frequency of maximum gain at pinna resonance for bush-cricket species with pinna cavity.

| Species | Frequency of max gain at pinna resonance (kHz) |
|---|---|
| *Arachnoscelis sp.* | 50.63 |
| *Arnobia pilipes* | 140.63 |
| *Balboana tibialis* | 47.50 |
| *Chibchella nigrospecula* | 66.33 |
| *Copiphora gorgonensis* | 75.50 |
| *Elimaea signata* | 95.63 |
| *Haenschiella sp.* | 71.25 |
| *Moncheca elegans* | 71.48 |
| *Phaulula galeata* | 157.73 |
| *Phygela marginata* | 71.48 |
| *Phyllomimus detersus* | 86.25 |
| *Ragoniella pulchella* | 80.63 |
| *Satizabalus jorgevargasi* | 86.20 |
| *Stictophaula sp.* | 131.95 |

**Table S3:** Comparison of tympanal membrane morphology in Phaneropterinae bush-cricket species with exposed tympana on one side and pinna-covered tympana on the other side.

| Species | Exposed tympana surface area ($\mu m^2$) | Pinnae-covered tympana surface area ($\mu m^2$) | Exposed tympana thickness ($\mu m$) | Pinna-covered tympana thickness ($\mu m$) |
|---|---|---|---|---|
| *Arnobia pilipes* | 1,143,780.67 | 1,413,480.02 | 32.36 | 24.79 |
| *Phaulula galeata* | 682,585.70 | 751,022.79 | 35.71 | 28.31 |
| *Phygela marginata* | 1,059,591.20 | 1,277,457.55 | 38.67 | 29.42 |
| *Stictophaula sp.* | 668,614.36 | 768,866.92 | 34.43 | 27.46 |
