## Supplementary Document 2 for "Comparative analysis of morphological and acoustic correlates of bush-cricket tympanic membranes"

*Arachnoscelis sp.*

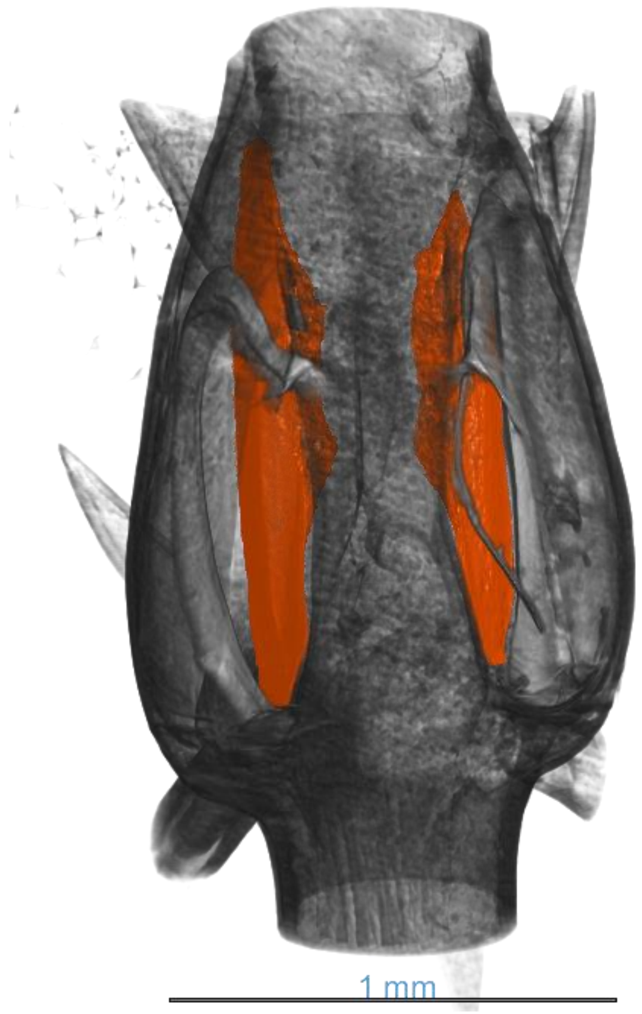


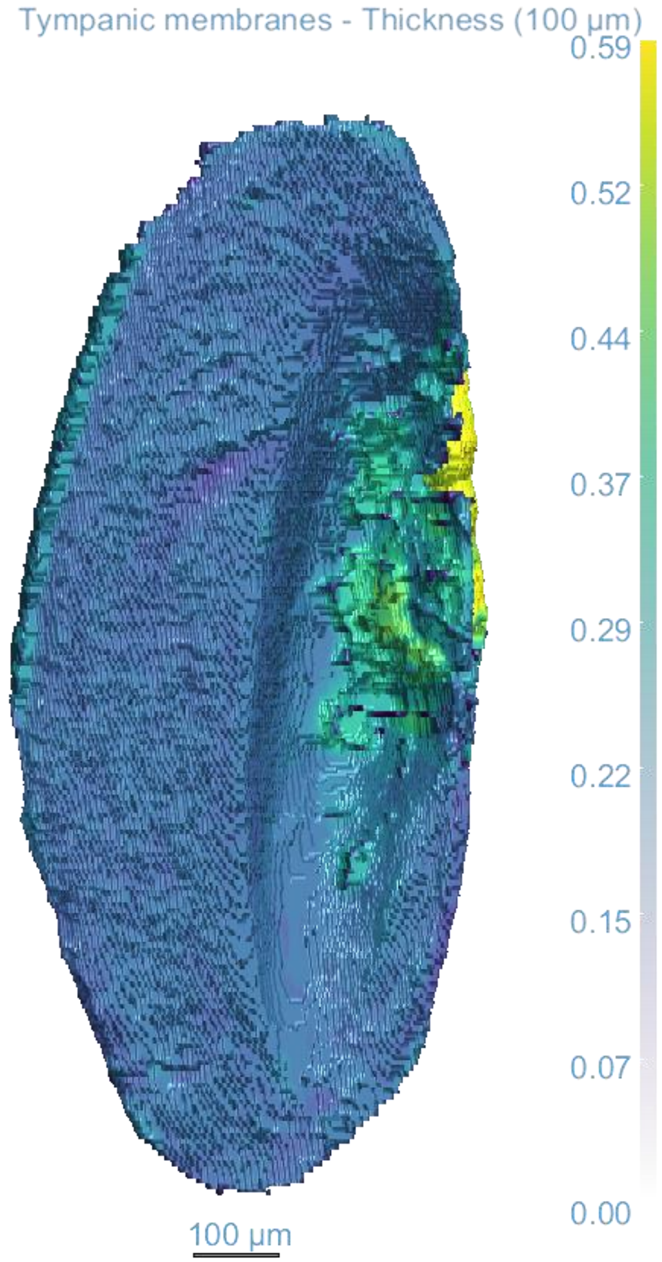

*Arachnoscelis sp.*

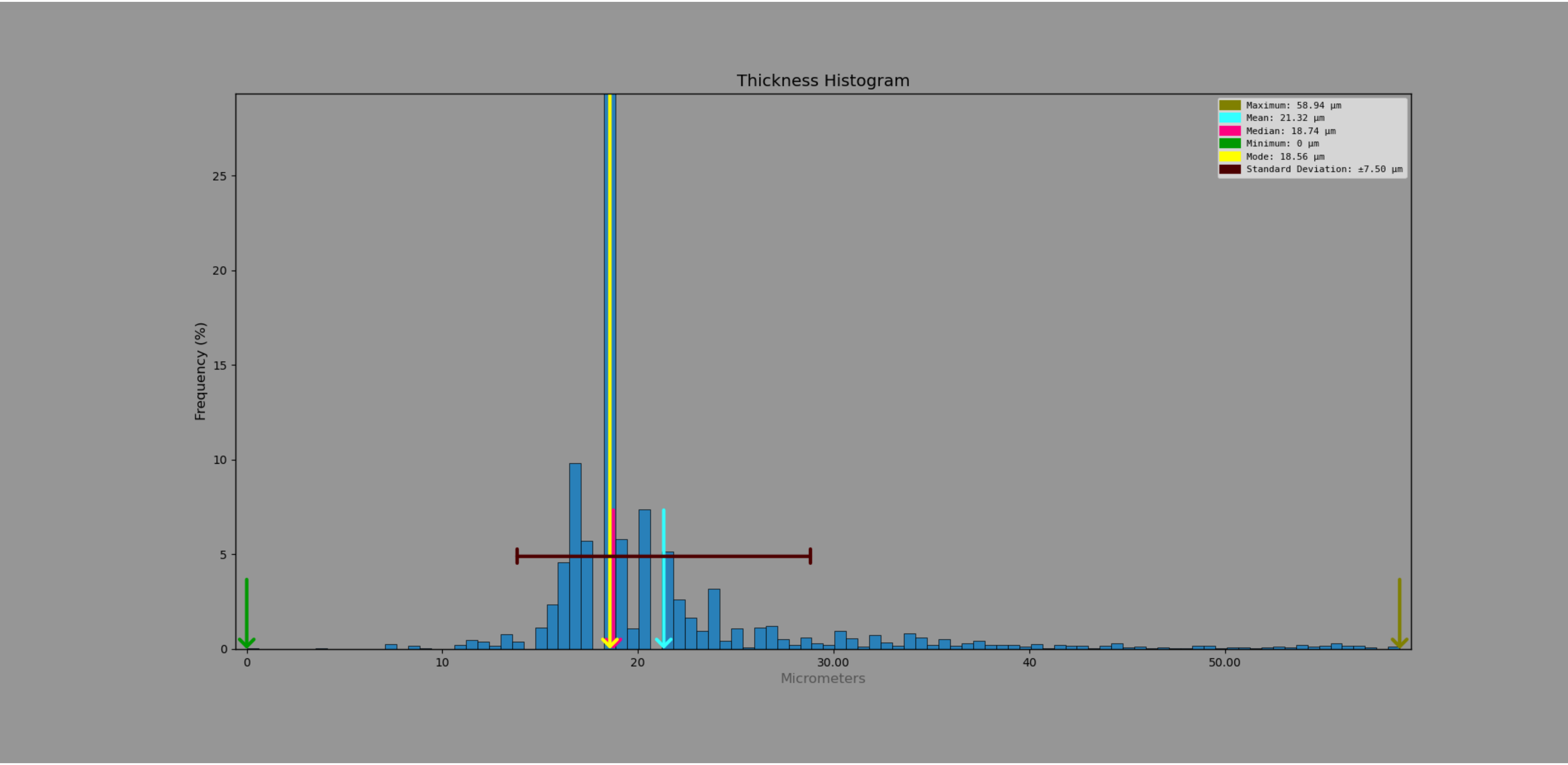

*Arnobia pilipes*

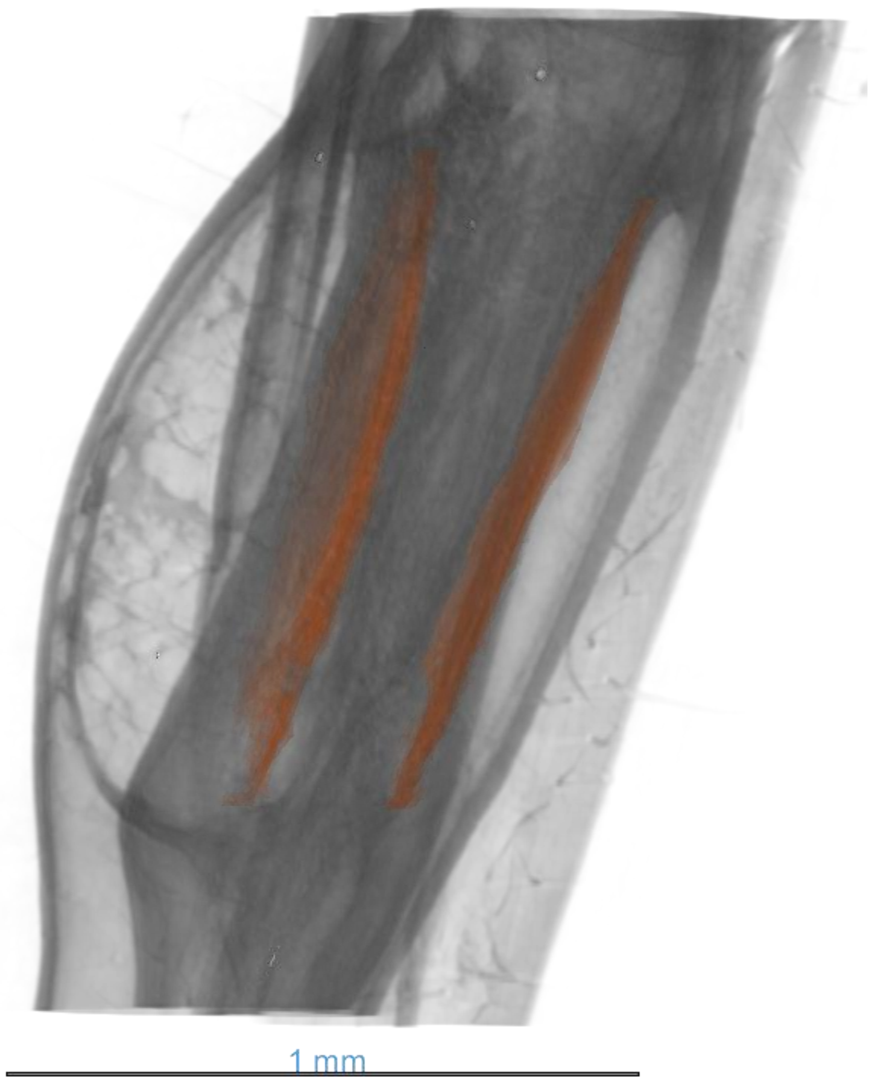


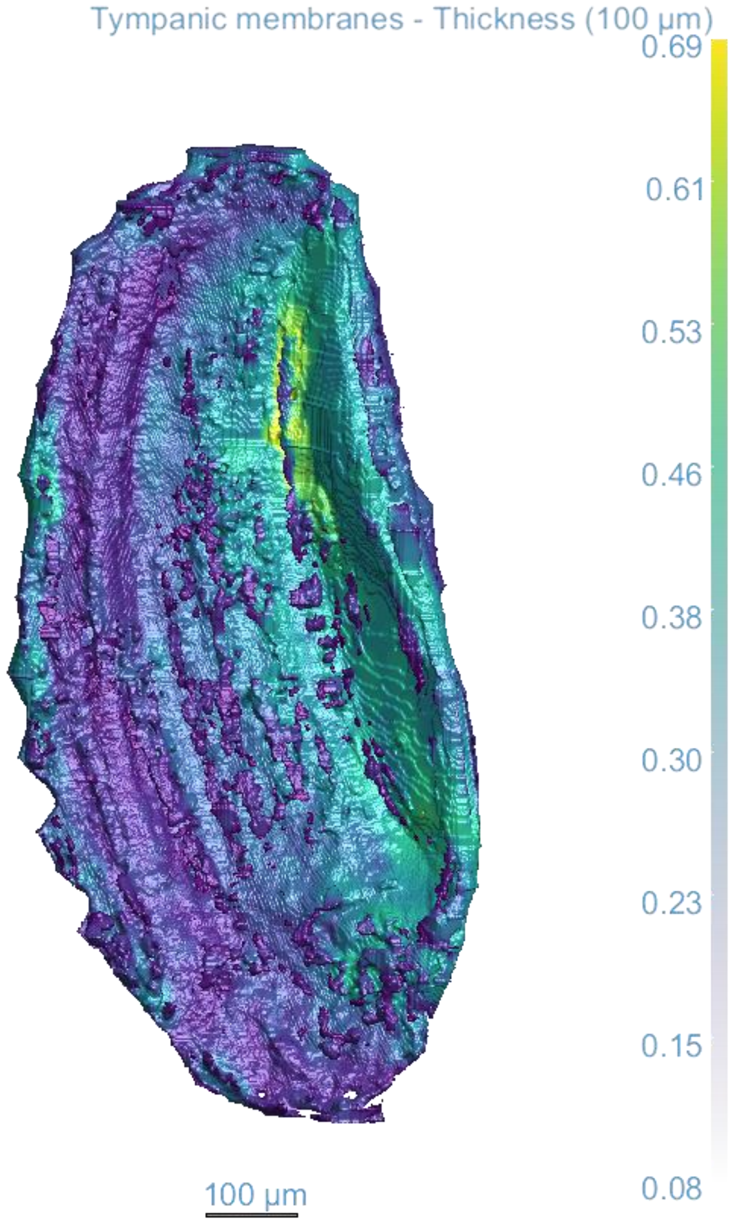

*Arnobia pilipes*

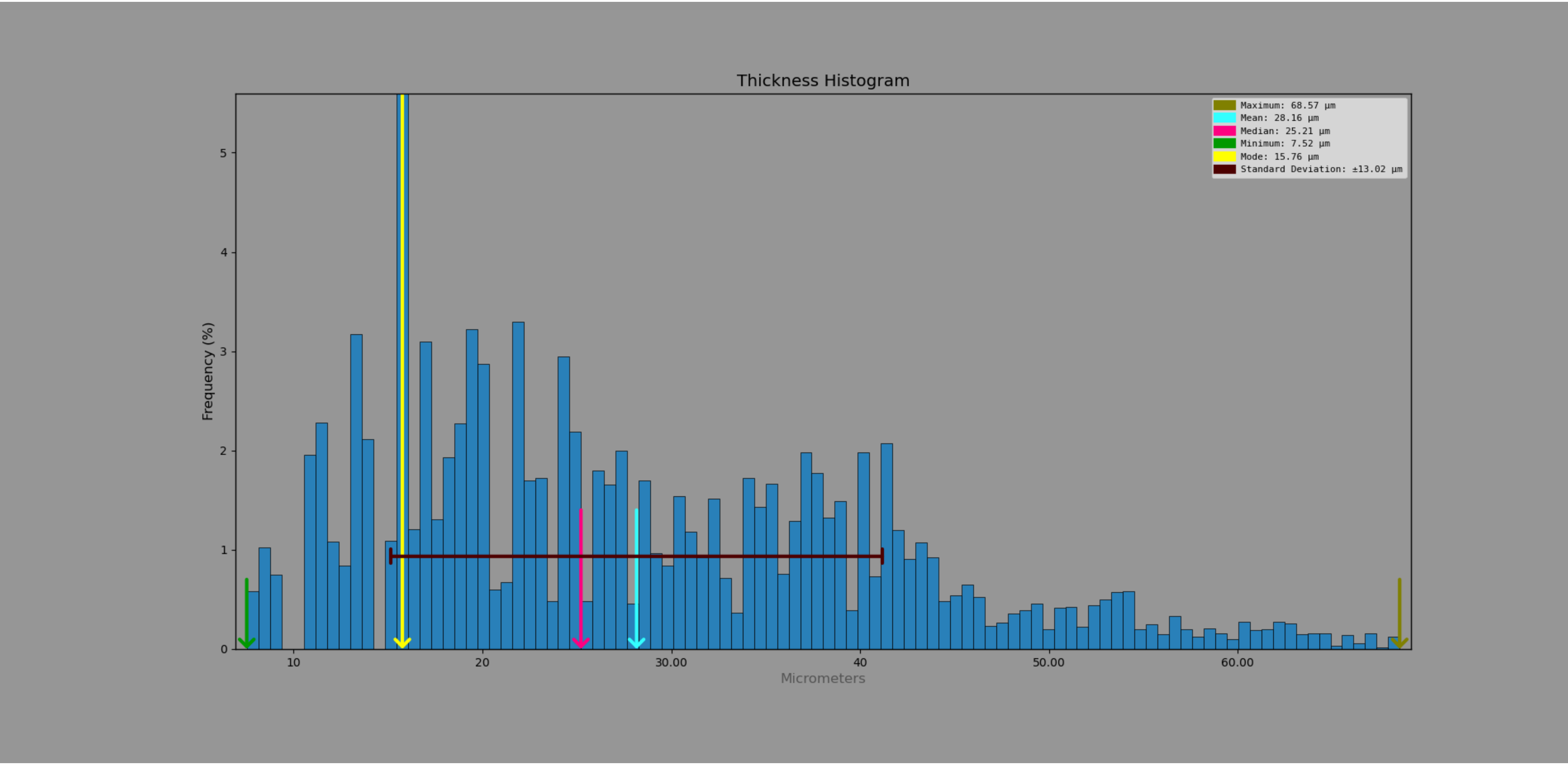

*Arnobia pilipes* – exposed tympana

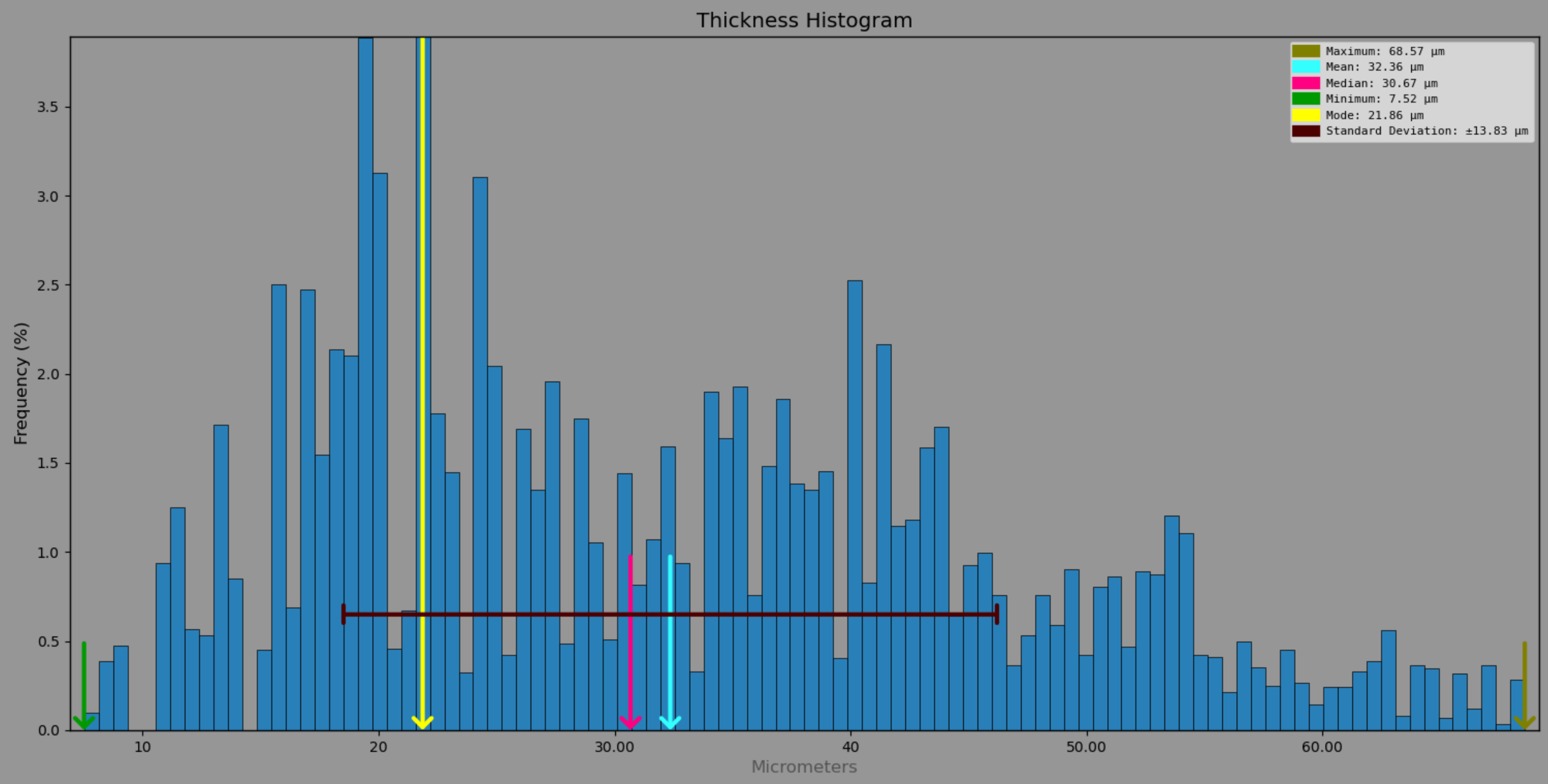

### *Arnobia pilipes* – pinna covered tympana

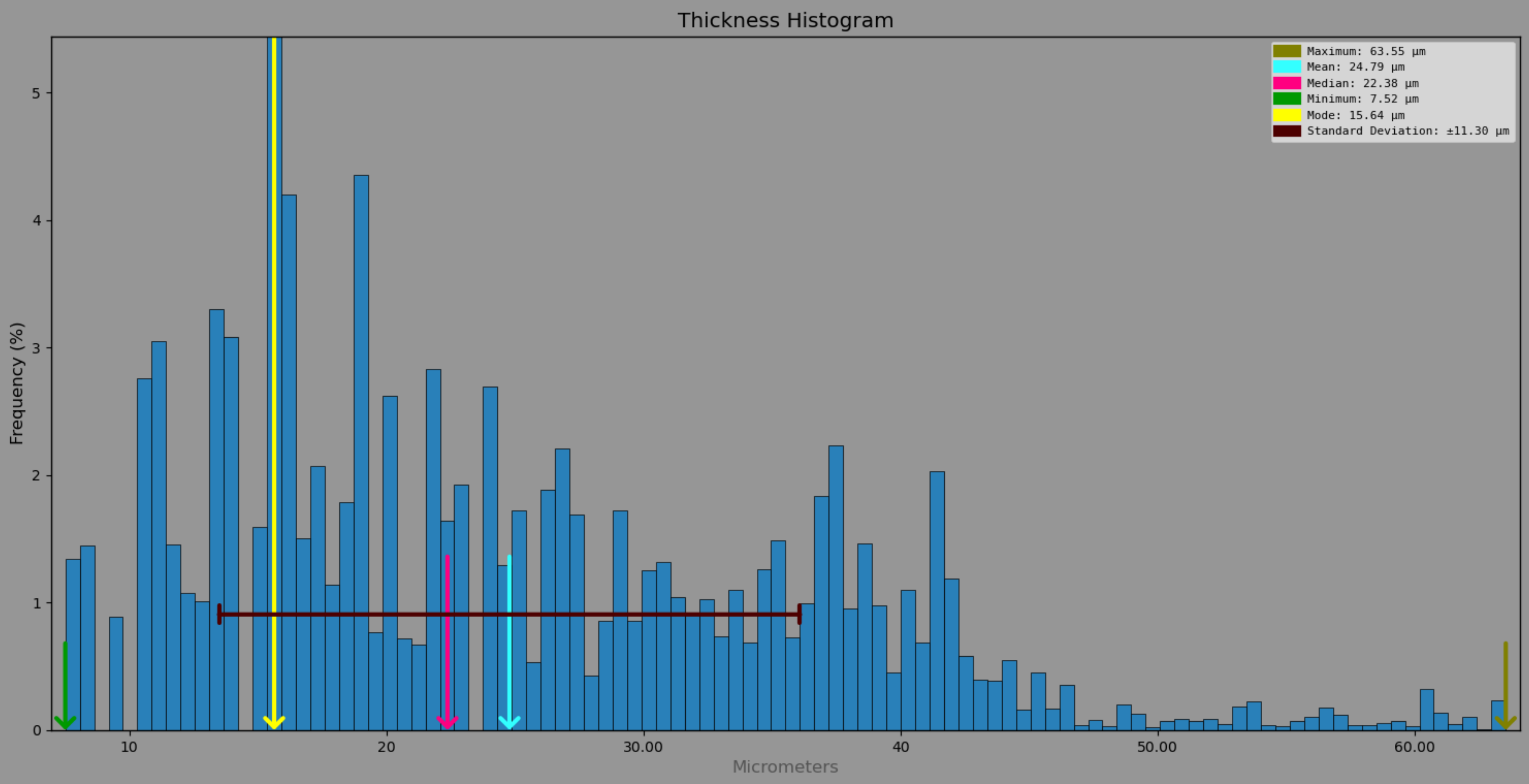

*Balboana tibialis*

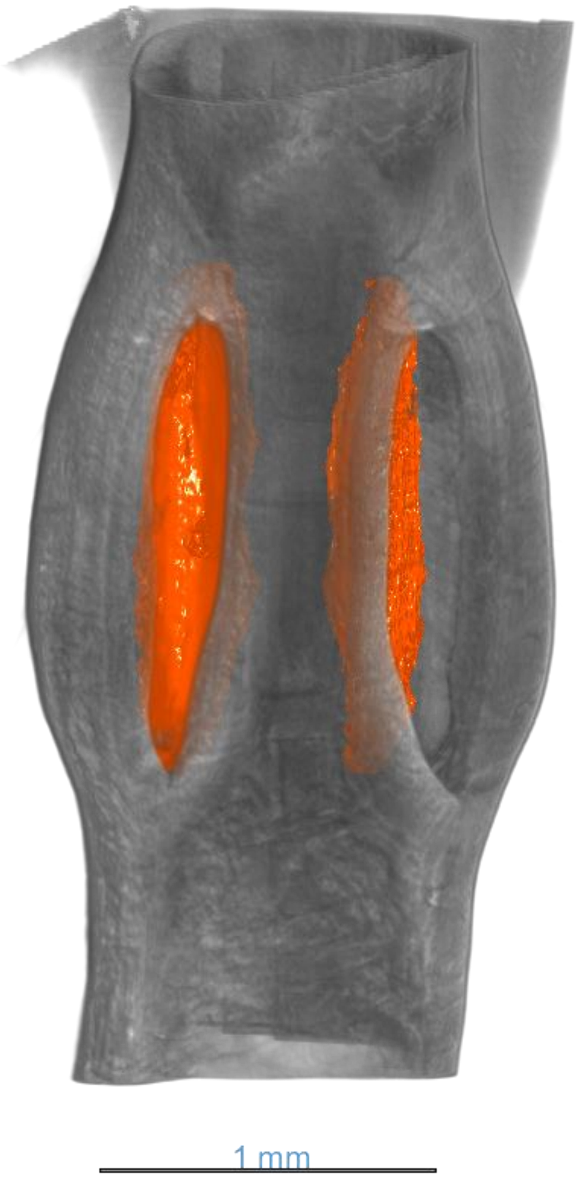


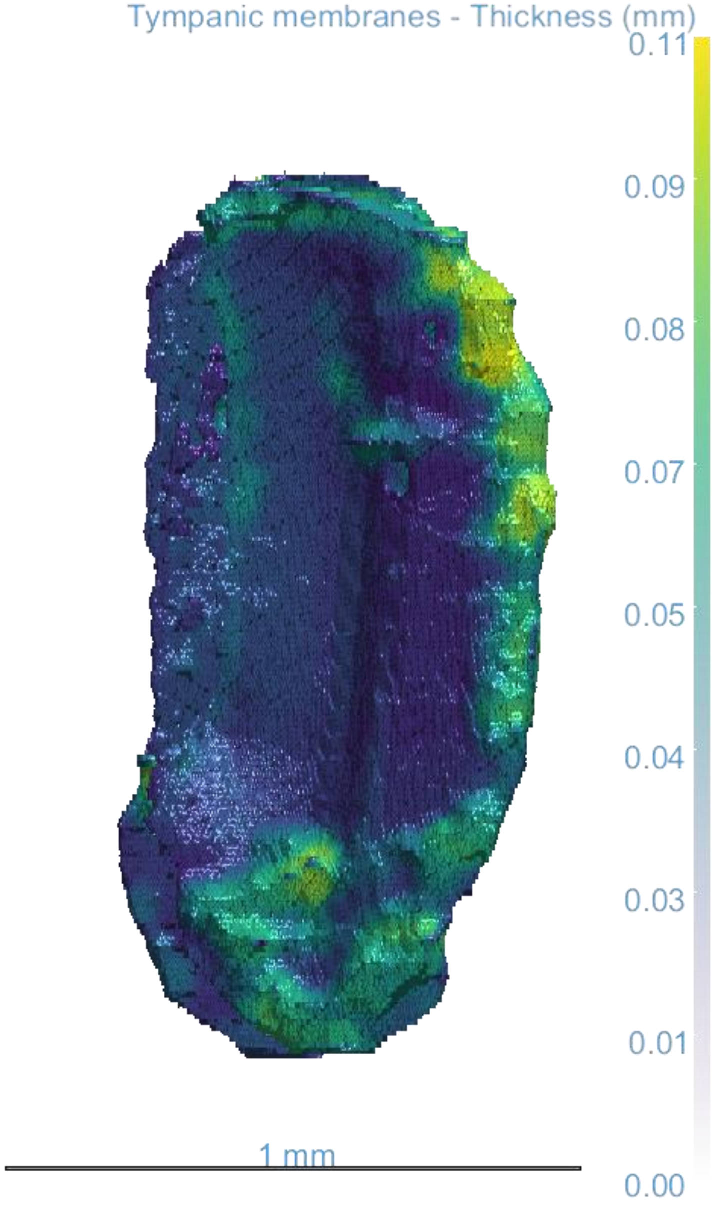

*Balboana tibialis*

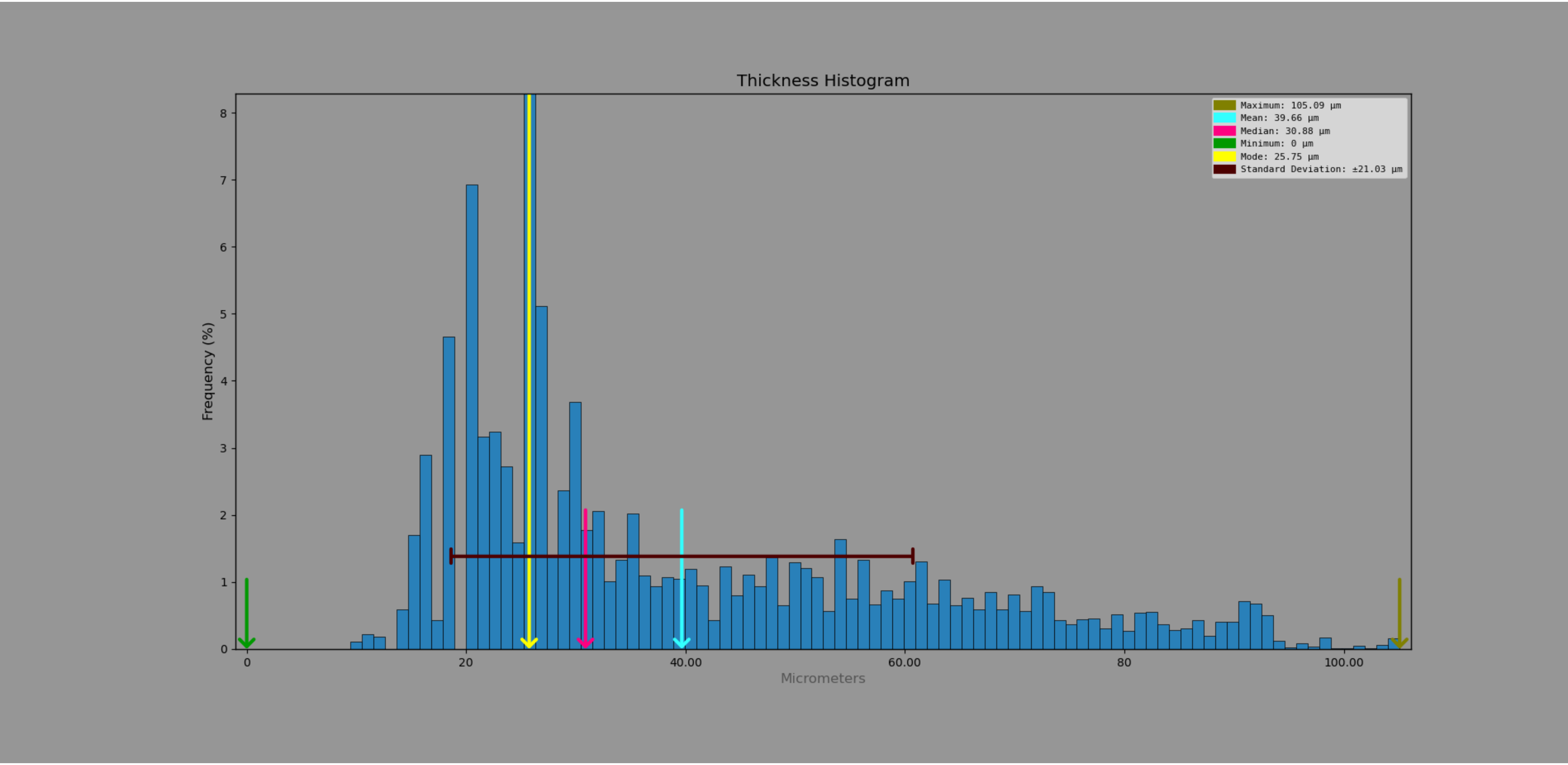

*Chibchella nigrospecula*

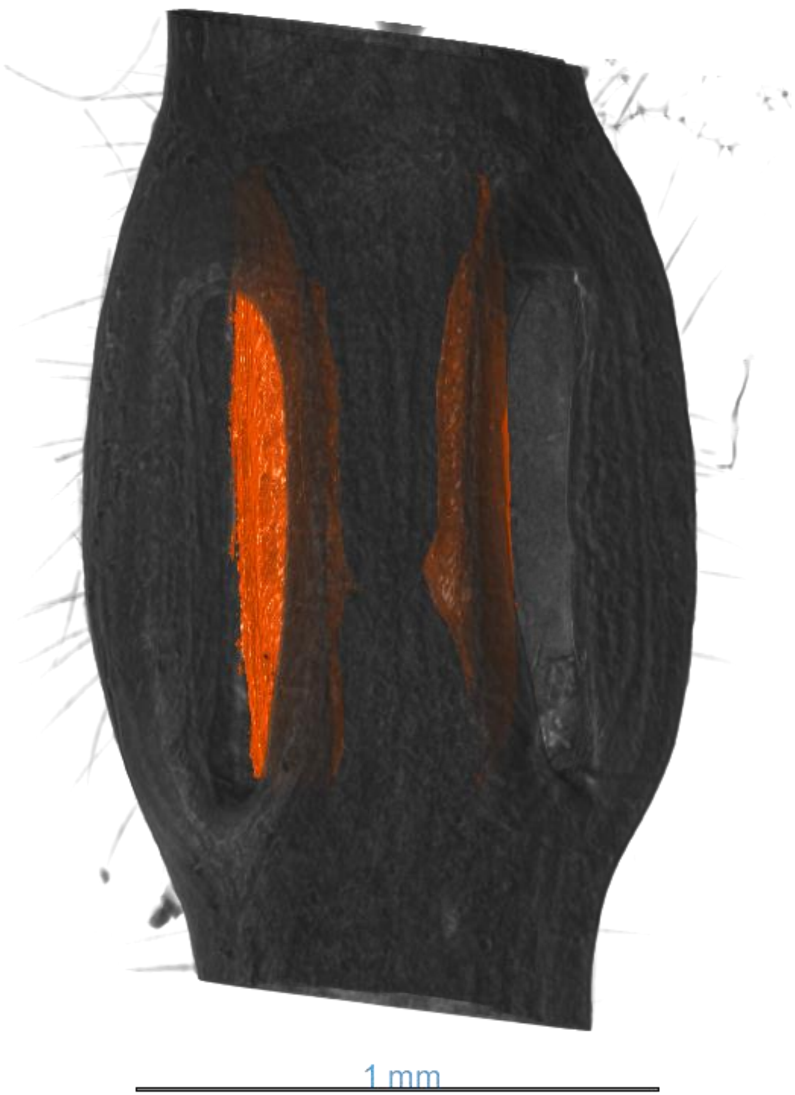


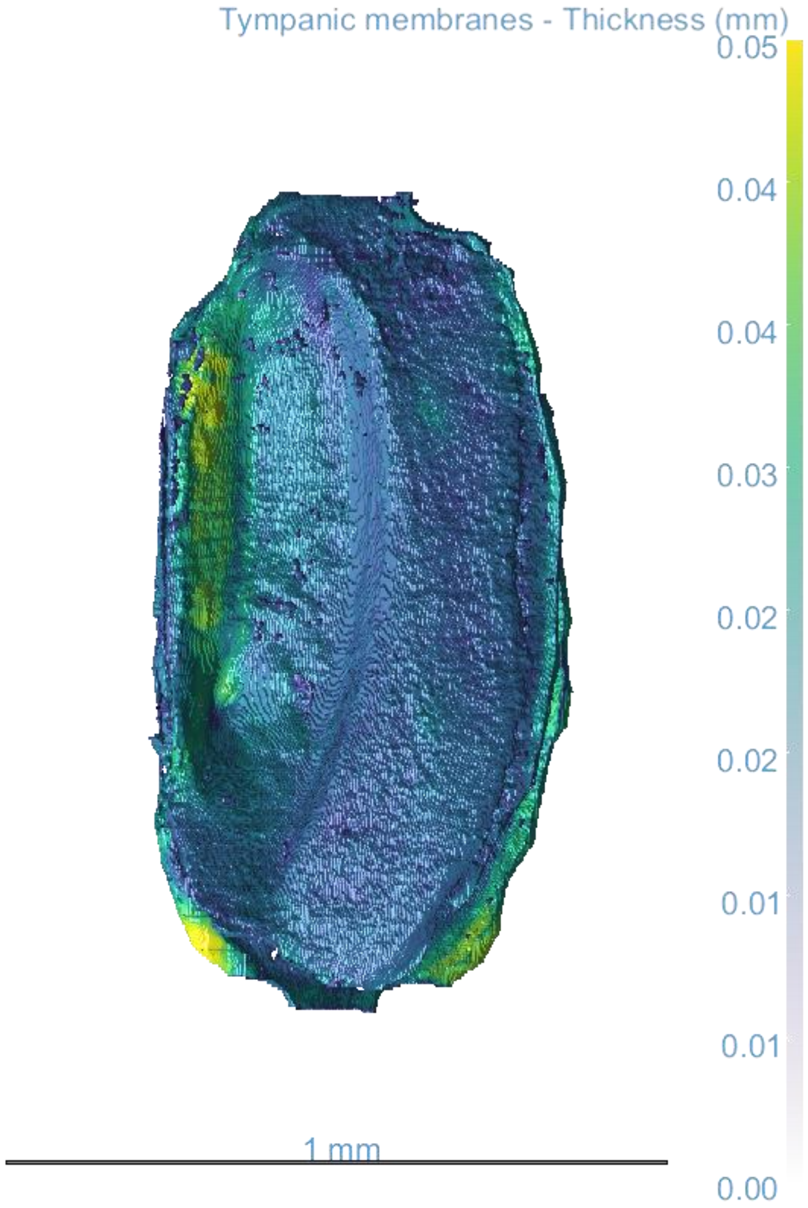

*Chibchella nigrospecula*

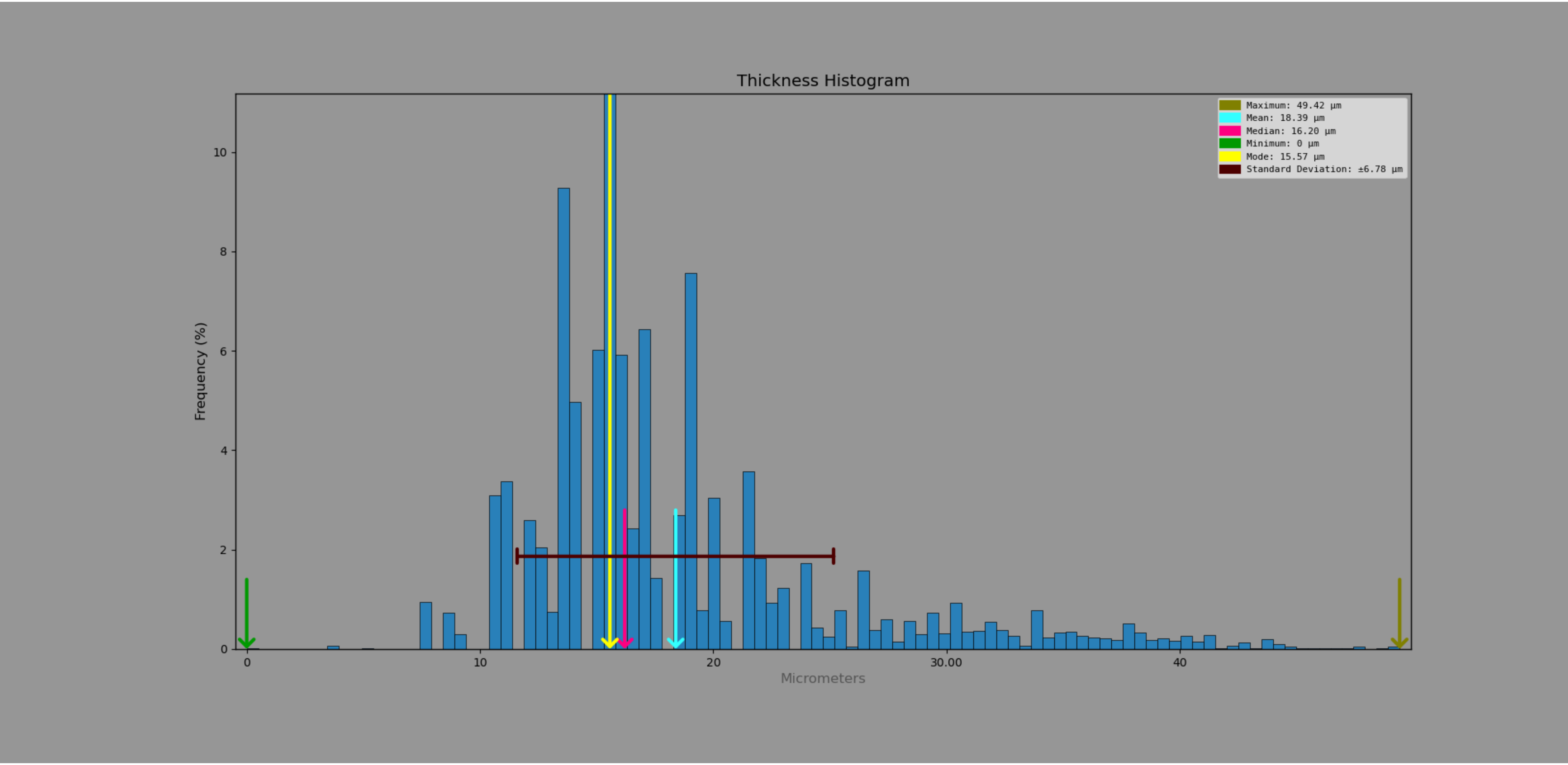

*Copiphora gorgonensis*

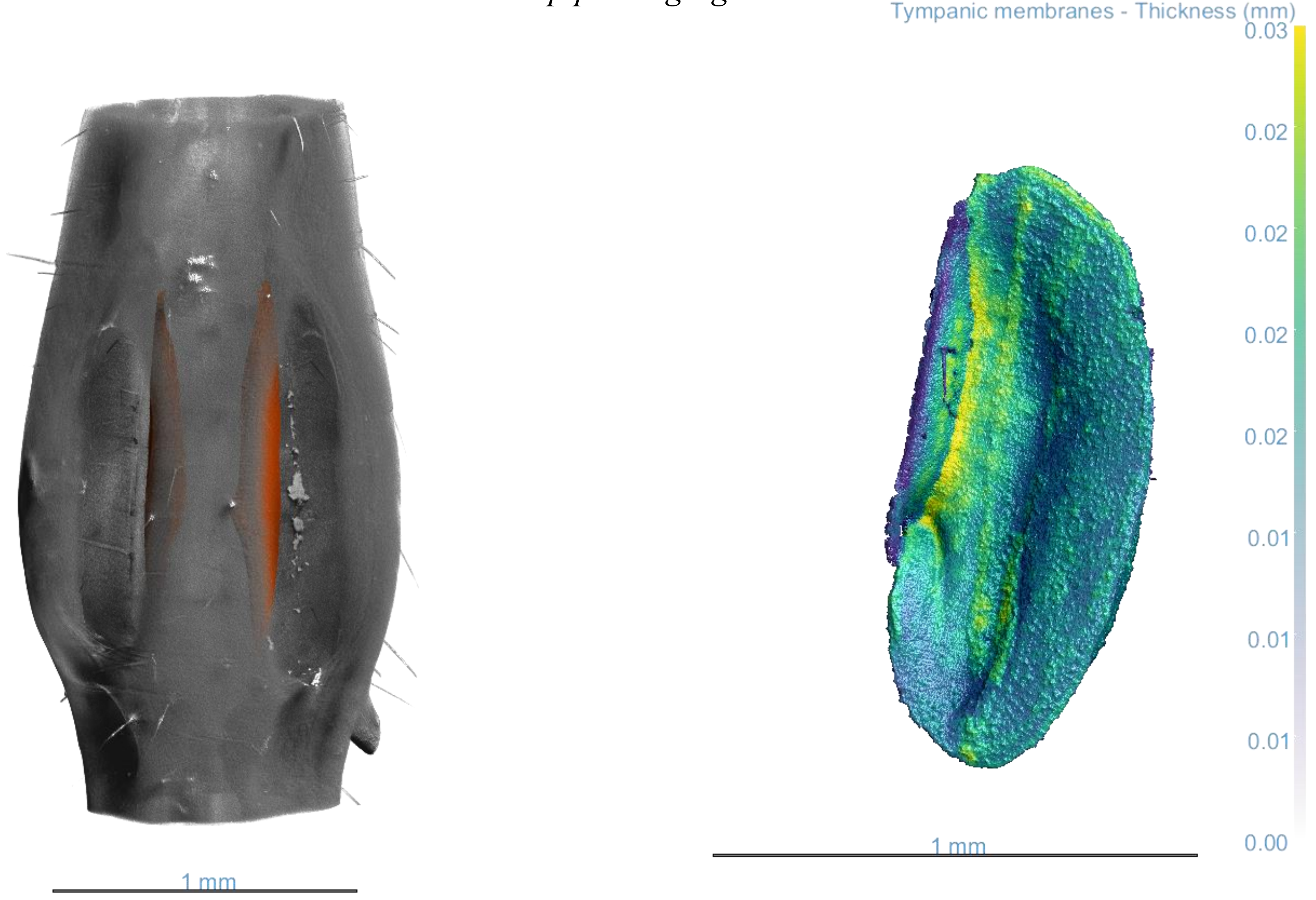

### Copiphora gorgonensis

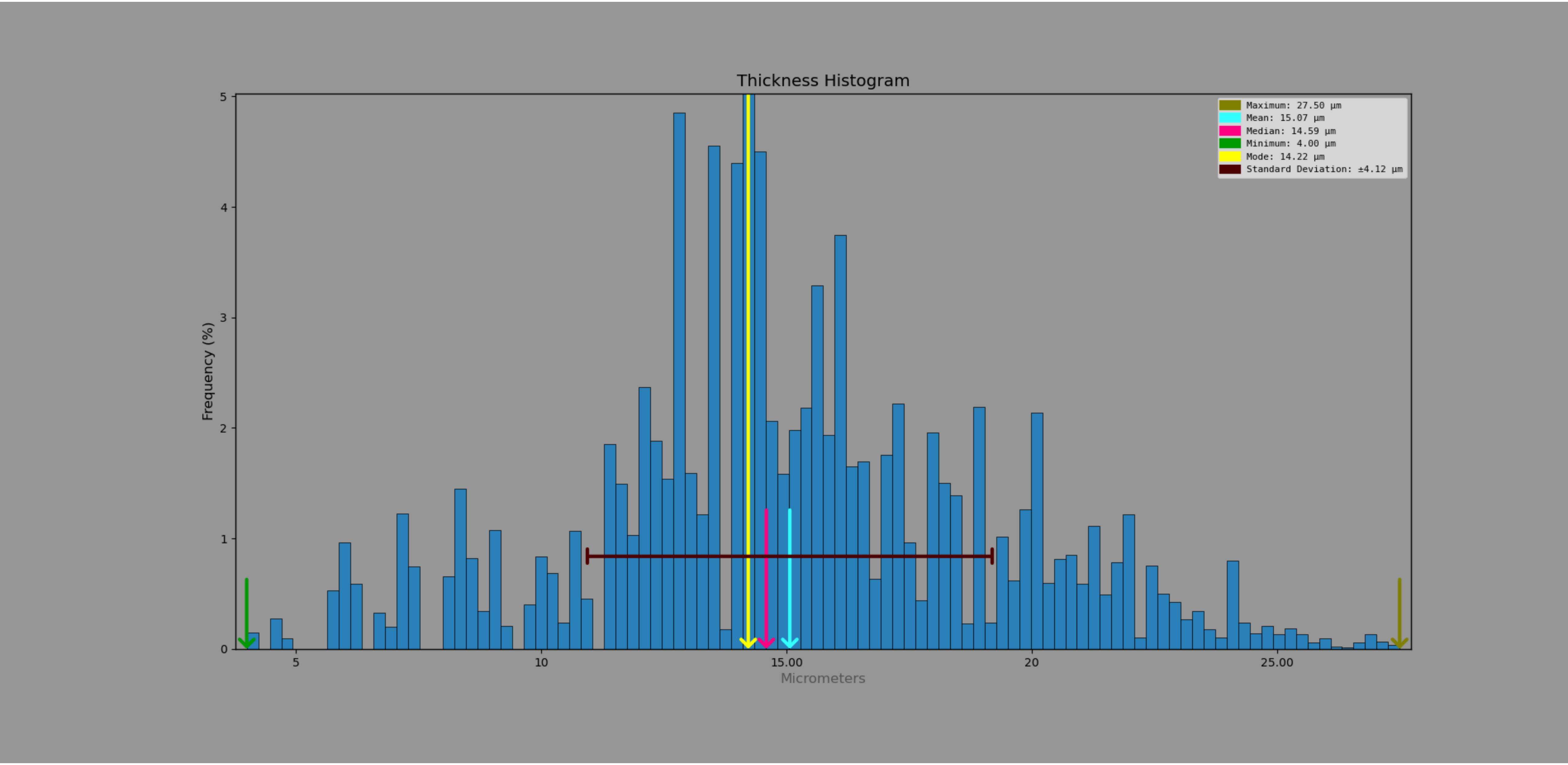

*Elimaea signata*

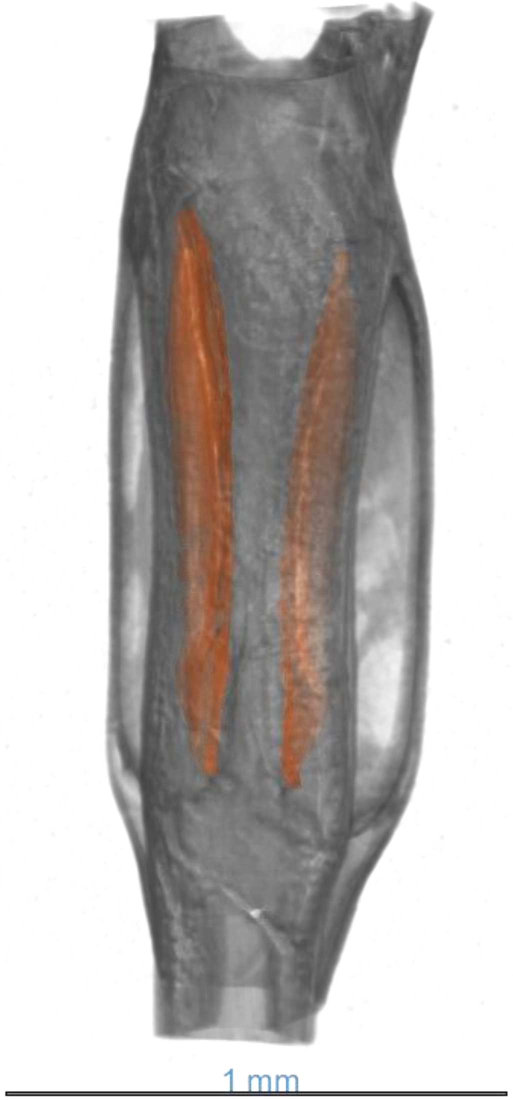


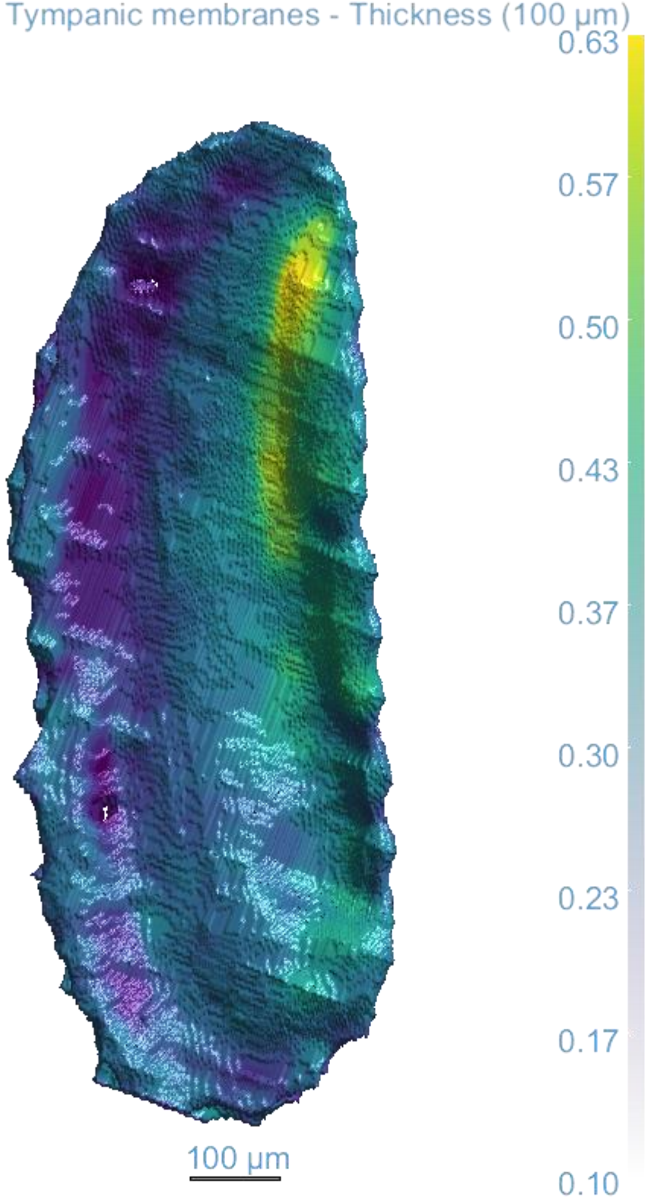

*Elimaea signata*

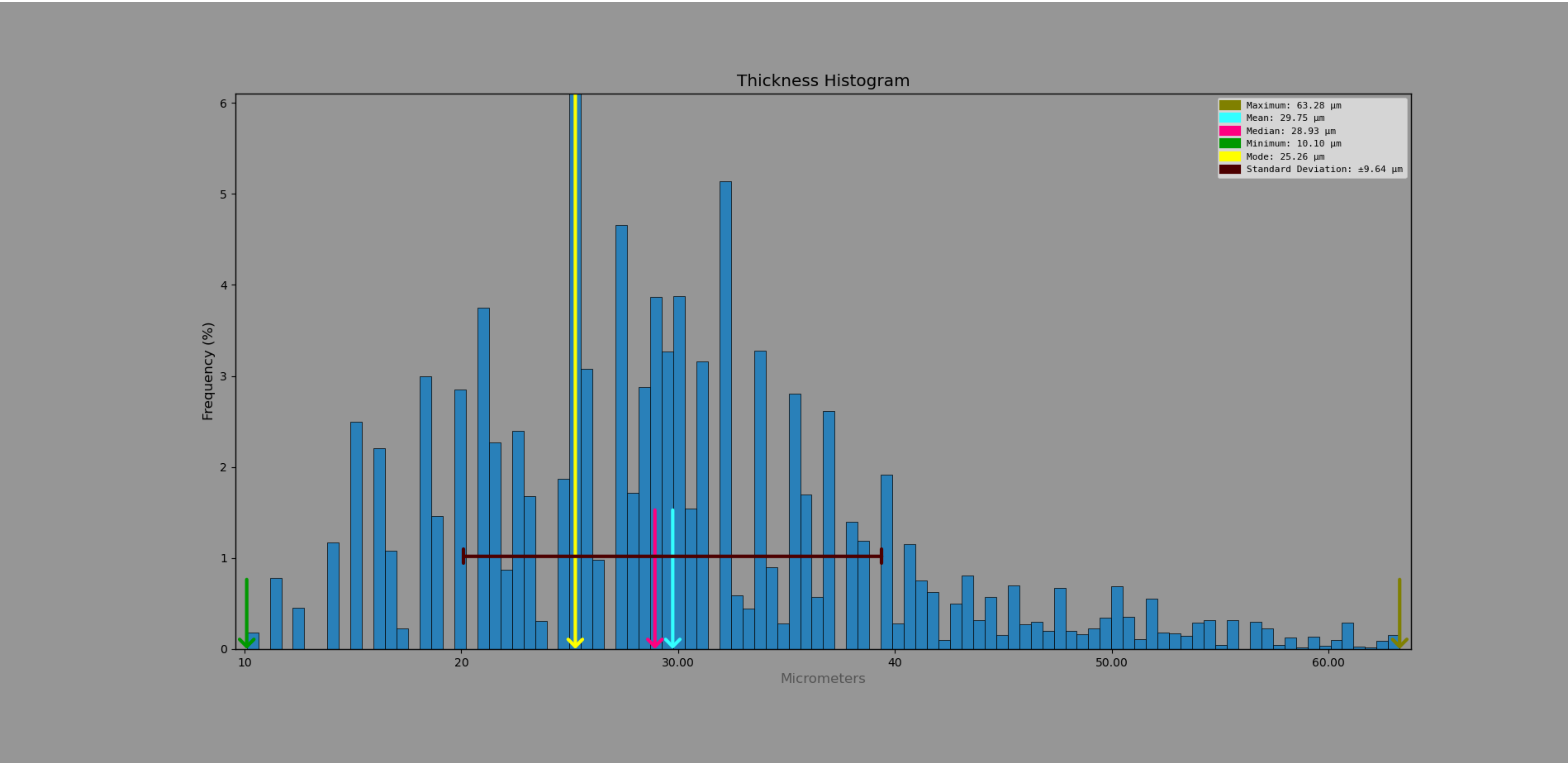

*Haenschiella sp.*

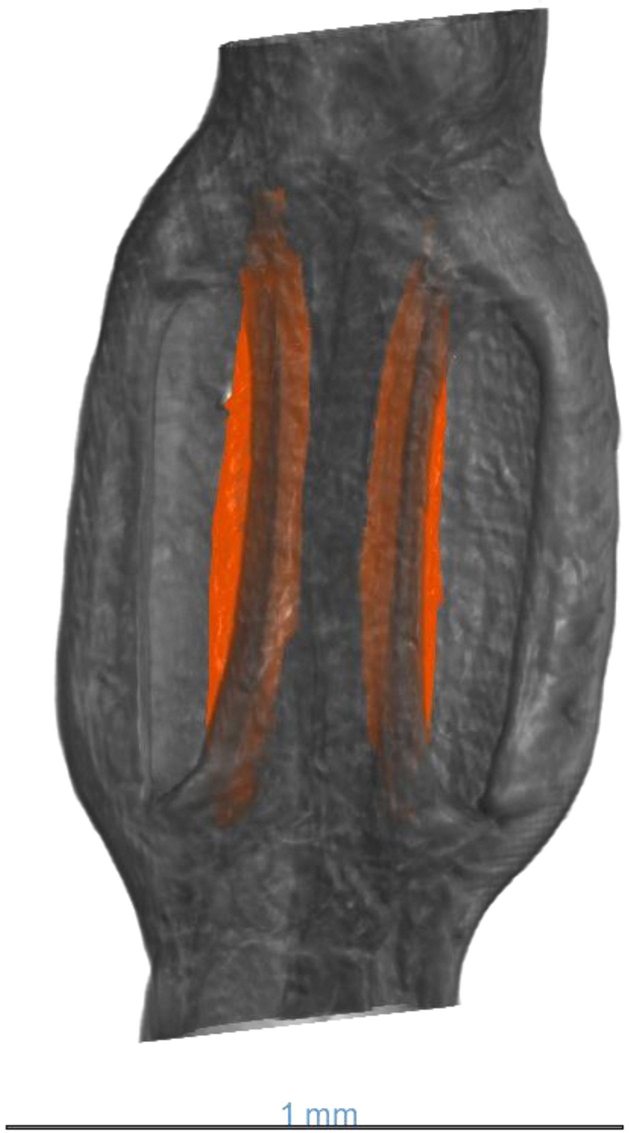


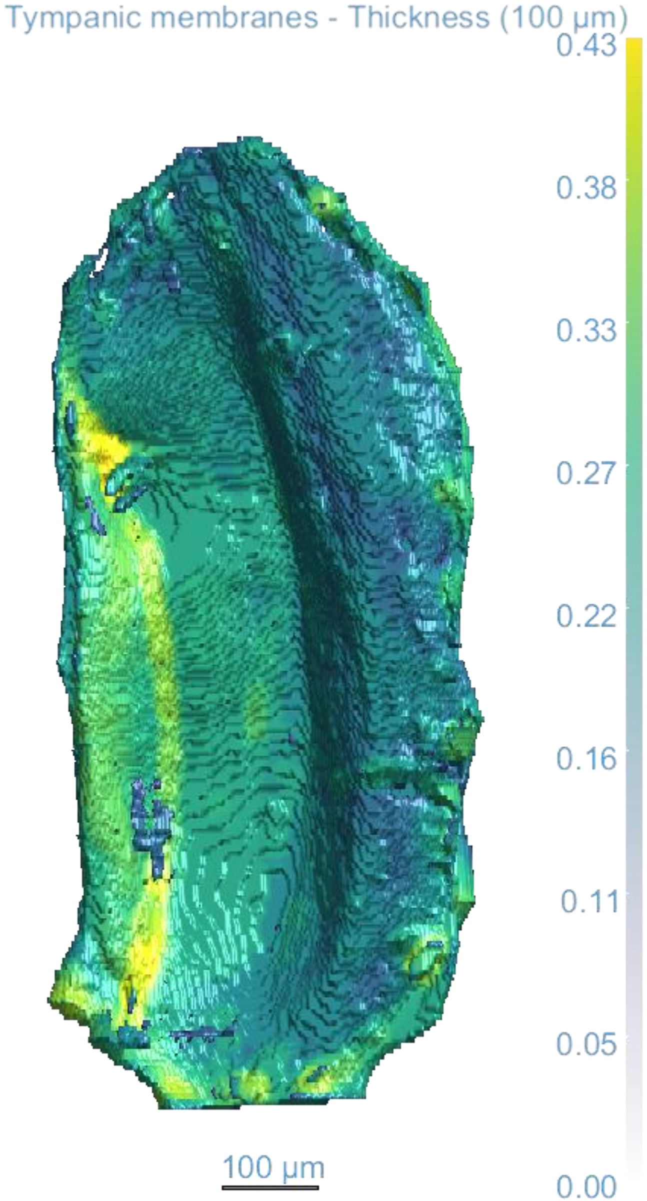

*Haenschiella sp.*

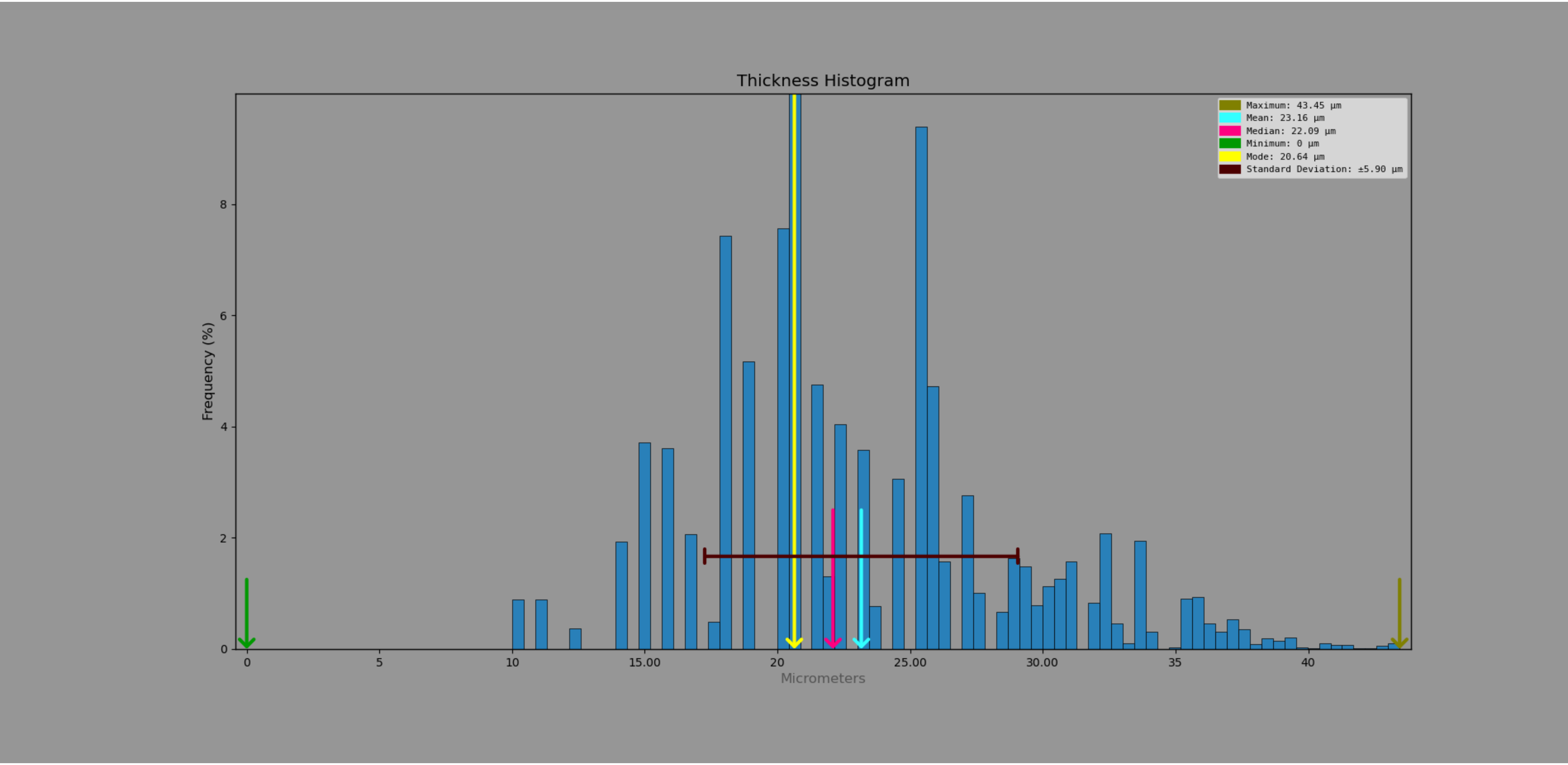

*Leptoderes ornatipennis*

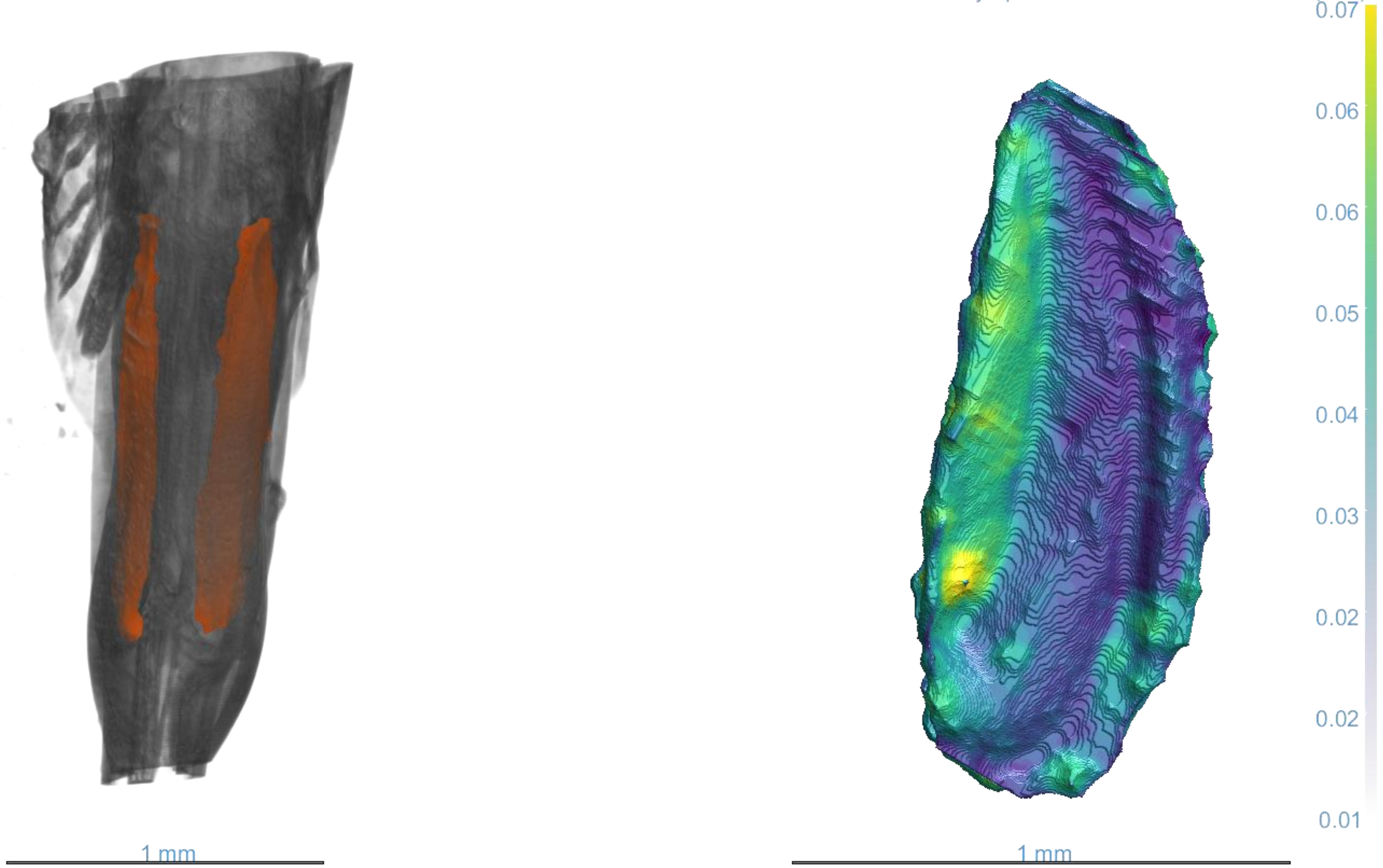

*Leptoderes ornatipennis*

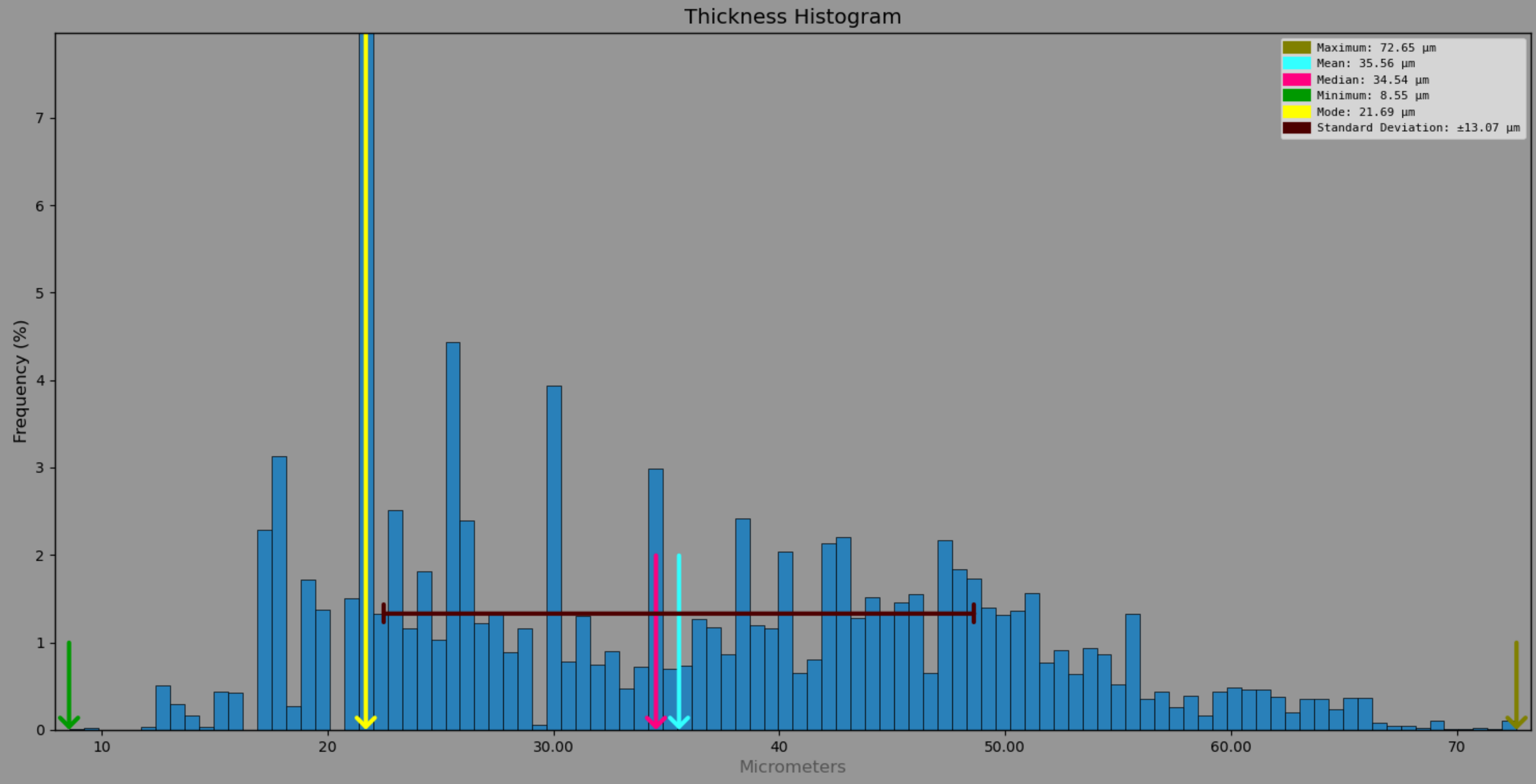

*Mecopoda elongata*

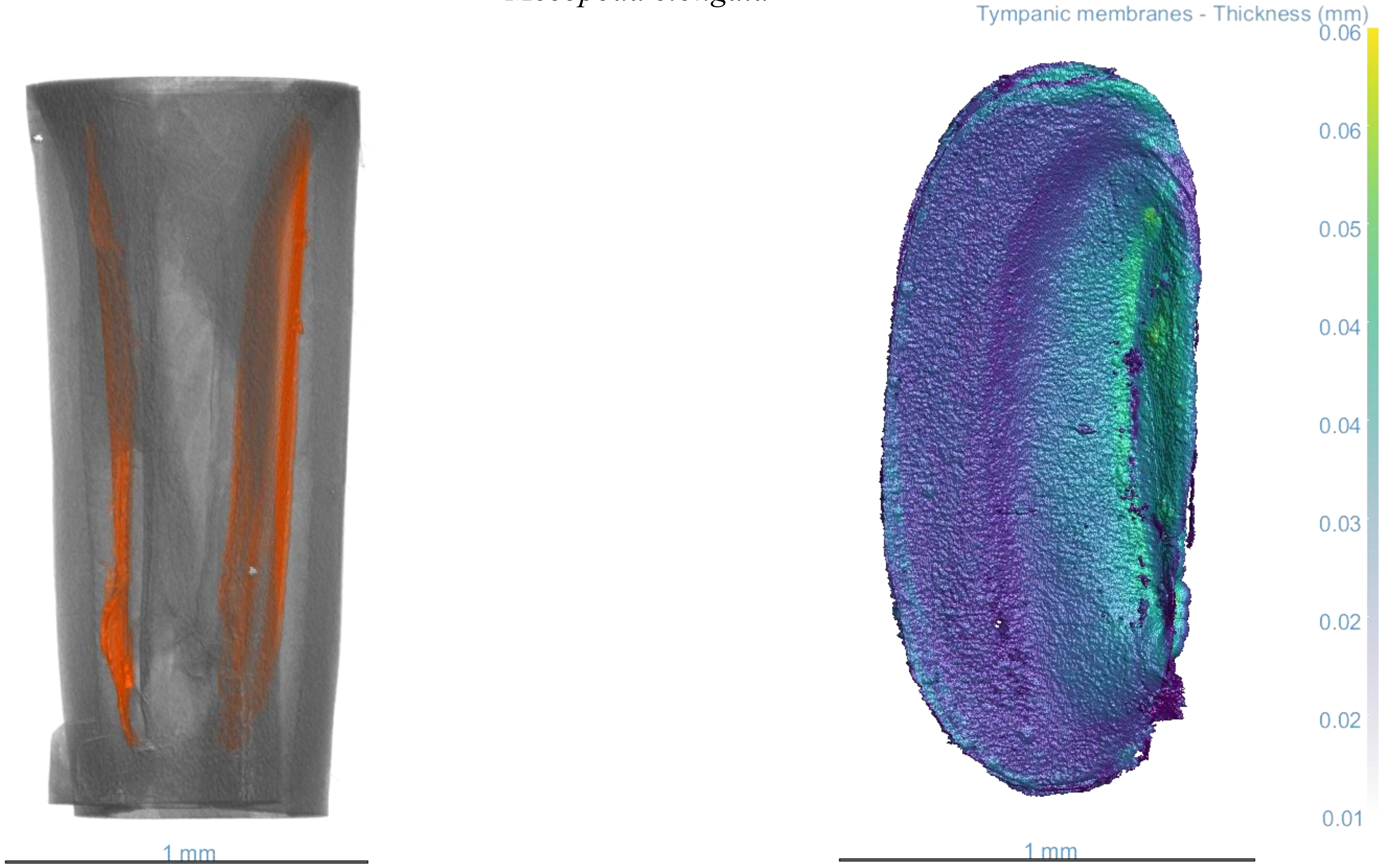

*Mecopoda elongata*

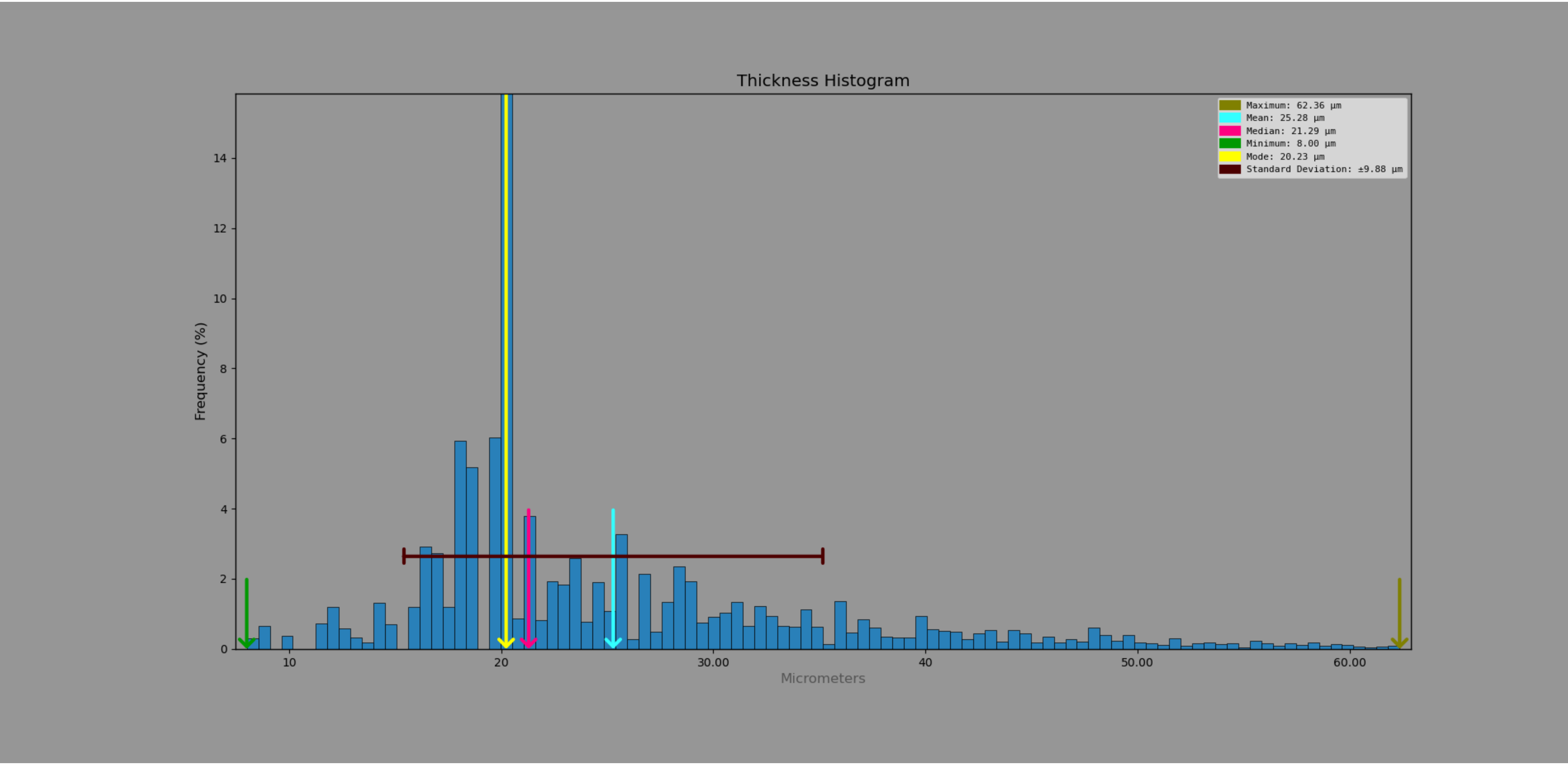

*Moncheca elegans*

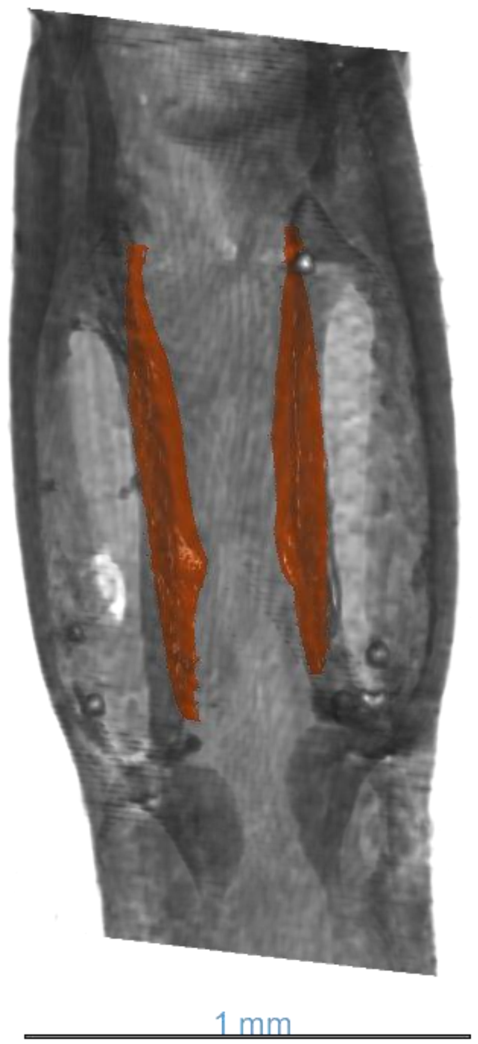


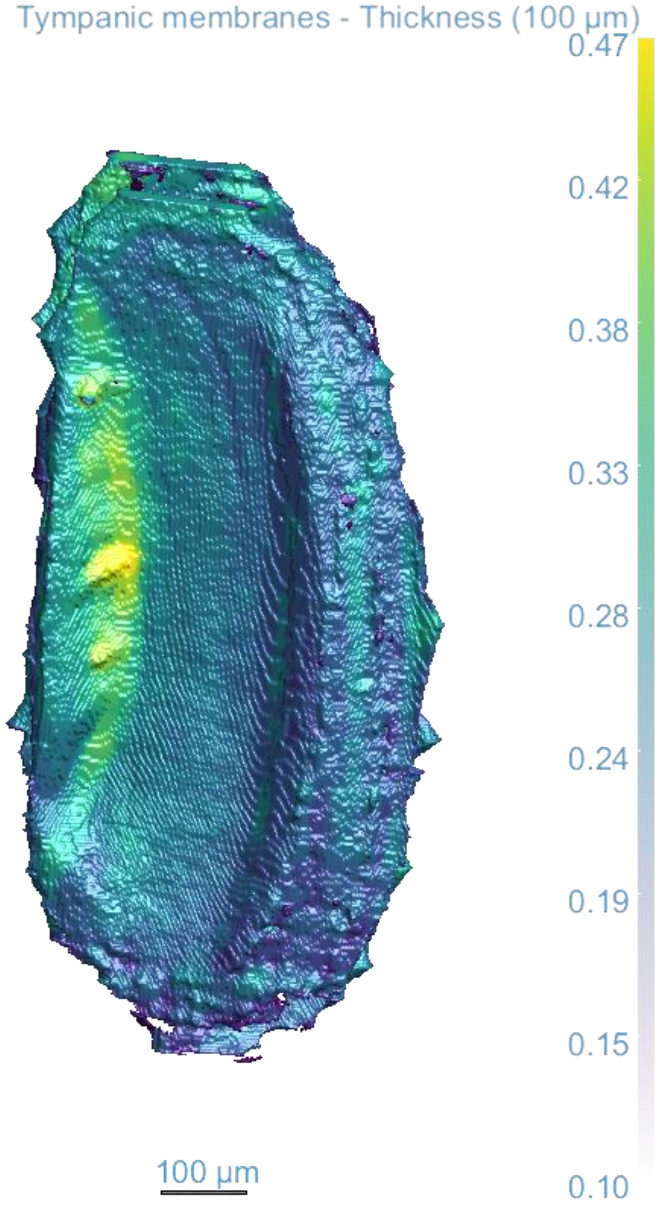

*Moncheca elegans*

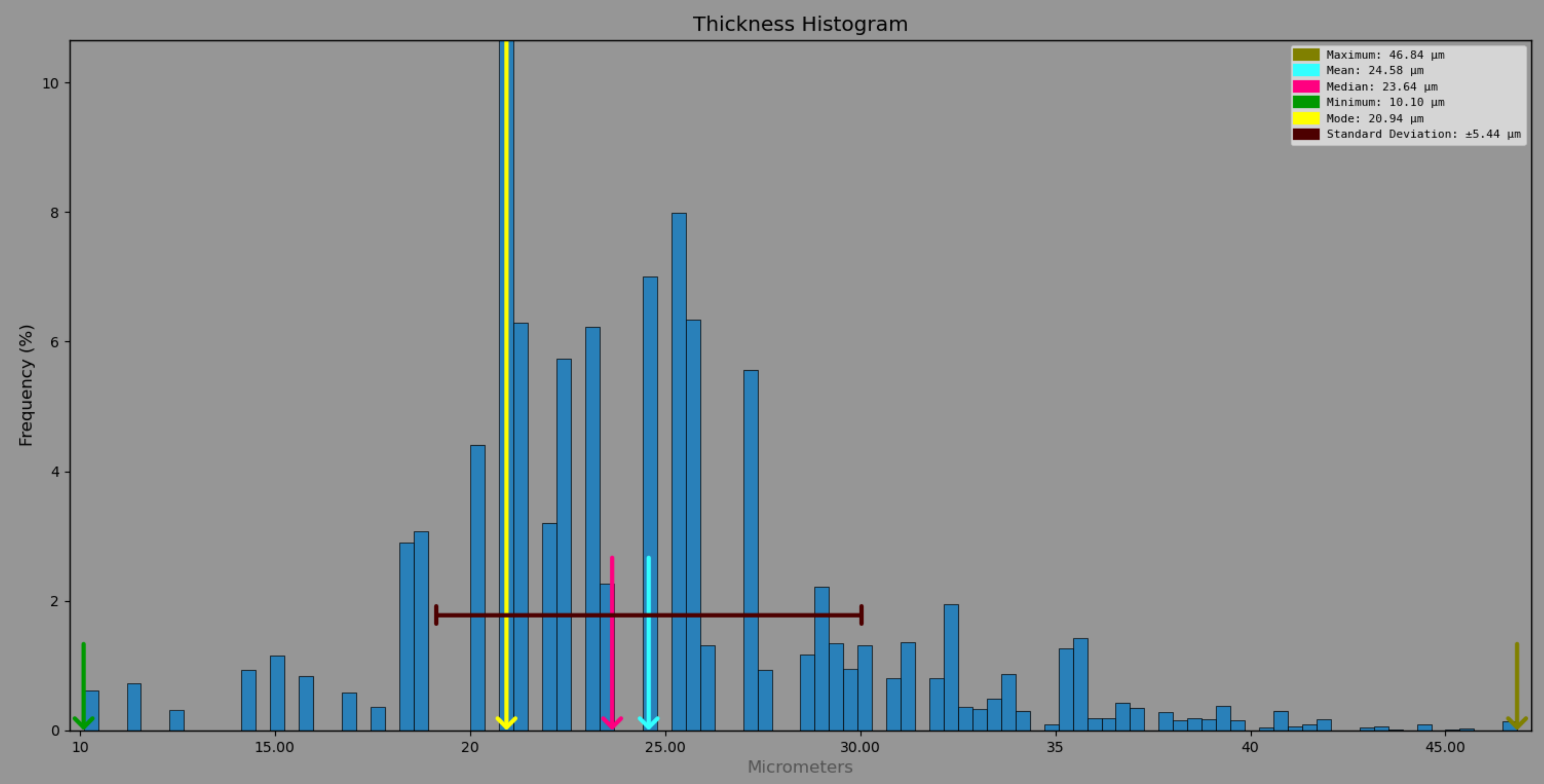

*Phaulula galeata*

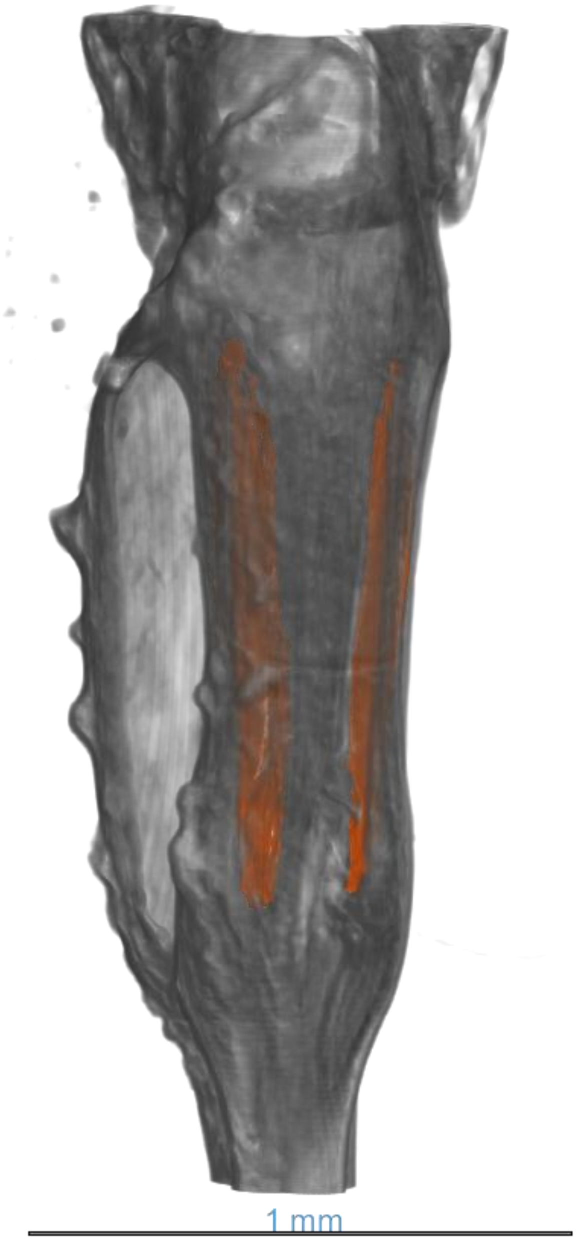


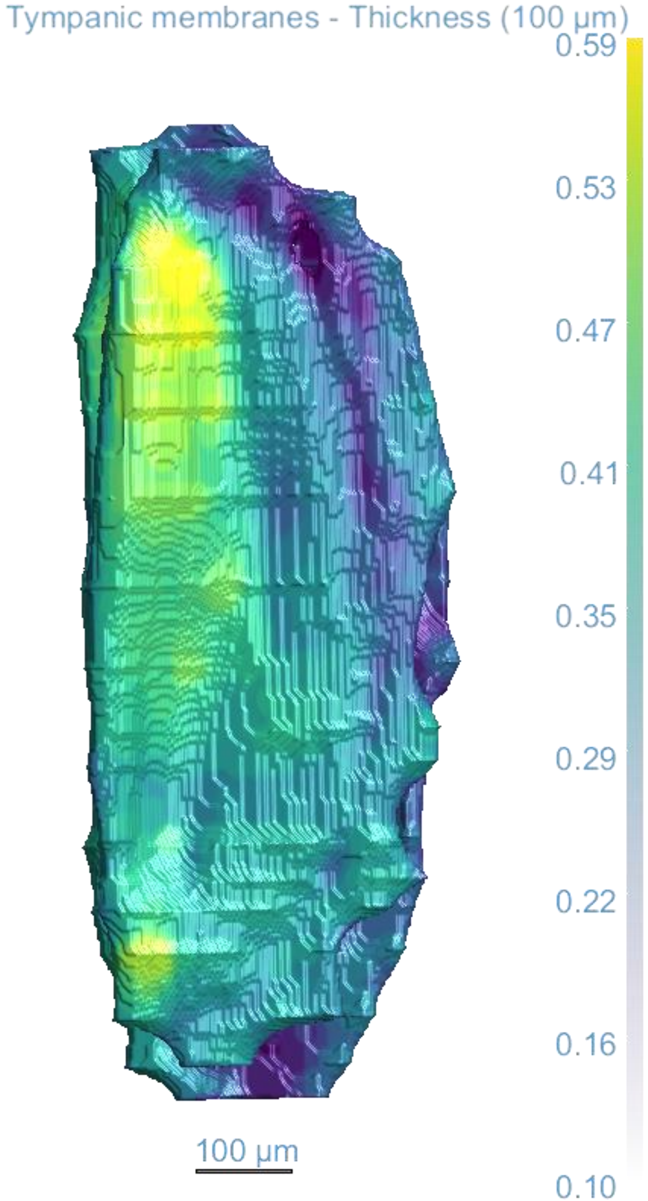

*Phaulula galeata*

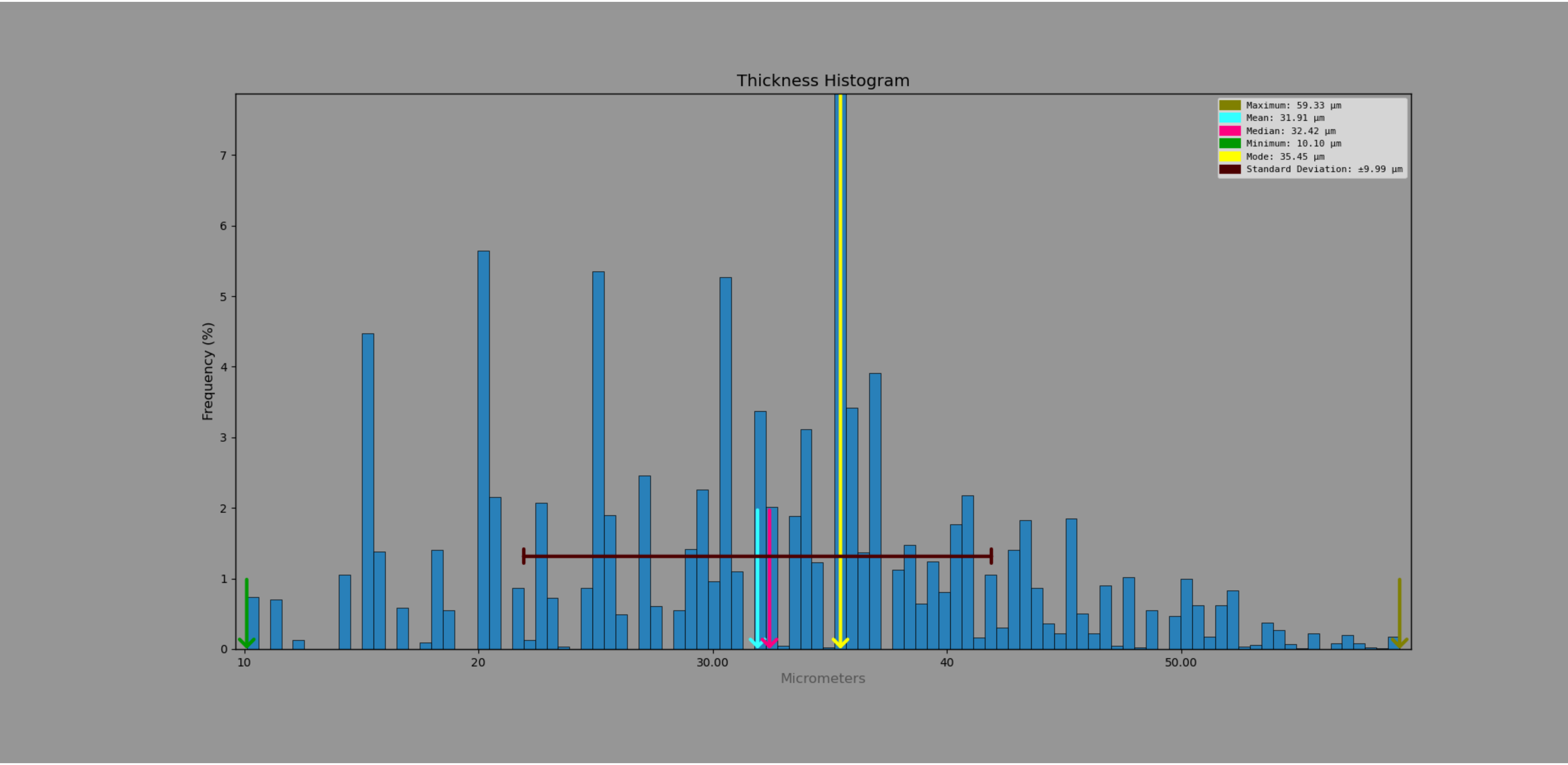

*Phaulula galeata* – exposed tympana

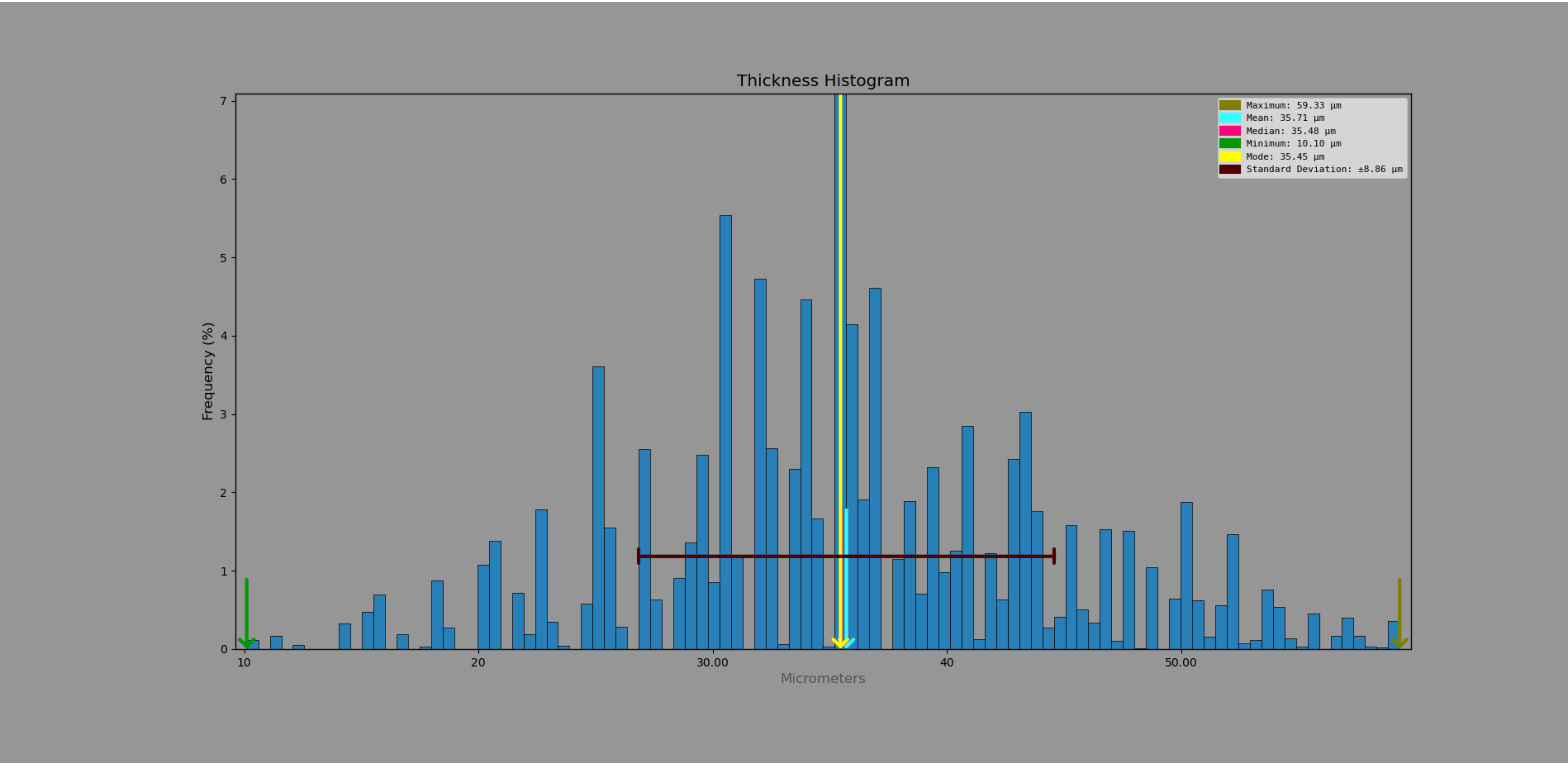

*Phaulula galeata* – pinna covered tympana

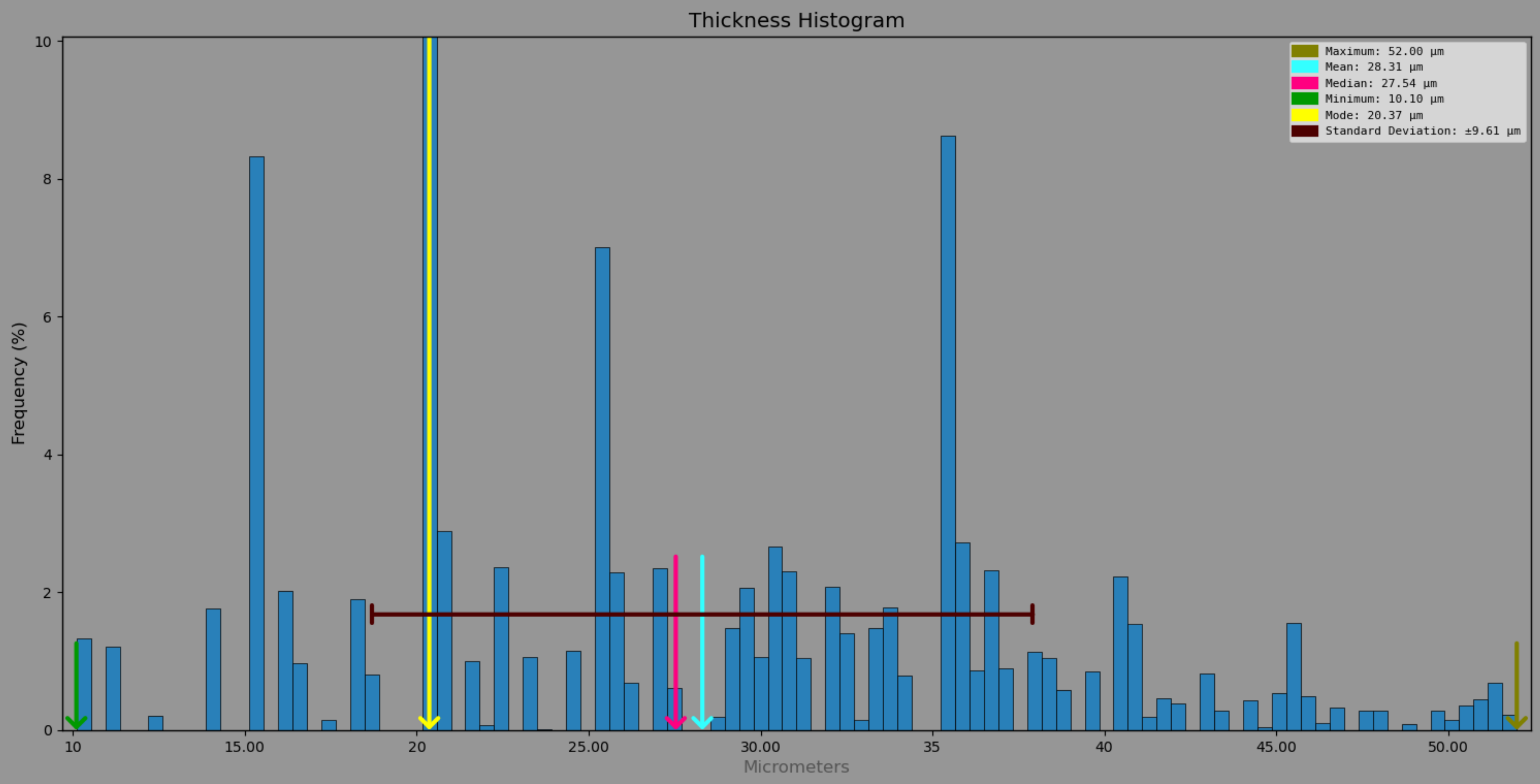

*Phlugis poecilla*

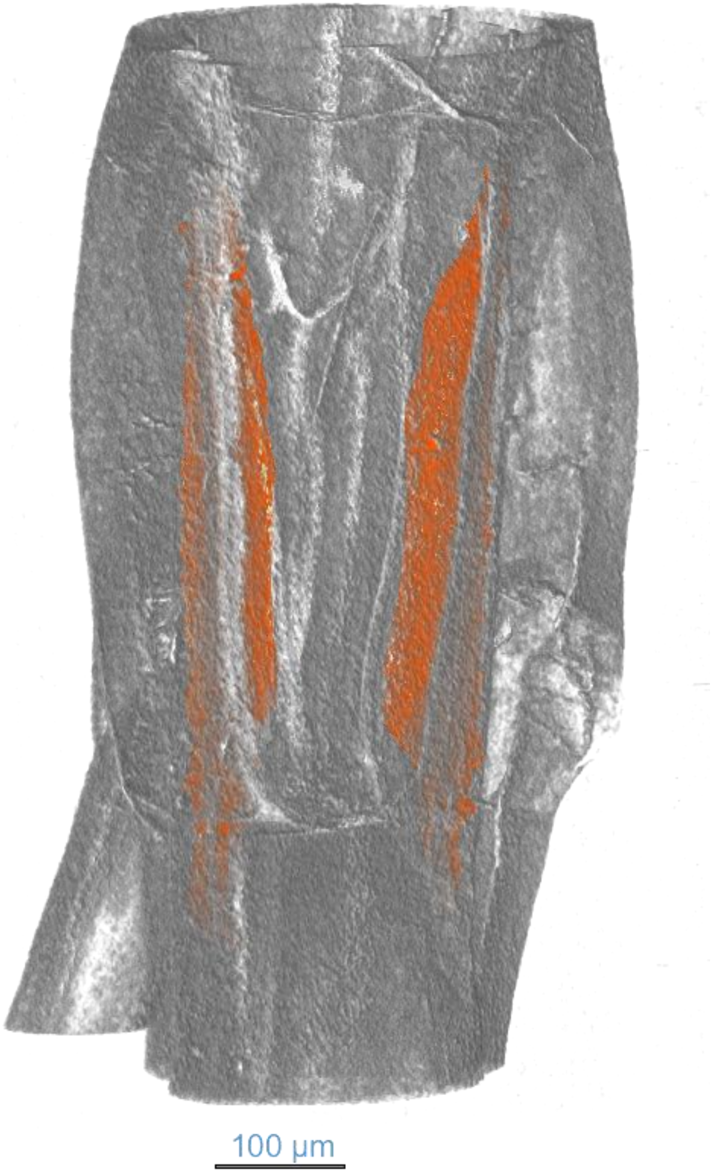


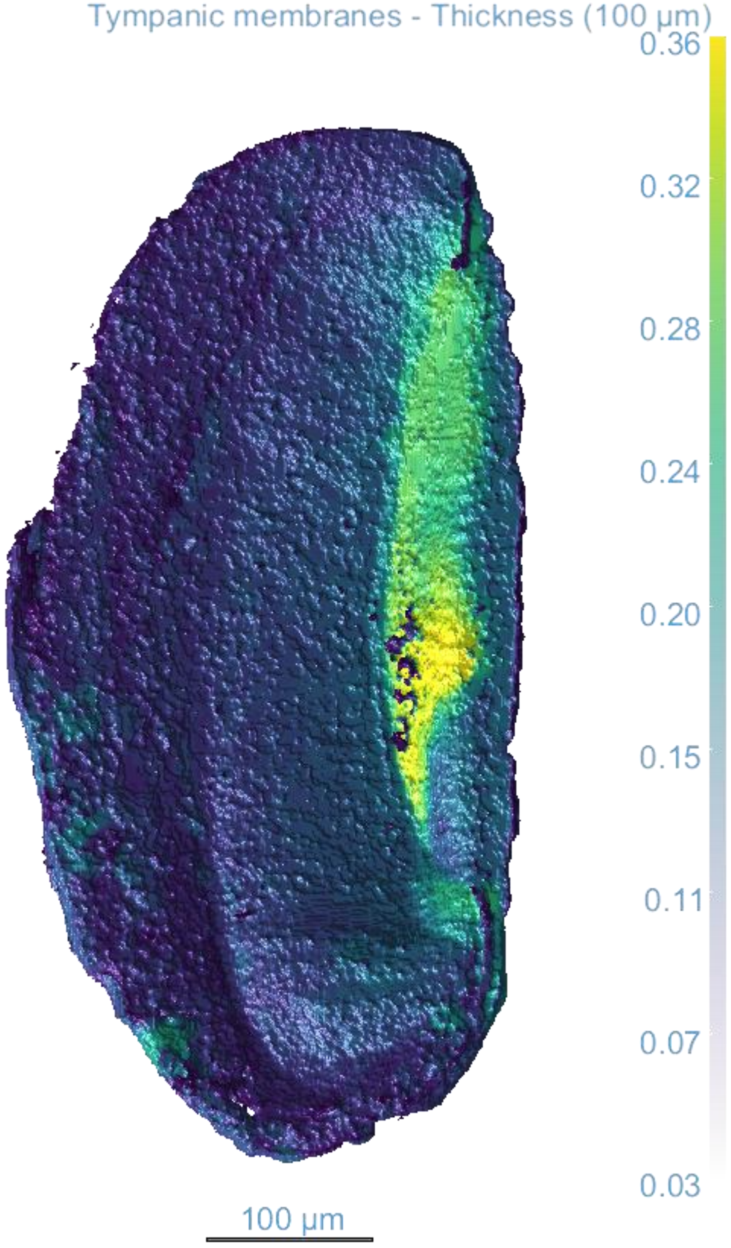

*Phlugis poecilla*

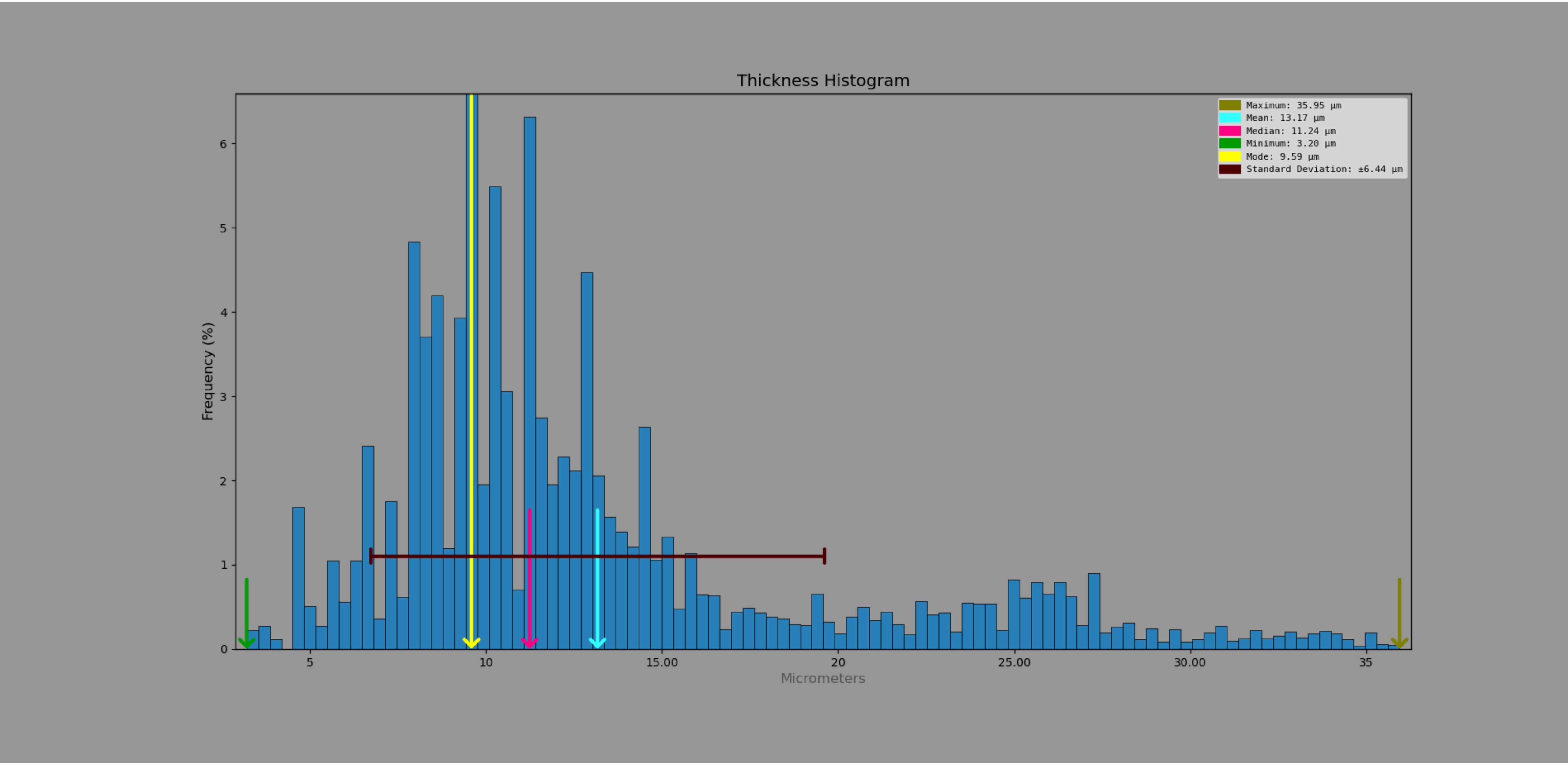

*Phygela marginata*

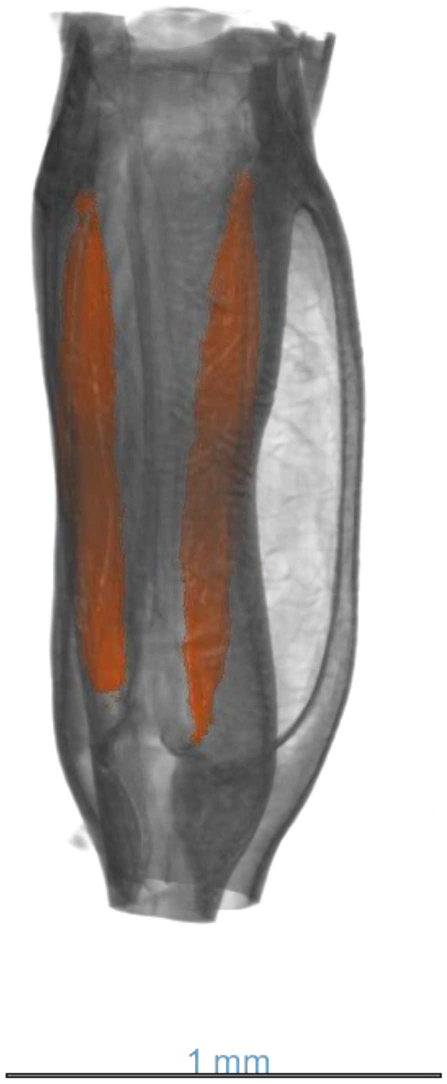


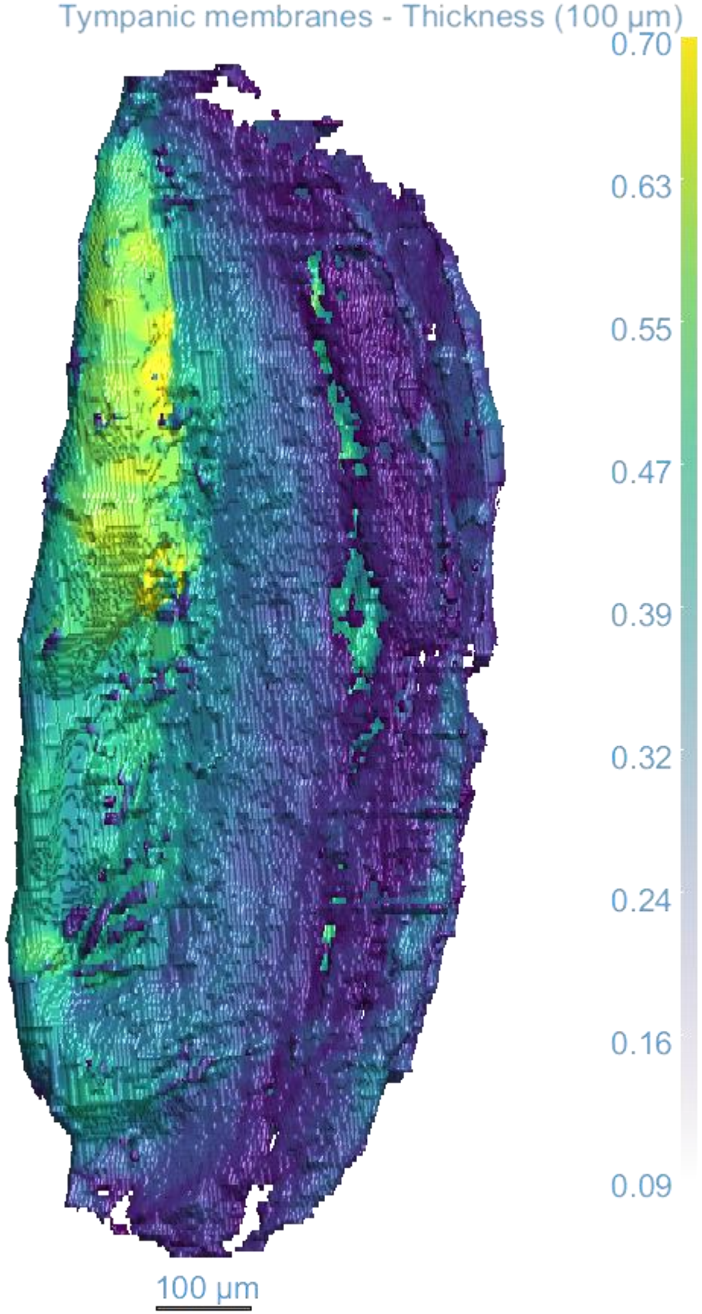

*Phygela marginata*

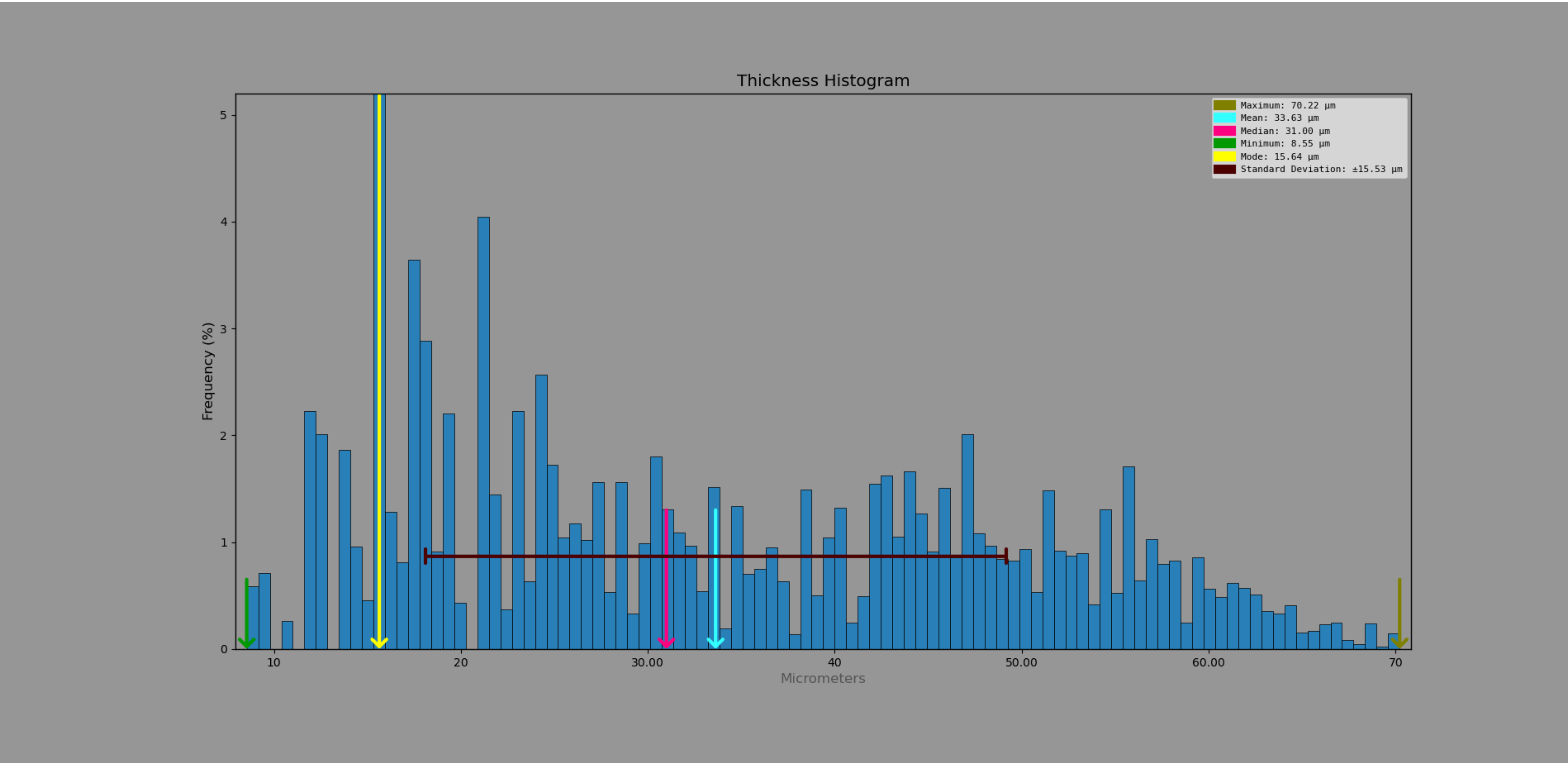

*Phygela marginata* - exposed tympana

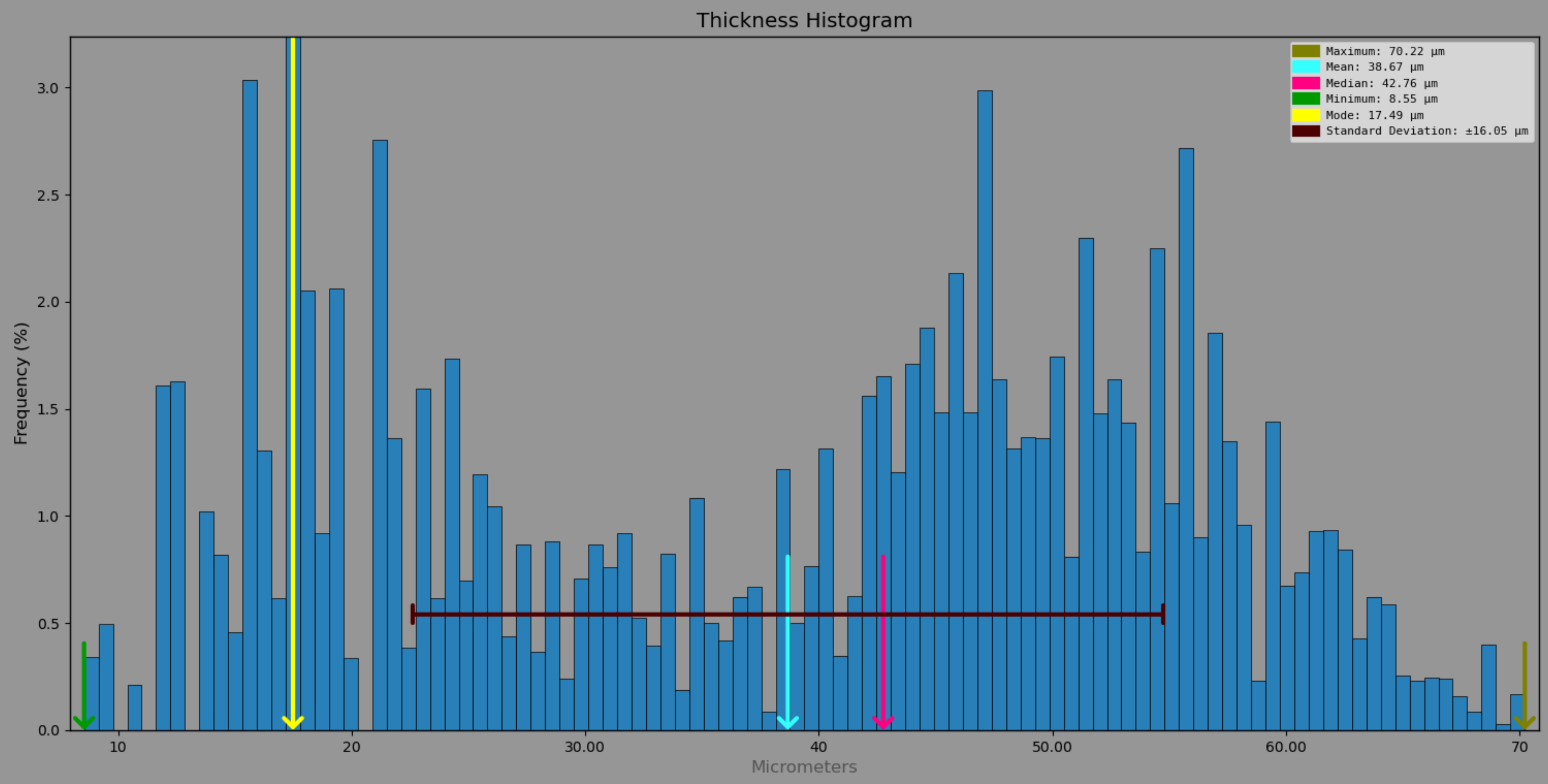

*Phygela marginata* – pinna covered tympana

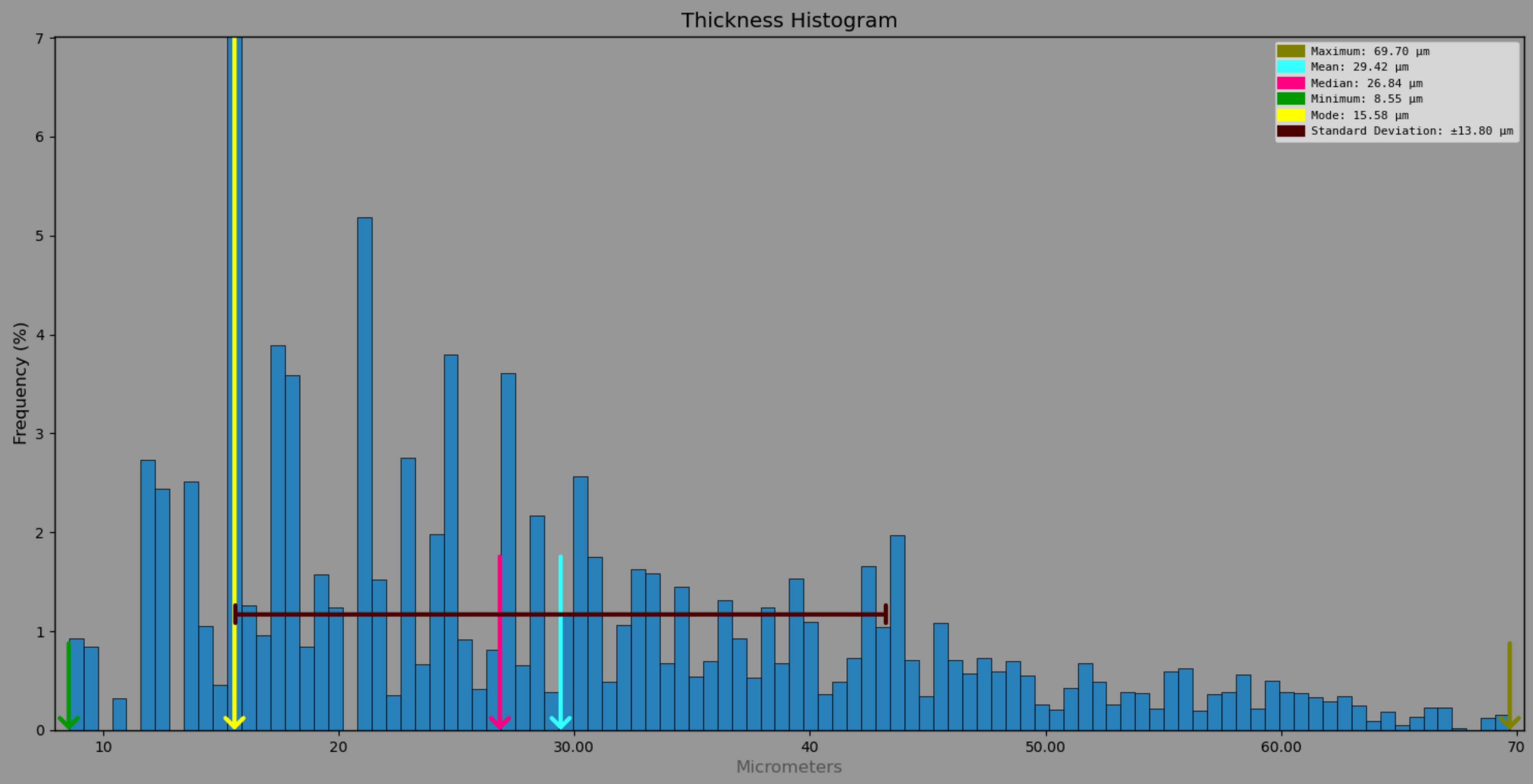

*Phyllomimus detersus*

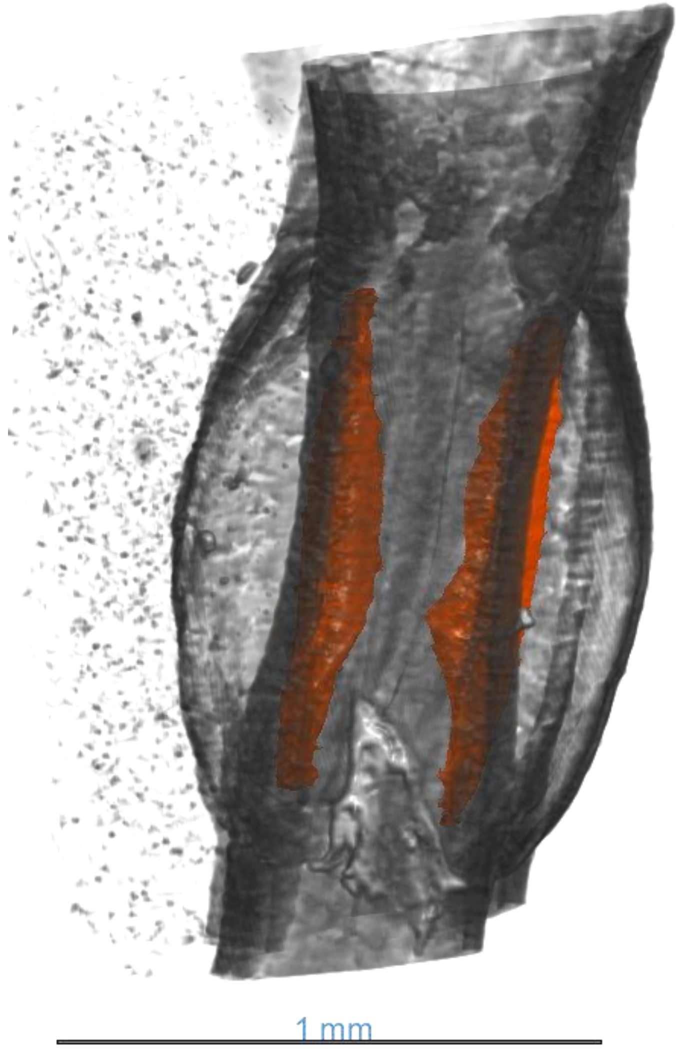


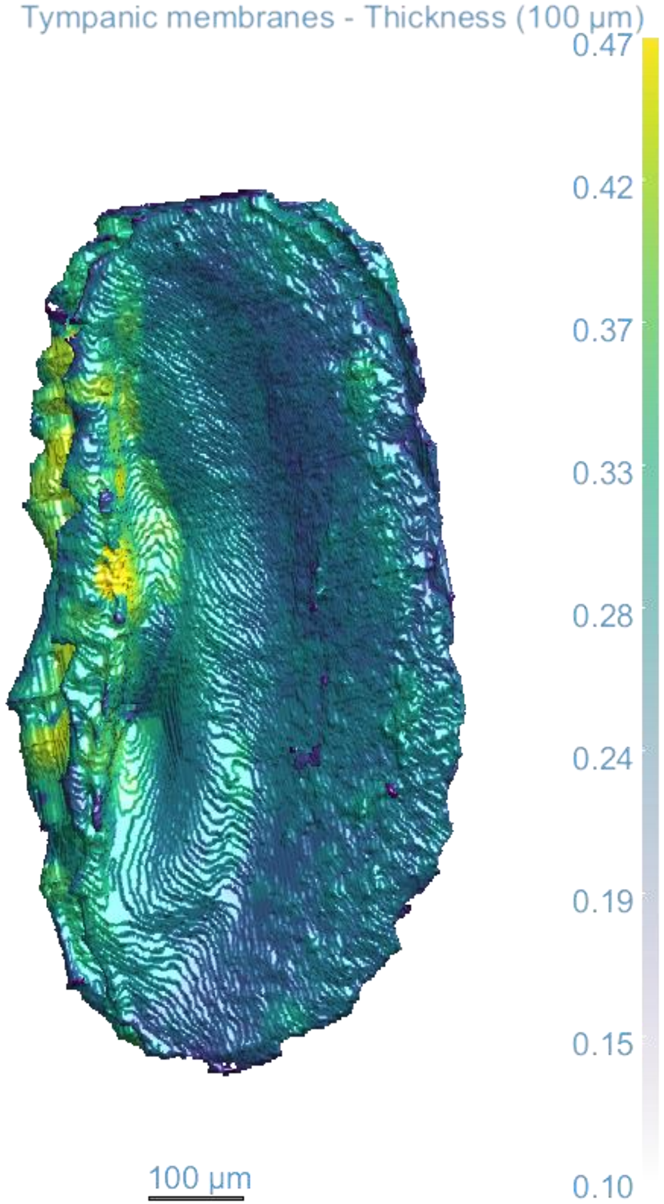

*Phyllomimus detersus*

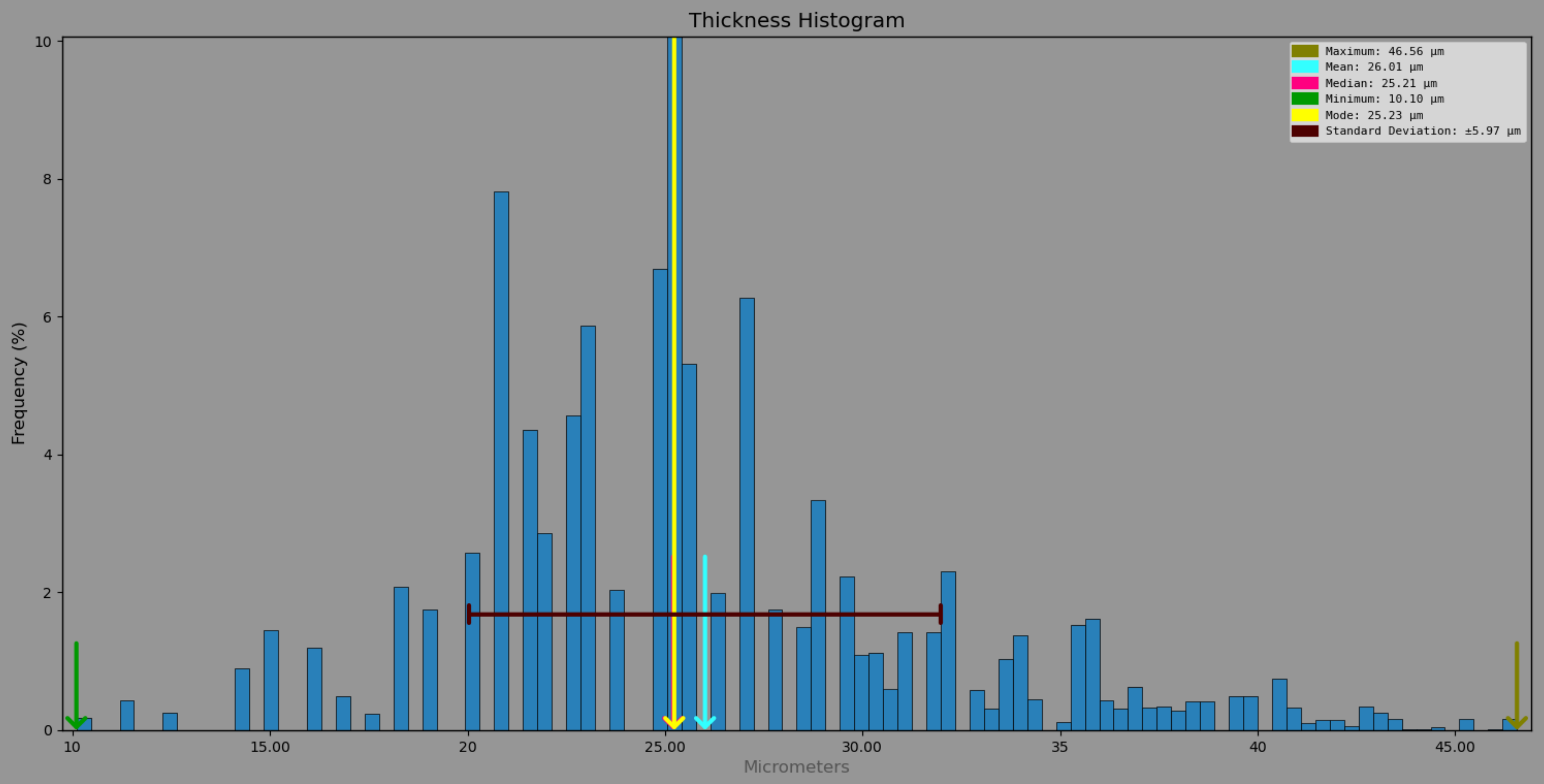

*Ragoniella pulchella*

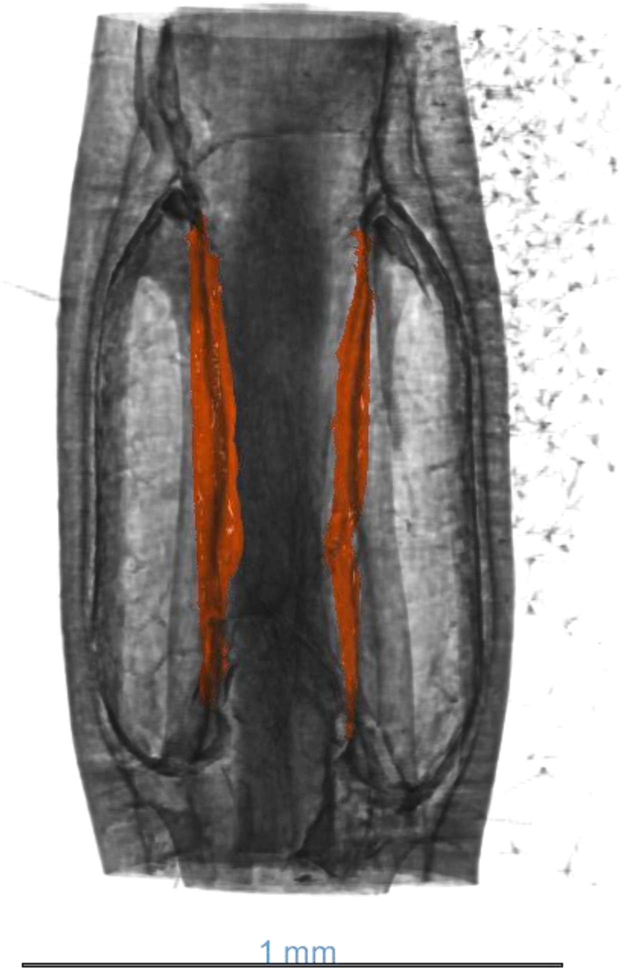


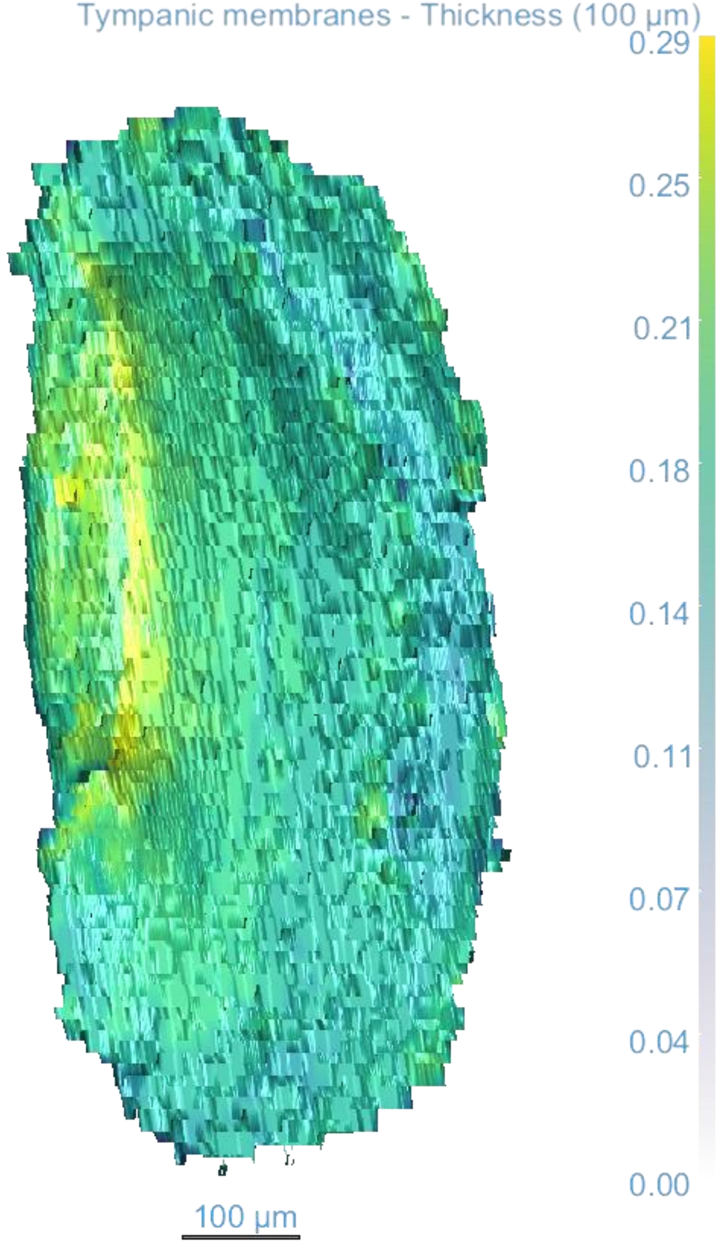

*Ragoniella pulchella*

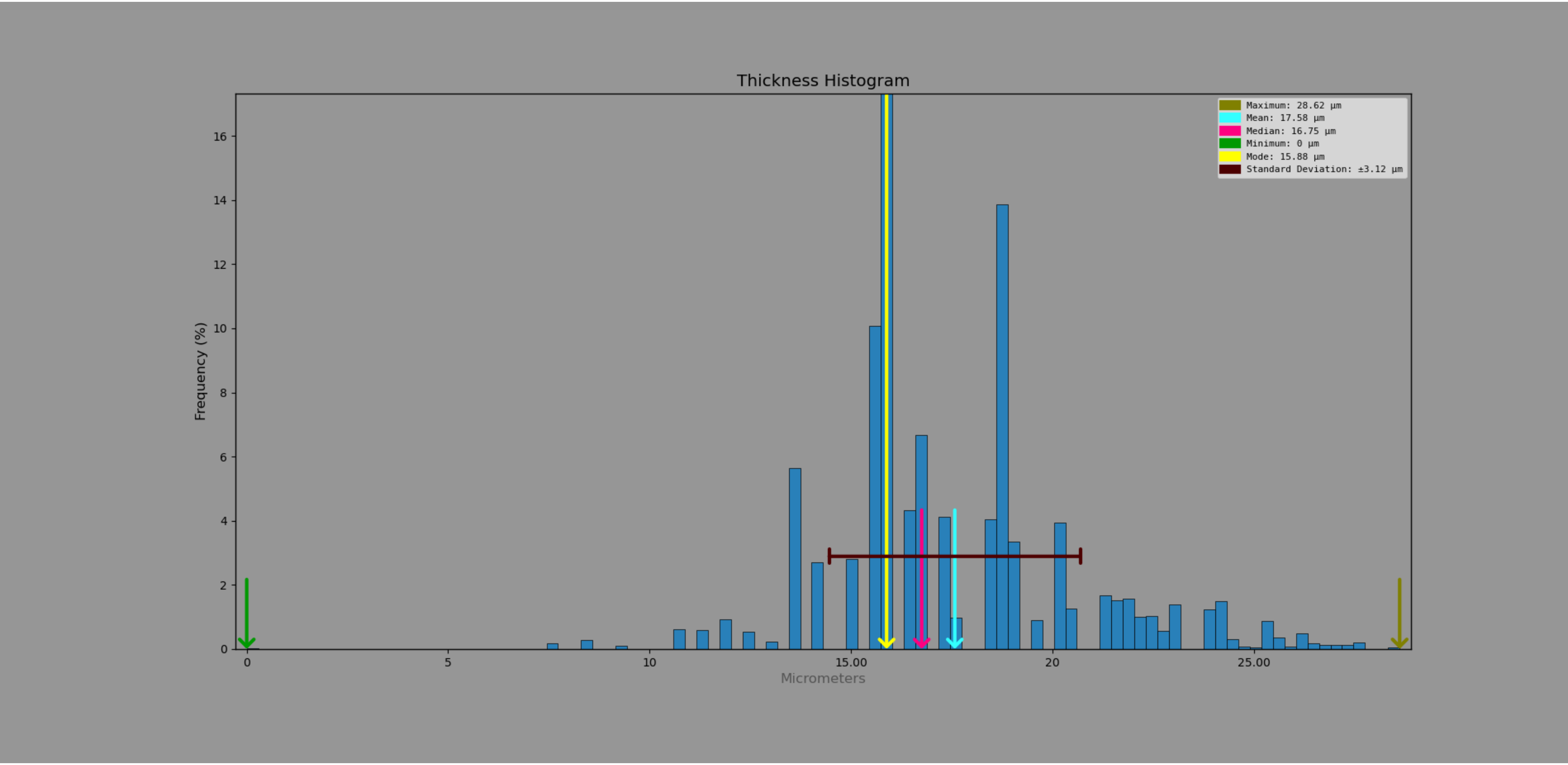

*Satizabalus jorgevargasi*

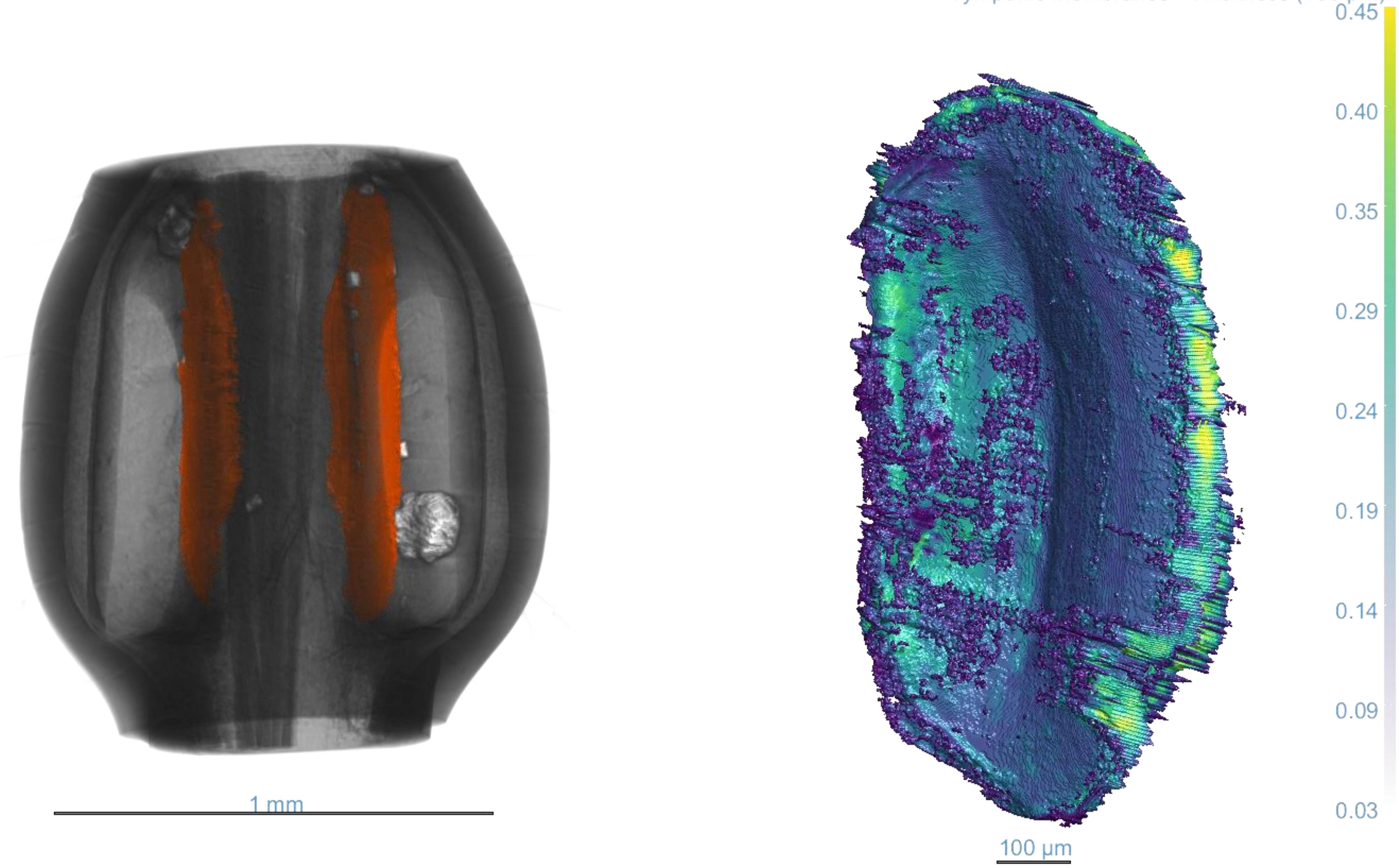

*Satizabalus jorgevargasi*

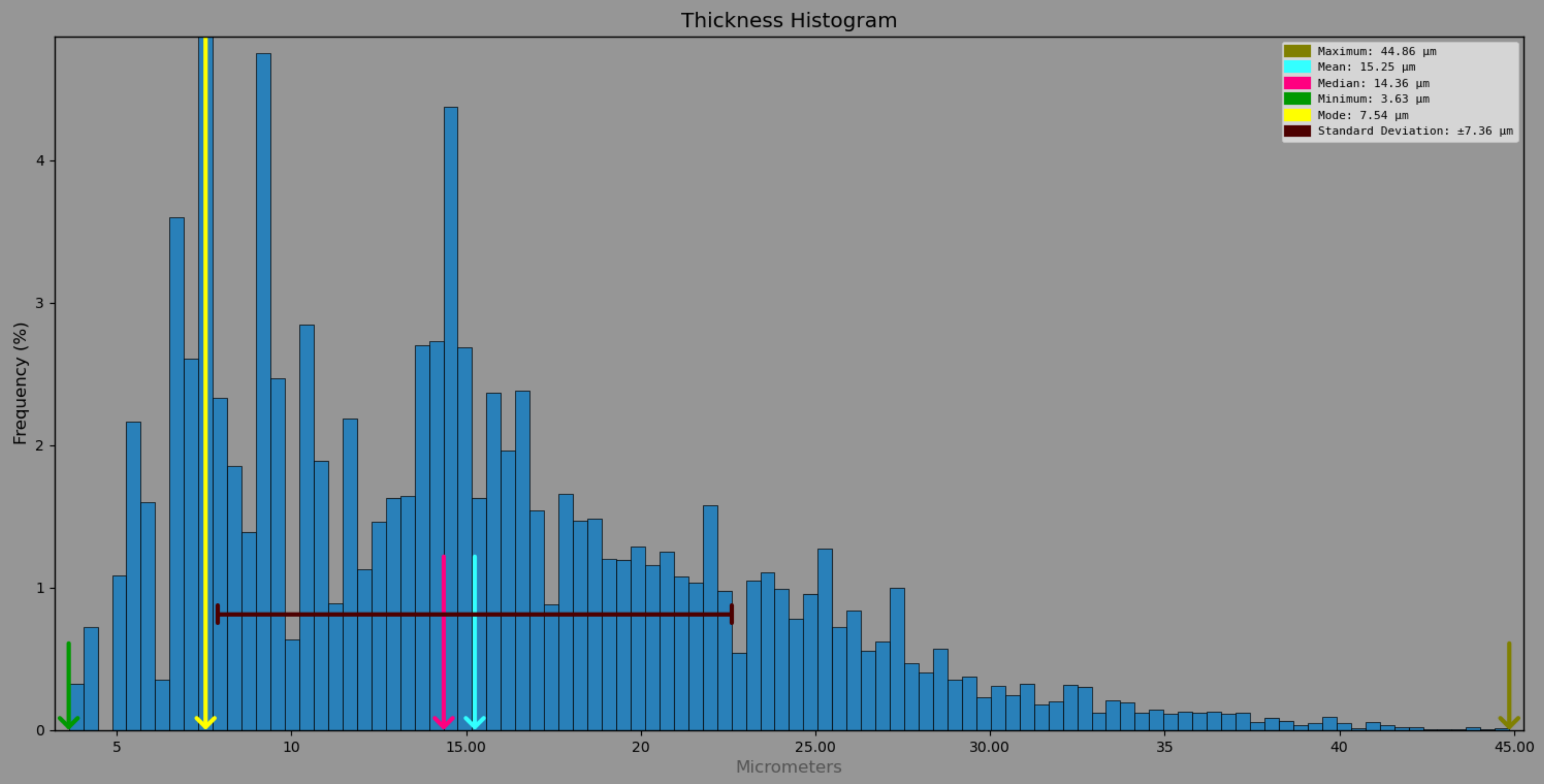

*Stictophaula sp.*

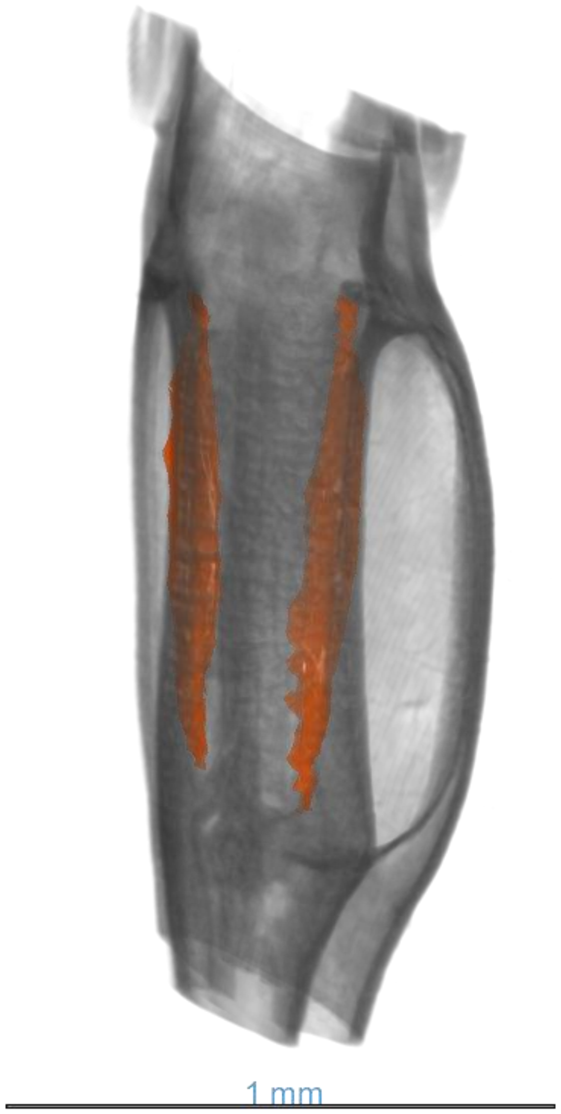


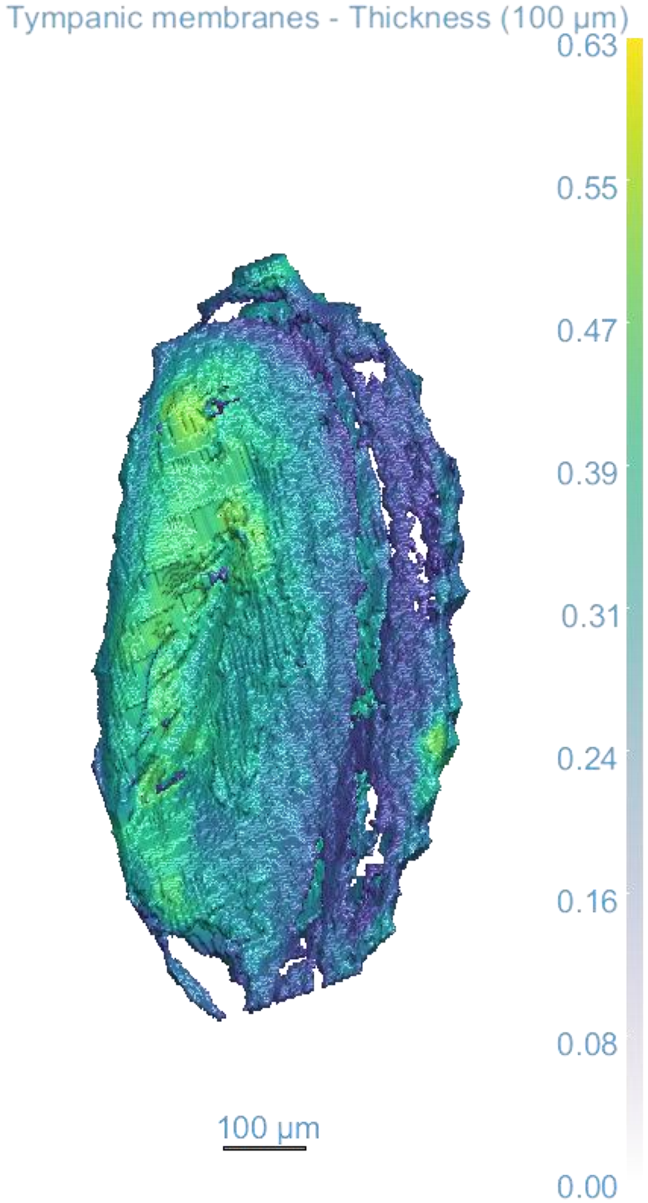

*Stictophaula sp.*

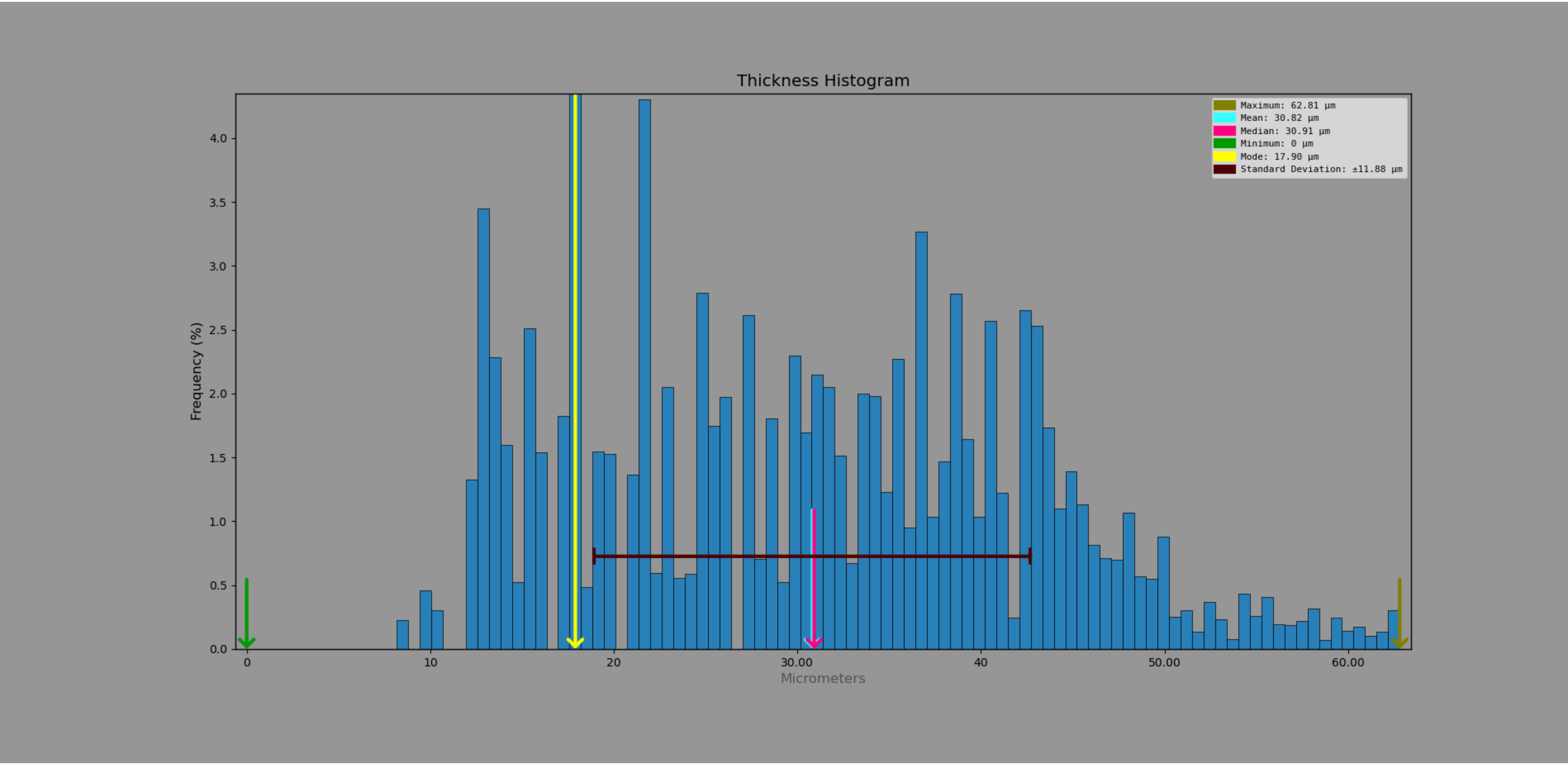

*Stictophaula sp.* - exposed tympana

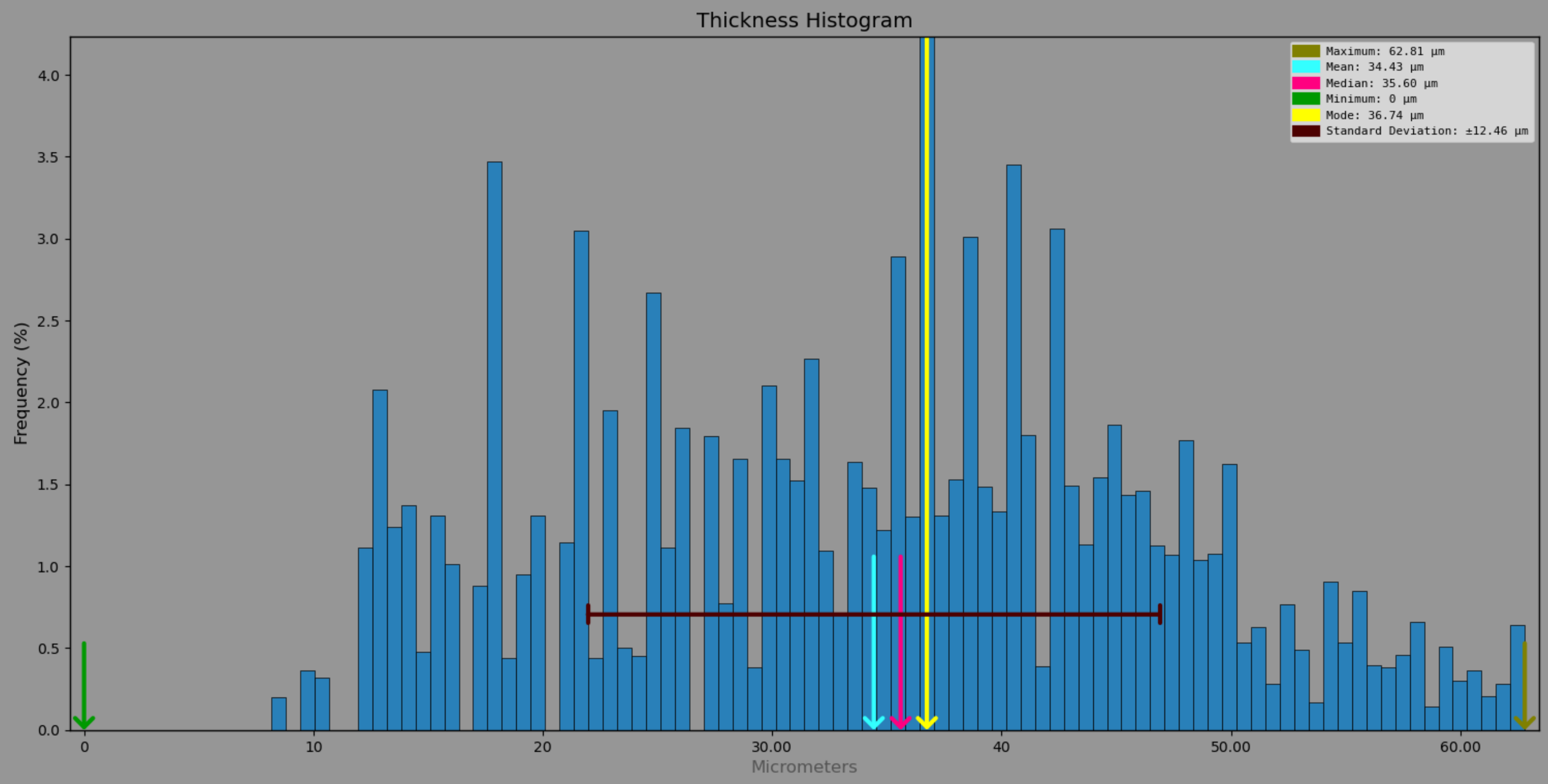

*Stictophaula sp.* – pinna covered tympana

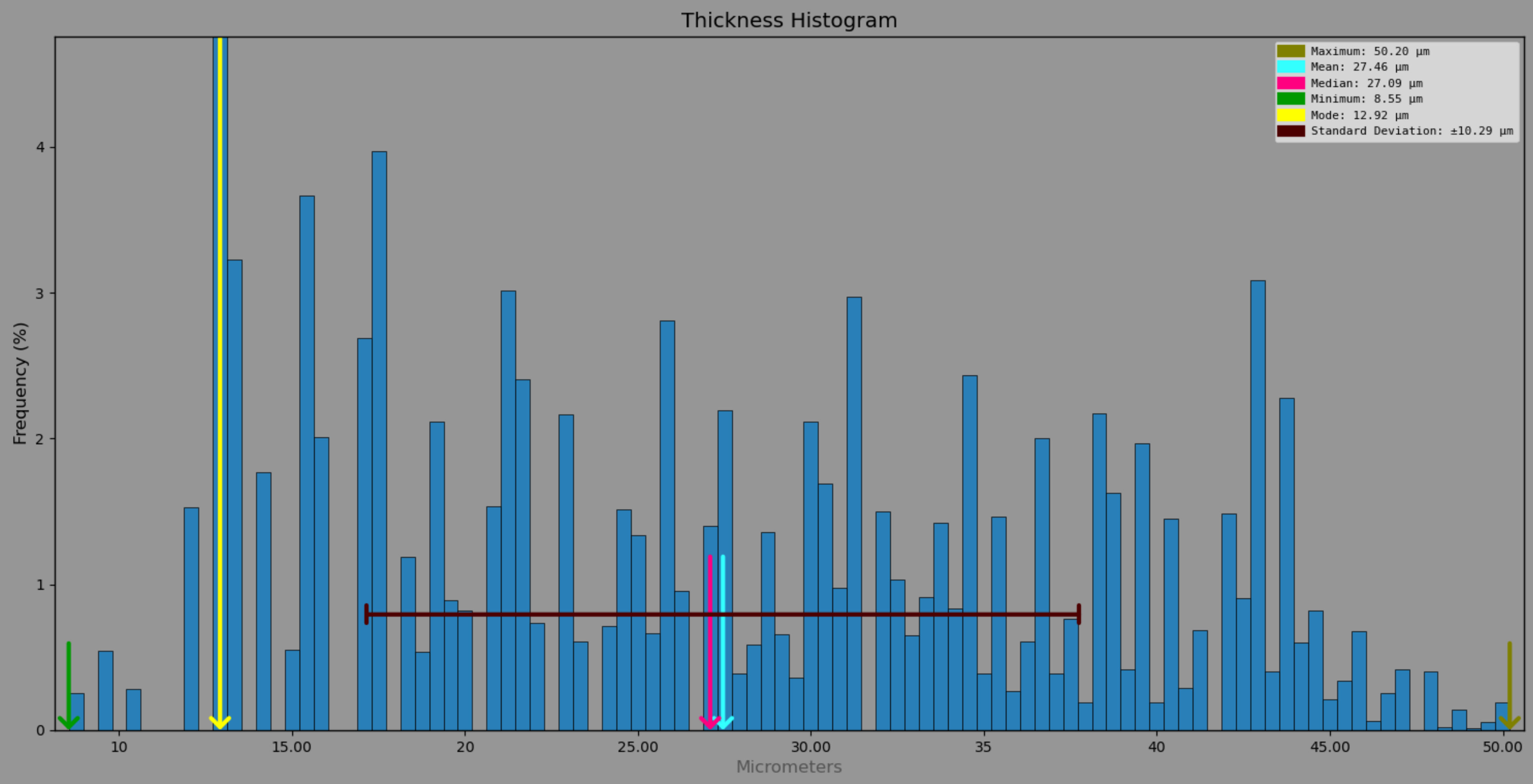

*Stilpnochlora sp.*

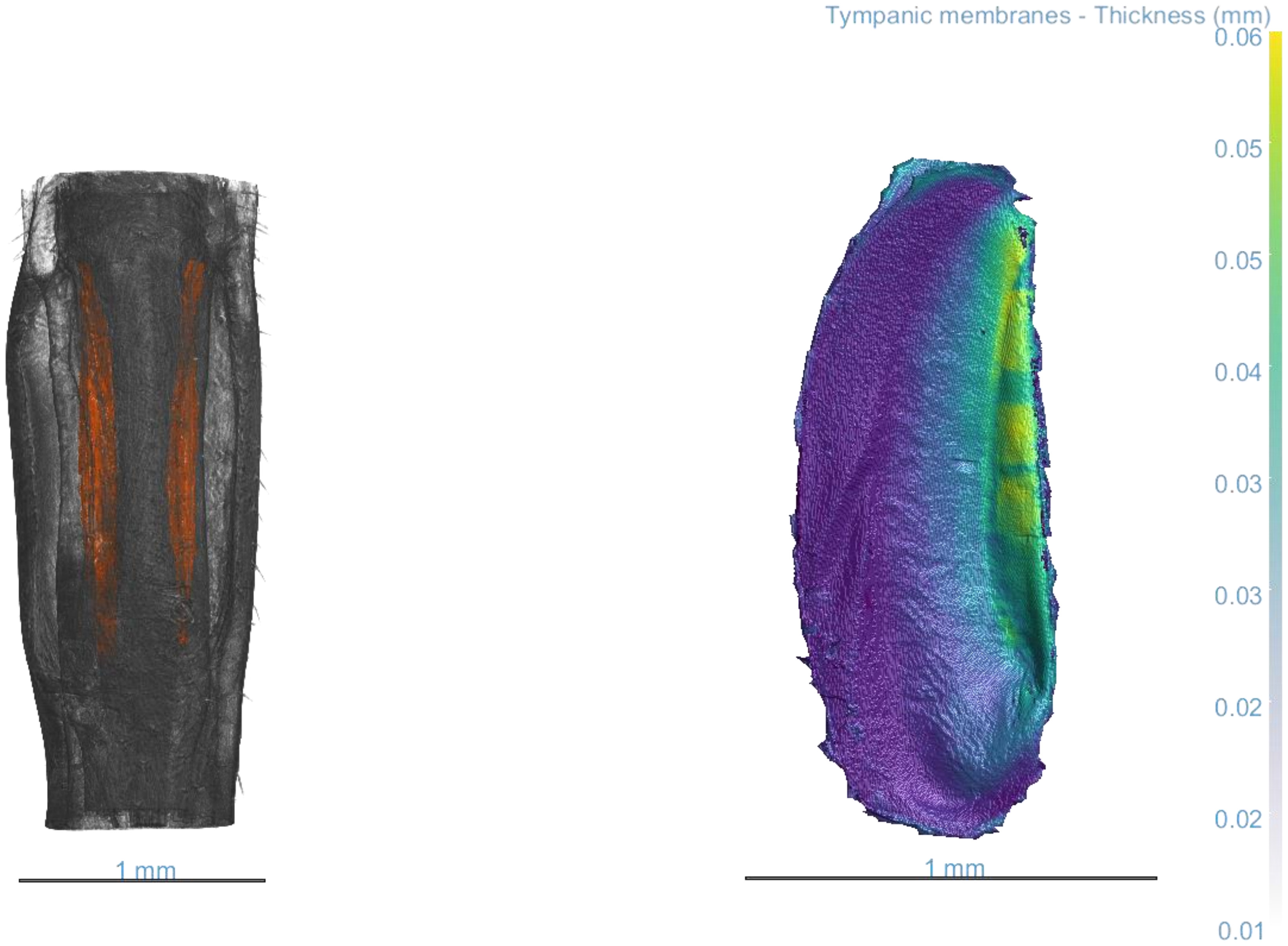

*Stilpnochlora sp.*

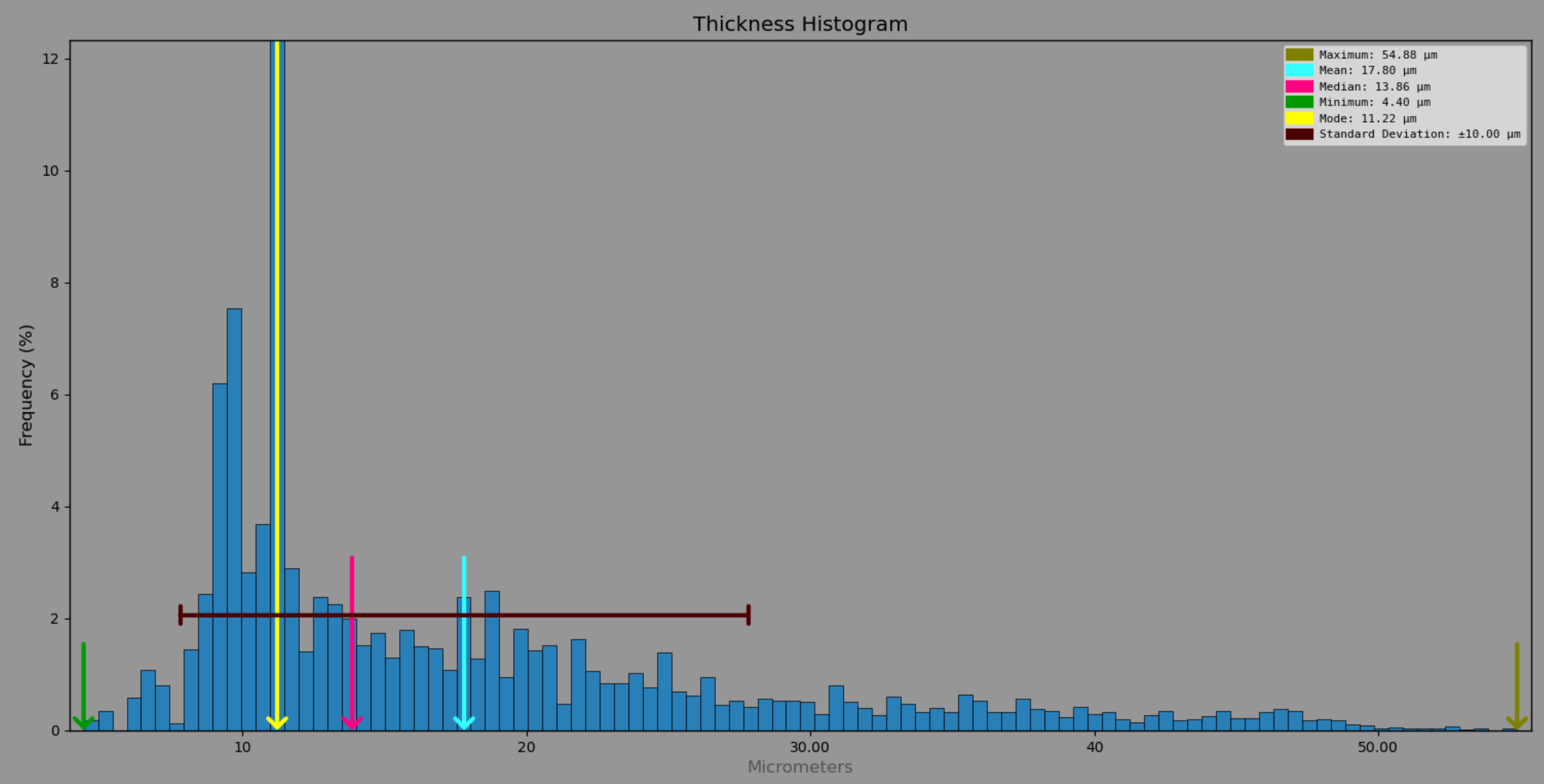
